# Nicotine biosynthesis completed by cryptic activating glucosylation

**DOI:** 10.64898/2025.12.04.692101

**Authors:** Benjamin T. W. Schwabe, Isabelle M. Angstman, Katharina Vollheyde, Zoe Ingold, Jiacheng Li, Ksenia S. Stankevich, Christopher D. Spicer, Martin A. Fascione, Gideon Grogan, Fernando Geu-Flores, Benjamin R. Lichman

## Abstract

Nicotine is a neuroactive alkaloid produced by tobacco (*Nicotiana tabacum*) as a defense against herbivory, and an addictive stimulant that has been used by humans for millennia. Despite its significance, the core steps of its biosynthesis have remained elusive. Here, we demonstrate *in vitro* reconstruction of nicotine synthase, a four-enzyme stereoselective biocatalytic cascade that forms (*S*)-nicotine from nicotinic acid and *N*-methylpyrrolinium. This cascade includes two glucose-processing enzymes that participate in a cryptic activating glucosylation step. We also reconstruct this pathway *in planta* and present high resolution X-ray structures of the key oxidoreductases A622 and BBL bound to their substrate and product, respectively. This work establishes the complete biosynthetic pathway to nicotine, providing new gene targets for controlling alkaloid production in *Nicotiana* and unlocking enzymatic routes to pyridine alkaloids.

## Main Text

Nicotine (**1**) is a plant-derived alkaloid of profound societal and scientific importance. It is the major neuroactive and addictive component of cultivated tobacco (*Nicotiana tabacum*) products, which have been used by humans for over 10,000 years^2^ and are responsible for an ongoing global health epidemic^6^. Its neurological properties are due to its agonistic interaction with eponymous nicotinic acetylcholine receptors^7^. The alkaloid protects plants from herbivory^1^ and its insecticidal activity led to its use as an agricultural pesticide, prior to replacement by neo-nicotinoids^8^.

Tobacco and its alkaloids have formed the basis of extensive investigations into plant biology including chemical ecology^9^, metabolite transport^10^, genetic regulation^11^ and metabolic evolution^12^. Recently, tobacco species, especially *Nicotiana benthamiana*, have gained traction as chassis for the recombinant production of therapeutic molecules and proteins, including vaccines^13^. However, the presence of nicotine can cause challenges for downstream processing^14^. Despite its notability, nicotine’s biosynthesis has not been elucidated: solving it will enable genetic manipulation of the pathway in native and heterologous producers, and will provide biocatalytic tools for the formation of pyridine-containing compounds.

Alkaloid scaffolds are typically formed via Mannich-like reactions, wherein an iminium electrophile is attacked by a carbon nucleophile, forming a carbon-carbon bond and a chiral center^15^. Nicotine (**1**) consists of pyridine and *N*-methylpyrrolidine rings, derived from nicotinic acid (**2**) and *N*-methylpyrrolinium (**3**) respectively (Fig. 1)^16,17^. Labelled precursor feeding studies indicate that that **2** undergoes C6-reduction and decarboxylation to yield the unusual nucleophile 1,2-dihydropyridine (**4a**), which reacts at C3 with the electrophilic iminium **3** to form 3,6-dihydronicotine (**5a**) followed by stereoselective removal of the C6-hydrogen to form **1** (Fig. 1, red hydrogen)^3,18–24^. The formation of **1** appears to be stereoselective, as the (*S*)-**1** enantiomer makes up 96% of **1** formed in *N. tabacum* roots, prior to its enrichment in leaves through enantioselective demethylation of (*R*)-**1**^25^. Related pyridine alkaloids anabasine (**6**) and anatabine (**7**) are formed when **3** is replaced by lysine-derived Δ^1^-piperideinium (**8**) or nicotinic acid-derived 2,5-dihydropyridinium (**9**) respectively (Fig. 1)^26^.

**Fig. 1.**
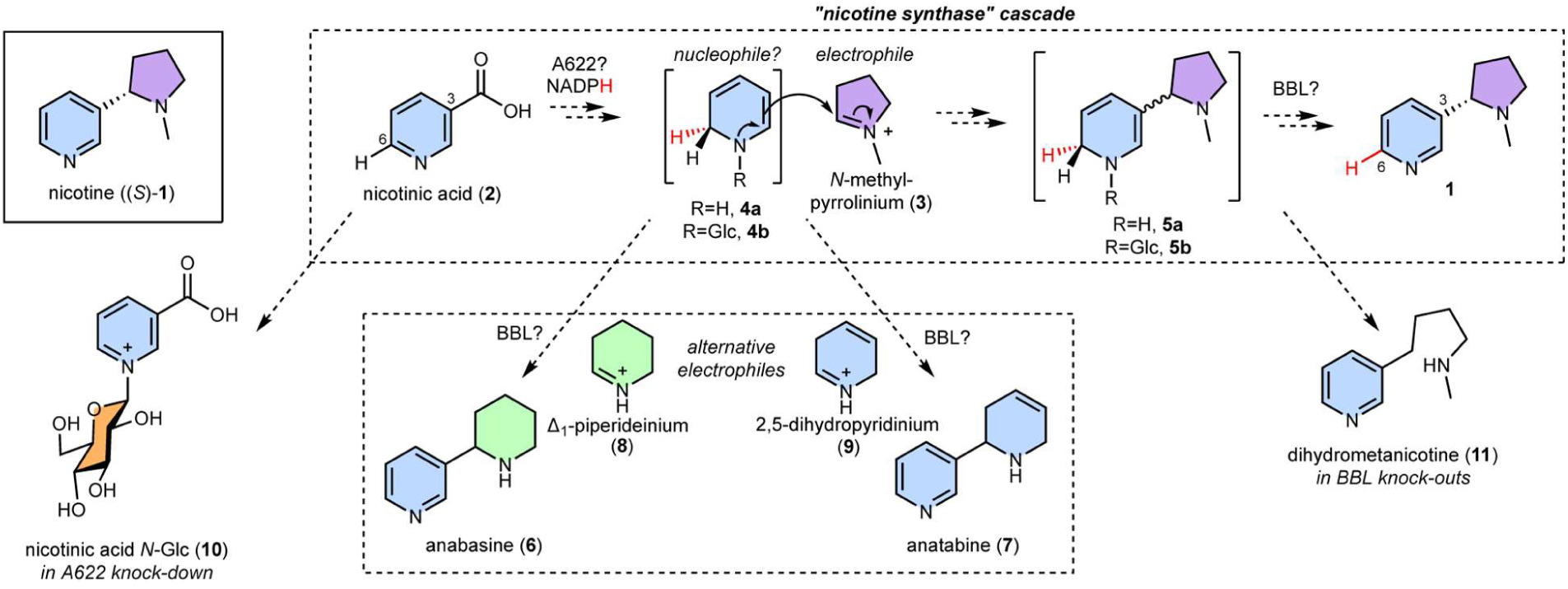
Proposed biosynthesis of nicotine. Hypothesized biosynthetic route to (*S*)-nicotine (**1**), showing origins of the pyridine (blue) and *N*-methylpyrrolidine (purple) rings. The Mannich-like scaffold formation is proposed to proceed via a multi-step “nicotine synthase” cascade. Compounds that accumulate in *Nicotiana* silencing/knock-out experiments are depicted. The red hydrogens follow the fate of the hydride transferred via enzymatic reduction of **2**.

In 1990, the isolation of “nicotine synthase” was reported, an *N. tabacum* crude enzyme preparation capable of forming (*S*)-**1** from **2** and **3**^3^. Subsequently, A622 and BBL were described, two oxidoreductases involved in nicotine scaffold formation first identified in low-nicotine tobacco mutants^4,5,27–31^. However, their substrates and products have not been determined. Silencing of *A622* and its close homolog *A622L* in hairy root cultures led to a reduction in pyridine alkaloids and an increase in nicotinic acid *N*-glucoside (**10**) (Fig. 1)^4^, suggesting that A622, an isoflavone reductase-like enzyme, is responsible for the reduction of **2** or a derivative. However, recombinant enzyme assays failed to demonstrate activity of A622 with **2** or **10**^4^. *BBLs* encode vacuolar-localized flavin-containing oxidases from the berberine bridge enzyme family^32^. RNAi-based suppression or CRISPR-based inactivation of *BBLs* led to a decrease in alkaloid content and an accumulation of dihydrometanicotine (**11**), which may be derived from dihydronicotine (**5a**) (Fig. 1)^5,31^. Furthermore, residual **1** in quintuple *BBL* knockouts was racemic and displayed a labelling pattern consistent with disruption of stereoselective H-loss at C6^31^. Single gene knockouts suggest that *BBLa* and *BBLb* are responsible for (*S*)-**1** formation whereas *BBLc* may be responsible for (*R*)-**1** formation^33^. Whilst it appears that BBL paralogs have roles in both oxidation and stereochemical control, the accumulation of **11** in knockouts suggests they are not responsible for the ring coupling step.

It is nearing two hundred years since the first isolation of nicotine, in 1828, by Posselt and Reimann^34^. Since then, the nicotine biosynthesis pathway has been under intense scrutiny due to its commercial and scientific significance. Nevertheless, progress in defining the crucial step in its formation—the coupling of the two rings—has been remarkably static, with current conceptions remaining essentially unchanged since those proposed in the 1960s^18^. In this work we resolve this enduring mystery, revealing the molecular and enzymatic basis for nicotine biosynthesis.

## Results

### Proposal of glucosylated intermediates in nicotine biosynthesis

Genes involved in plant specialized metabolism pathways can be arranged in biosynthetic gene clusters, where non-homologous but functionally related genes are found in close genomic proximity^35^. We investigated the genomic context of nicotine biosynthesis genes (Table S1), and identified a cluster consisting of *A622* (*884g0010* from Edwards *et al*^36^), the nicotine-related tonoplastic *MATE1* transporter (*884g0030*)^4,37,38^ and a previously unreported gene predicted to encode a vacuolar localized *β*-glucosidase (*β-GD1*, *884g0020*) (Extended Data Fig. 1A, Fig. S1). We identified the homeologous cluster on chromosome 12, featuring *A622L* (*31g0370*), *MATE2* (*31g0420*) and a *β-GD* paralog (*β-GD2, 31g0400*) (Extended Data Fig. 1A). All clustered genes, as well as an unclustered *β*-GD paralog (*β-GD3, 795g0060*) (Table S1), had highest expression in roots and co-expressed tightly with known nicotine biosynthesis genes (Extended Data Fig. 1B and C, Extended Data Table 1)^39^.

The clustering and co-expression pattern of the *β-GD*s suggested they are involved in nicotine biosynthesis, implying on-pathway deglucosylation. This led us to reconsider nicotinic acid *N*-glucoside (**10**) as the substrate of A622, hypothesizing that 1,2-dihydropyridine glucoside (**4b,** R=Glc) and 1,2-dihydronicotine glucoside (**5b**, R=Glc) may be pathway intermediates (Fig. 1). Therefore, the remaining step requiring a candidate gene was the glucosylation of nicotinic acid (**2**). We identified a UDP-dependent glycosyltransferase (UGT) candidate (*UGT1*, *6222g0020*) whose expression mirrored *A622* expression after topping, a process that elicits nicotine biosynthesis (Extended Data Fig. 1D)^40^. *UGT1* expression closely correlates with nicotine biosynthesis genes across multiple tissues (Extended Data Fig. 1B and C, Extended Data Table 2). Having identified *β-GD* and *UGT* candidates and proposed functions for A622 and BBL, we set out to test our hypothesis through pathway reconstruction.

### *In vitro* reconstitution of nicotine biosynthesis

We first reconstructed the nicotine biosynthesis pathway *in vitro*, selecting this simple system as it lacks interfering enzymes, non-relevant metabolites and complications of subcellular and tissue localization. Four heterologously expressed and purified enzymes (UGT1, A622, BBLa and *β*-GD1, Table S2, Fig. S2) were combined in one-pot reactions alongside co-substrates (UDP-glucose, NADPH) and precursors (Fig. 2, Extended Data Fig. 2, Fig. S3-S7, Table S3). In the presence of all four enzymes, (*S*)-nicotine ((*S*)**-1**) was formed from nicotinic acid (**2**) and *N*-methylpyrrolinium (**3**), reconstituting “nicotine synthase” activity (Fig. 2A-C, Extended Data Fig. 2, Fig. S3)^3^.

**Fig. 2.**
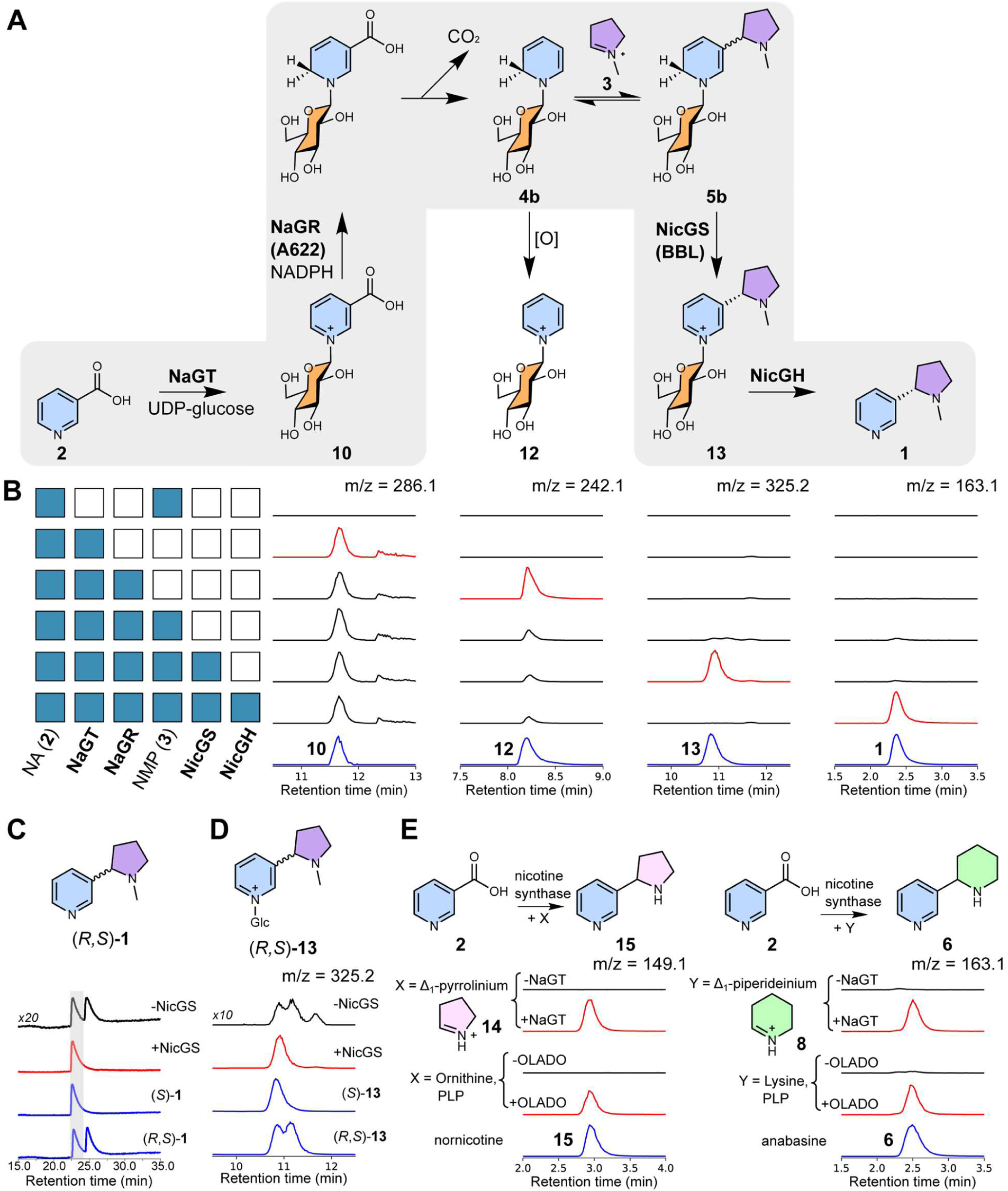
*In vitro* reconstitution of nicotine biosynthesis. (**A**) Proposed chemical transformations in nicotine biosynthesis reconstitution. (**B**) Products of *in vitro* cascades. Each row corresponds to an *in vitro* reaction, with the matrix showing presence/absence (blue/white) of reaction components, alongside UDP-Glc and NADPH. Each column shows extracted ion chromatograms (EICs, *m/z*±0.15). Blue chromatograms are chemically verified standards, scaled for clarity. Red chromatograms highlight key products. Quantification is available in Extended Data Fig. 2 and Table S3, MS^2^ data is available in Fig. S3. (**C**) Stereoselective formation of (*S*)-nicotine (**1**). Nicotine formation was achieved through one-pot enzymatic cascade (**2**, **3**, NaGT, NaGR, NicGH) with and without NicGS detected by chiral HPLC (260 nm). Signal intensity of peaks and standards (blue) were scaled for comparison. (**D**) Induction of stereoselectivity by NicGS. EIC (*m/z* 325.2±0.15) showing (*R*,*S*)-nicotine glucoside (**13**) formation from **2** with NaGT, NaGR and NMP, with or without NicGS. Chemically verified standards are blue. Signal intensity was scaled for comparison. (**E**) Formation of nornicotine (**15**) and anabasine (**6**) via the nicotine synthase cascade (**2,** NaGT, NaGR, NicGS, NicGH) with **14** or **8** respectively either synthesized or formed enzymatically by OLADO.

The UGT1 enzyme was determined to be a nicotinic acid *N*-glucosyltransferase (NaGT), producing nicotinic acid *N*-glucoside (**10**) from nicotinic acid (**2**) and UDP-glucose (Fig. 2B, Extended Data Fig. 2, Fig. S4). A622 is nicotinic acid *N*-glucoside reductase (NaGR), catalyzing the NADPH dependent reduction of **10** (Fig. 2B, Extended Data Fig. 2, Fig. S5, Fig. S6A), with the inferred product 1,2-dihydropyridine glucoside (**4b**) undergoing non-enzymatic oxidation to yield pyridine glucoside (**12**). When *N*-methylpyrrolinium (**3**) was included, the levels of **12** decreased, and a diastereomeric mixture of (*R*,*S*)-nicotine glucoside ((*R,S*)-**13**) appeared (Extended Data Fig. 2, Fig. 2D), alongside traces of dihydrometanicotine **11** and dihydrometanicotine *N*-glucoside (Glc-**11**) (Fig. S5). This is likely due to a non-stereoselective Mannich reaction between **3** and **4b**, forming (*R*,*S*)-dihydronicotine glucoside (**5b**) which undergoes non-enzymatic oxidation to yield (*R,S*)-**13**, or decomposition into Glc-**11** and **11**. Inclusion of BBLa led to a significant decrease in **12**, increase in **13** and rendered the reaction diastereoselective, with the (*S*)-**13** isomer in excess (Fig. Extended Data Fig. 2, Fig. 2D), reflected in the exclusive production of the (*S*)-**1** enantiomer in the full pathway reconstruction (Fig. 2C). In the cascades, the peaks corresponding to **11** and Glc-**11** are absent in the presence of BBLa; yet Glc-**11** is not a substrate of the enzyme (Fig. S6B), implying BBLa’s substrate is upstream. These results redefine BBLa as (*S*)-nicotine glucoside synthase (NicGS). The final step of the pathway is the deglucosylation of (*S*)-(**13**) to form (*S*)**-1**, which can be catalyzed by *β*-GD1, now defined as nicotine glucoside hydrolase (NicGH) (Fig. 2, Extended Data Fig. 2, Fig. S7).

### *In vitro* biosynthesis of alternative tobacco alkaloids

We anticipated that we could exchange *N*-methylpyrrolinium (**3**) for different electrophiles to form related pyridine alkaloids. We demonstrated this with Δ^1^-pyrrolinium (**14**) and Δ^1^-piperideinium (**8**), to form the natural products nornicotine (**15**) and anabasine (**6**) respectively (Fig. 2E). We generated the electrophiles via two approaches, chemically and via *in situ* oxidative decarboxylation from ornithine and lysine respectively, catalyzed by a recently discovered ornithine/lysine/arginine decarboxylase-oxidase (OLADO) from *N. tabacum*^41^.

### *In vitro* assays with isotopically labelled nicotinic acid

Feeding experiments with isotopically labelled precursors have been crucial in establishing atomic level information about nicotine biosynthesis, such as the loss of the C6 hydrogen from nicotinic acid^19–21,31^. We reconstructed nicotine synthase using nicotinic acid-(*ring*-d_4_) (**2-**d_4_) or nicotinic acid-d^6^ (**2-**d^6^) as substrates and observed the products **1**-d_3_ or **1**-d_0_ respectively (Fig. 3A-B, Extended Data Fig. 3). This labelling is consistent with NaGR (A622) catalyzed reduction of **2** and the NicGS (BBLa) catalyzed oxidation of 1,2-dihydronicotine glucoside (**5b**) both occurring at C6, with a protide introduced by A622 and the deuteride abstracted by BBLa. In contrast, non-enzymatic oxidation favors protide abstraction due to the kinetic isotope effect, as seen in the abundance of more highly deuterated products (e.g. **1**-d_4_ and **1**-d^6^ from **2-**d_4_ and **2-**d^6^ respectively) in the absence of NicGS (BBLa) (Fig. 3A-B, Table S4), and in the predominance of **12**-d_4_ in the **2**-d_4_ fed pathway (Extended Data Fig. 3). The inclusion of NicGS (BBLa) significantly increases the ratio of **12**-d_3_ to **12**-d_4_ which implies that it can bind and catalyze the oxidation **4b** as well as **5b** (Extended Data Fig. 32, Table S4). Cascade formation of nornicotine-d_3_ (**15**-d_3_) and anabasine-d_3_ (**6**-d_3_) from **2**-d_4_ confirmed they are also formed via NicGS (BBLa) catalyzed oxidation (Fig. 3C and D).

**Fig. 3.**
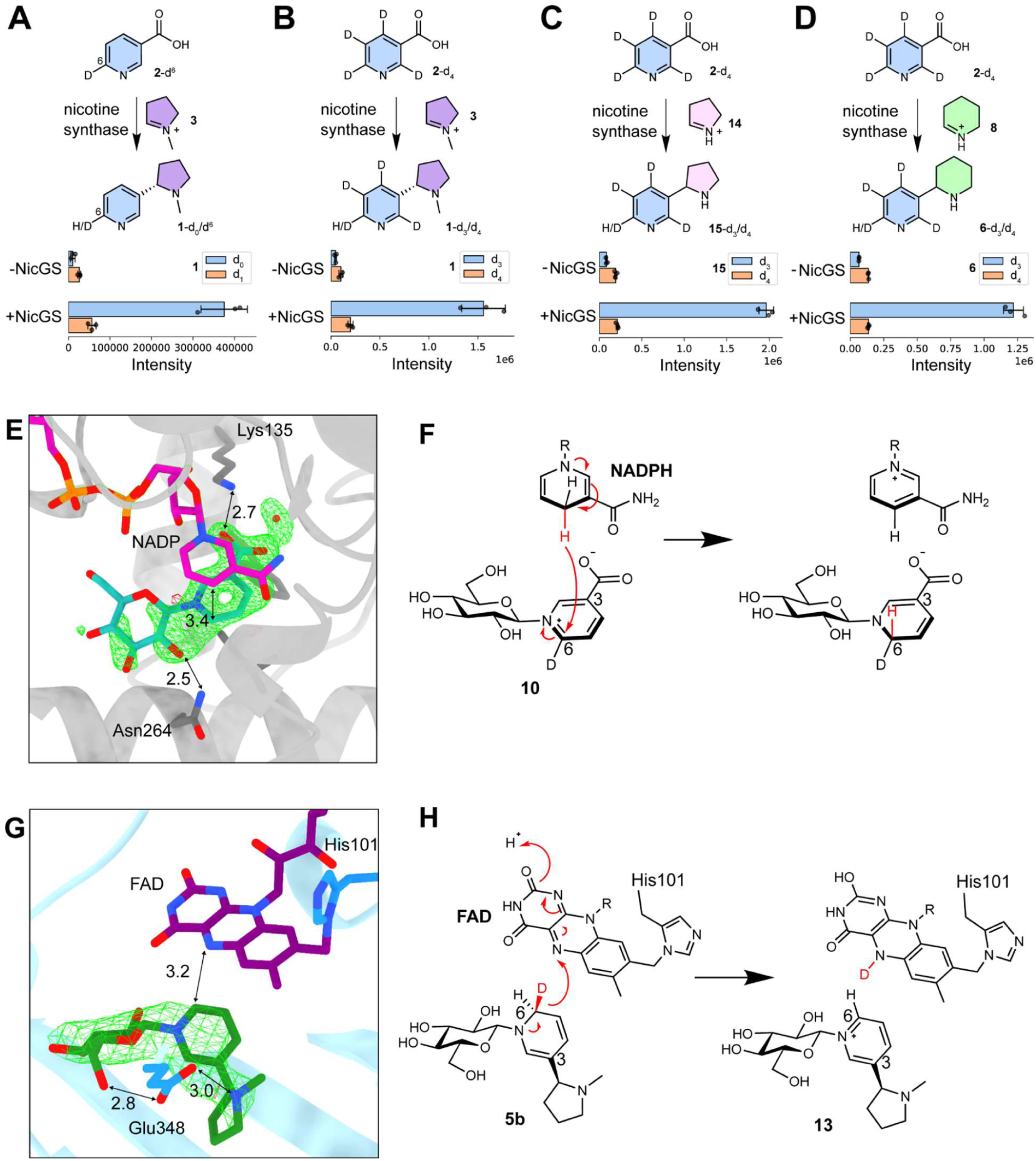
Resolving the mechanism of NaGR (A622) and NicGS (BBLa). (**A-D**) Conversion of nicotinic acid (**2**) isotopologues by nicotine synthase (NaGT, NaGR, NicGH, NADPH and UDP-Glc) with and without NicGS. Bar charts show peak areas of products from EICs. Reactions were performed in triplicate (points); bars show mean peak area; error bars show standard deviation. (**E**) The crystal structure of NaGR bound to NADP^+^ and nicotinic acid *N*-glucoside (green net shows unrefined ligand density Fo-Fc at 3 σ). (**F**) Proposed mechanism of NaGR: NADPH reduces nicotinic acid *N*-glucoside-d^6^ on the *re*-face adding a hydride in the pro-*R* position. (**G**) The crystal structure of NicGS with covalently bound FAD and ligand (*S*)-nicotine glucoside (green net shows unrefined ligand density Fo-Fc at 3 σ). (**H**) Proposed mechanism of NicGS, abstracting the pro-*S* hydride from dihydronicotine glucoside-d^6^ to yield nicotine glucoside.

### Structure of NaGR (A622)

To shed light on the mechanisms of NaGR (A622) and NicGS (BBLa), we determined their structures in complex with co-factors and relevant ligands using X-ray crystallography (Table S5). In the case of NaGR (A622), we obtained a 1.3 Å resolution crystal structure in complex with NADP^+^ and nicotinic acid *N*-glucoside (**10**) (9RDD) (Extended Data Fig. 4). NaGR (A622) is most similar to dehydrogenases of the isoflavone reductase family (*e.g.* 2GAS, 57% sequence identity)^42^, with the cofactor NADP^+^ bound between the Rossmann fold and the substrate binding domain. Omit density corresponding to the ligand nicotinic acid *N*-glucoside **10** was observed adjacent to the cofactor, with the C6 atom of the pyridine ring poised to receive hydride from C4 of the nicotinamide ring of the cofactor at a distance of 3.4 Å (Fig. 3E). This matches the regio-and stereo-selectivity of the hydride transfer from NADPH onto the *re*-face of **10** at C-6, and into the pro-*R* position (Fig. 3F). The carboxylate is bound by the side chain of Lys135. The sugar density was not complete, but an interaction was observed between the side chain of Asn264 and the C2 hydroxyl of the sugar.

### Structure of NicGS (BBLa)

We obtained 2.5 Å resolution structures of NicGS (BBLa) in complex with FAD (9RUW), and with both FAD and the putative reaction product (*S*)-nicotine glucoside (**13**) (9RDR) (Table S5, Fig. 3G, Extended Data Fig. 4). NicGS is structurally most similar to the reticuline oxidase from *Eschscholzia californica* (3D2D)^43^, but the FAD is not bi-covalently bound as in the case of 3D2D, with C166 replaced by G165 in NicGS. Omit density was observed adjacent to the FAD density that was successfully modelled as (*S*)-**13** (Fig. 3G). The C-6 position of **13** is 3.2 Å from the reactive nitrogen of the FAD cofactor, with the *si*-face of the pyridine ring facing the FAD. This proximity indicates that the C6 pro-*S* hydride of the proposed dihydronicotine glucoside (**5b**) substrate reduces the FAD cofactor (Fig. 3H). The side chain of E348, not conserved in 3D2D, has a role in substrate recognition, interacting with the nitrogen of the pyrrolidine ring and also the exocyclic C6 hydroxyl of **13** at distances of 3.0 and 2.8 Å respectively. In the absence of ligand binding (9RUW) the loop containing this residue is unstructured (Extended Data Fig. 4). Overall, the structures of NaGR (A622) and NicGS (BBLa), with bound glucoside substrates or products lend strong support to our pathway proposal. Furthermore, the atomic details provide a structural/mechanistic validation of isotope labelling experiments, where NicGS (BBLa) abstracts the opposite hydride to that added by NaGR (A622).

### *In planta* pathway reconstitution

To demonstrate that the pathway assembled *in vitro* could be operational *in planta*, we reconstituted nicotine biosynthesis in leaves of *N. benthamiana* via transient gene expression (Fig. 4, Table S6)^44^. Whilst *N. benthamiana* is capable of producing nicotine, we were able to independently study the reconstituted pathway as the key endogenous genes *A622* and *BBL* are not expressed in leaves (Extended Data Fig. 5A, Extended Data Table 3)^45^. Furthermore, we could distinguish between the native nicotine transported into the leaves and the nicotine produced *in situ* by feeding the engineered leaves with a labelled precursor (nicotinamide-d_4_) and analyzing labelled nicotine.

**Fig. 4.**
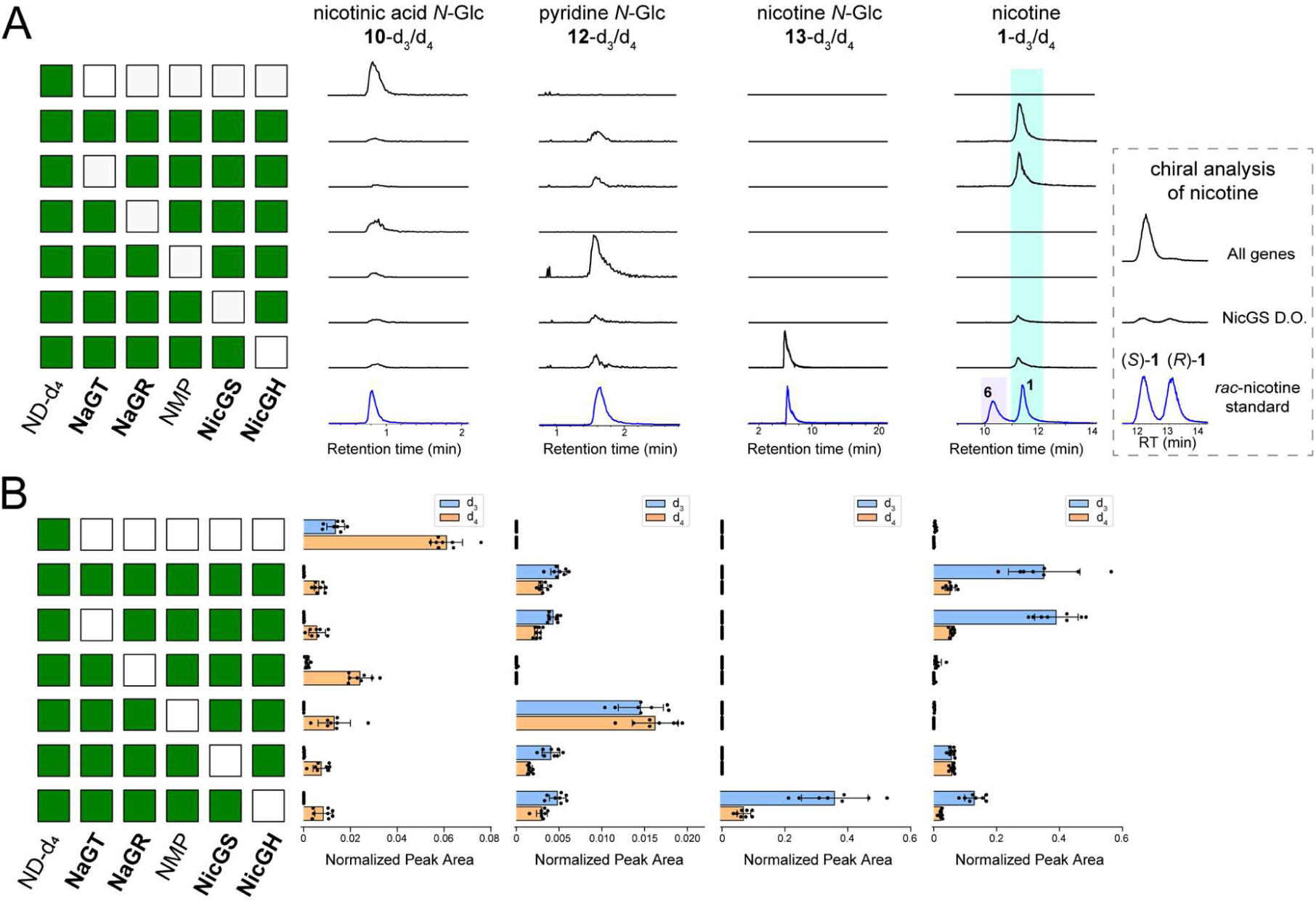
*In planta* reconstitution of nicotine biosynthesis. Transient expression of nicotine synthase genes together with labelled precursor feeding (nicotinamide-d_4_ (ND-d_4_)) in the leaves of *N. benthamiana*. The gene combination producing the *N-*methylpyrrolinium (ODC, PMT and MPO) is abbreviated as NMP. (**A**) Products of *in planta* expression and feeding. Each row corresponds to a gene combination, with the matrix showing presence/absence (green/white) of co-infiltrated genes. Columns show EICs (*m/z*±0.005) corresponding to the labeled compounds in comparison to chemically synthesized or commercial standards (blue) (see Fig. S7 for MS^2^ spectra). The nicotine and anabasine standards were analyzed as a mixed sample. The insert shows the chiral analysis of nicotine in the presence of all genes and in the NicGS drop-out. Signal intensities are normalized to the largest peak in the column; standards were re-scaled to match the largest peak. (**B**) Individual quantification of d_3_ and d_4_ isotopologues corresponding to the compounds and gene combinations shown in panel (A). Bars show average EIC peak area (*n* = 8); points are individual samples; error bars show standard deviation (see statistics on Table S7).

To produce *N*-methylpyrrolinium (**3**), we co-expressed the known tobacco enzymes ODC, PMT and MPO (Table S1 and S6)^46–48^. When the entire pathway was expressed, we observed substantial production of (*S*)-nicotine-d_3_ ((*S*)-**1**-d_3_) (Fig. 4, Fig. S8), validating our *in vitro* observations. Additionally, stepwise reconstruction of the pathway mirrored results from the *in vitro* stepwise reconstruction, including production of anabasine-d_3_ (**6-**d_3_) enabled by co-expression of OLADO in place of the enzymes forming **3** (Extended Data Fig. 6, Table S7).

We then conducted a drop-out experiment to validate the role of the individual components (Fig. 4, Table S8). Drop-out of NicGH (*β*-GD1) led to a decrease in labelled nicotine (**1**) and accumulation of labelled nicotine glucoside (**13**), which we did not detect in any other condition. Dropping out NicGS (BBLa/b), resulted in a similar reduction in labelled nicotine but the residual nicotine was racemic (Fig. 4A). Omission of the *N-*methylpyrrolinium forming enzymes eliminated nicotine production and led to the accumulation of labelled pyridine glucoside (**12**). Dropping out NaGR (A622) abolished production of labelled pyridine glucoside (**12**), blocking the pathway at nicotinic acid *N-*glucoside (**10**). It was not possible to confirm the *in planta* activity of NaGT (UGT1), as control leaves fed with labelled precursor accumulated labelled **10,** likely due to high leaf expression of the native *UGT1* homolog (Extended Data Fig. 5B). Similar to the *in vitro* experiments, the *in planta* experiments were able to ascribe the loss of deuterium to NicGS (BBLa/b) (Fig. 4, Extended Data Fig. 6, Table S7 and S8).

### Metabolites in wildtype and mutant plants

We further probed our pathway hypothesis by searching for proposed intermediates or shunt metabolites in wildtype roots of *N. tabacum* and *N. benthamiana.* We were able to detect nicotine glucoside (**13**) in *N. tabacum* and pyridine glucoside (**12**) in both *N. tabacum* and *N. benthamiana* (Fig. 5, Fig. S9-S10). We also examined roots of existing *N. benthamiana* knockout lines in NaGR (*A622/A622L*)^30^ and NicGS (*BBLa/b/c/d/d’*)^31^. The NaGR (A622) knockout accumulated nicotinic acid *N*-glucoside (**10**) but not pyridine glucoside (**12**), supporting the identity of **12** as a shunt product of NaGR (A622) (Fig. 5A). The NicGS (BBL) knockout accumulated pyridine glucoside (**12**) and nicotine glucoside (**10)**, similar to wildtype *N. tabacum* roots (Fig. 5A).

**Fig. 5.**
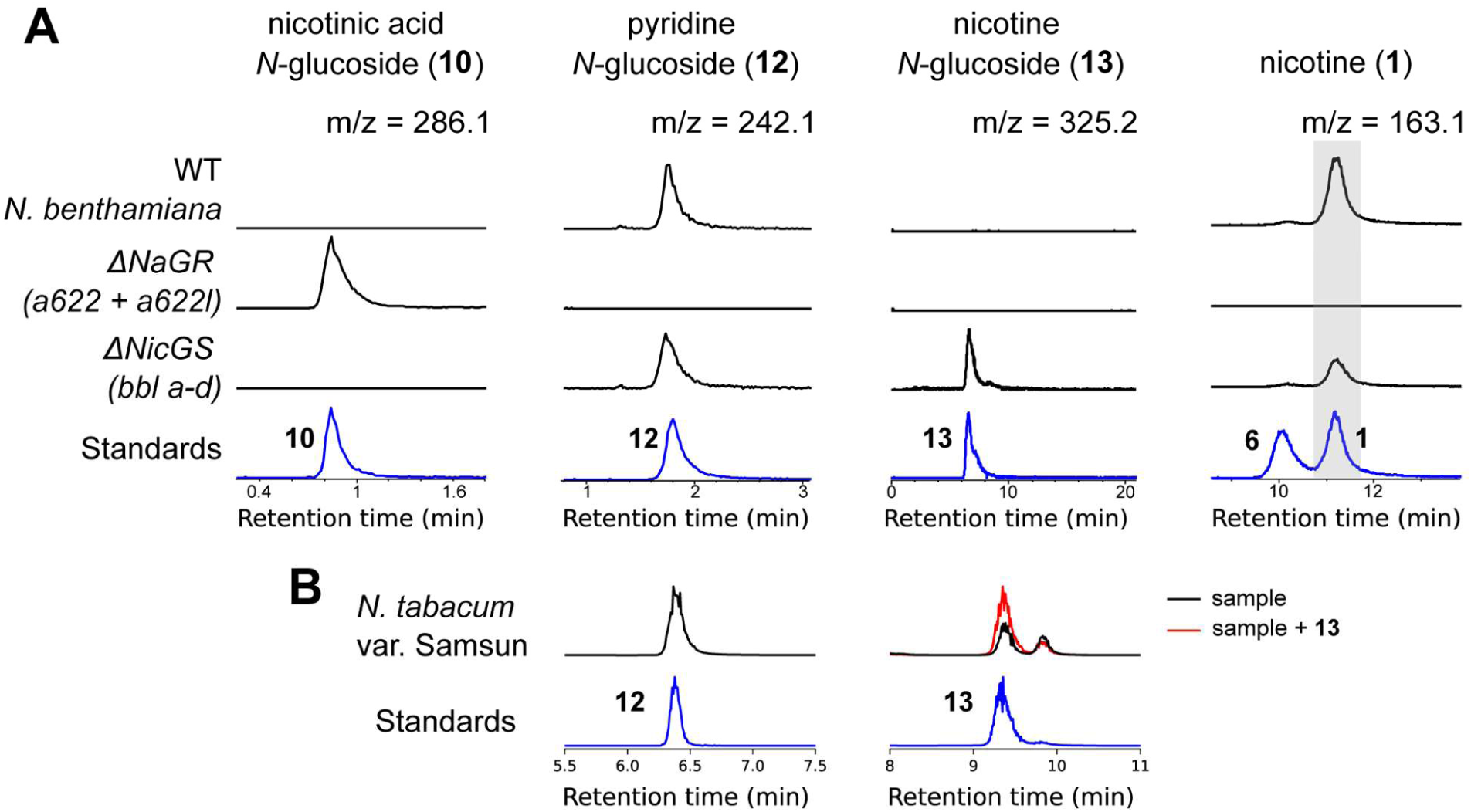
Identification of pathway intermediates and shunt products in *Nicotiana* roots. (**A**) Metabolite analysis of roots from *N. benthamiana* wild type, *ΔNaGR*^30^ and *ΔNicGS*^31^ lines (black) in comparison to standards (blue) (see MS^2^ verification in Fig. S9). Signal intensities represent EICs (*m/z*±0.005) and were normalized to the largest peak in each column; standards were re-scaled to match the largest peak. (**B**) Metabolite analysis of roots from *N. tabacum* var Samsun (black), compared to standards (blue) (see MS^2^ verification in Fig. S10). Red trace is root extract sample spiked with 10 μM nicotine *N*-glucoside (**13**) to verify peak identity. Signal intensities are EICs (*m/z*±005); standards were re-scaled to match the largest peak in each plot.

## Discussion

It is remarkable that, despite the massive global use of tobacco, its economic importance, and the volume of investigation into nicotine biosynthesis, key steps in its formation have remained enigmatic. Here, we have reconstituted, *in vitro* and *in planta*, a minimal nicotine synthase cascade, forming (*S*)-**1** from nicotinic acid (**2**) and *N*-methylpyrrolinium (**3**). It consists of four enzymes: NaGT, NaGR, NicGS, and NicGH. Cryptic biosynthetic steps cannot be predicted based on the structures of precursors or products, making them a bottleneck in pathway elucidation. In nicotine biosynthesis, NaGT and NicGH are responsible for cryptic glucosylation, which acts like an activating group, increasing the electrophilicity of nicotinic acid (**2**) making it susceptible to reduction by NADPH.

The substrate for NaGR (A622) is nicotinic acid *N*-glucoside (**10**), which is consistent with its accumulation when *A622* is silenced or knocked-out^4^. Previous attempts to characterize this reaction through spectrophotometry, typical for NADPH dependent reactions, may have been hampered by overlapping absorbance spectra in the NADPH substrate and the 1,2-dihydropyridine glucoside (**4b**) product. Furthermore, **4b** appears unstable *in vitro*: here its presence is inferred through the identification of pyridine glucoside (**12**), or through the reaction outcome in the presence of an electrophile.

The cascade operates stereoselectively, with formation of (*S*)-enantiomer of **1** dominating both *in vitro* and *in planta* (Fig. 2 and 4). NicGS (BBLa) is responsible for determining the stereoselectivity of the pathway, as we previously showed *in planta*^31^. The crystal structure of NicGS bound to (*S*)-nicotine glucoside ((*S*)-**13**) indicates it catalyzes the diastereoselective oxidation of dihydronicotine glucoside (**5b**), but it is unclear whether it also catalyzes the stereoselective Mannich reaction and/or a dynamic kinetic resolution process. The accumulation of the shunt product pyridine glucoside **12** is inhibited in the presence of **3** even in the absence of NicGS, indicating facile non-enzymatic coupling between **4b** and **3**. However, the decreased concentration of **12,** higher conversion into **13** and induction of stereoselectivity in the presence of NicGS suggests enzymatic control of this step, as does the indication that NicGS can catalyze oxidation of **4b**, which would match stepwise binding of **4b** and **3**. Further work is required to resolve the intricacies of the NicGS mechanism given the complexity of the enzyme cascade (Extended Data Fig. 7).

The modularity of the cascade with respect to the electrophile has been demonstrated by the formation of anabasine and nornicotine, and suggests the system could form the basis of a biocatalytic route towards diverse functionalized pyridines. New genetic understanding of nicotine biosynthesis also has potential to inform engineering of *Nicotiana* sp., including the elimination or rerouting of alkaloid biosynthesis for new synthetic biology chassis. This work resolves a centuries old puzzle in chemistry, plant biology and biochemistry, and will lead to the development of novel biotechnological and biocatalytic tools to produce valuable chiral chemicals.

## Methods

### Expression analysis

Genes of interest were defined by assessment of the literature related to nicotine biosynthesis (Table S1). Protein sequences were obtained via GenBank or UniProt accession numbers, and used as tblastn queries against the coding sequences from the 2017 *N. tabacum* genome^36^ to obtain gene names and sequences. Biosynthetic gene clusters were identified using functional annotations and RNA-seq profiles of genes neighboring genes of interest, using JBrowse hosted on the SolGenomics^49^, querying the 2017 *N. tabacum* genome^36^. Gene names were used as queries for the Plant Gene Expression Omnibus (PEO) (https://peo.ku.dk/)^39^. The PEO *N. tabacum* dataset consists of TPMs for 824 samples over twelve tissue types. Precomputed Pearson Correlation Coefficient (PCC) values were used to identify top-correlating genes.

For gene co-expression analysis, TPMs of genes of interest were extracted from PEO and analysed in R-Studio (tidyverse and pheatmap packages). For plotting gene expression from *N. tabacum* across tissues, log_2_(TPM+1) values were grouped by Plant Ontology (i.e. tissue type), averaged, and plotted as a heatmap, with Z-score normalisation across tissue types. For expression correlation, an all-by-all PCC matrix was computed (824 samples, log_2_(TPM+1)), depicted as a heatmap, with genes grouped with complete-linkage hierarchical clustering on Euclidean distance. *Nicotiana benthamiana* nicotine biosynthesis genes homologs were identified by top blastn hit using *N. tabacum* genes as queries of the *N. benthamiana* v1.0.1 genome^49^. Gene names were used as queries for PEO and TPM values were extracted (476 samples), grouped by Plant Ontology and plotted as boxplots (log_2_(TPM+1))^39^.

For assessment of gene expressions after topping, we extracted a list of differentially expressed genes (DEGs) in tobacco roots after topping, from Qin *et al* 2020^40^. Using the mean FPKM value at t=0 we converted log_2_ fold change values to FPKM values across the time course and then calculated PCCs with A622 (Nitab4.5_0000884g0010) expression. Genes were filtered by annotation “UDP-glucosyltransferase” and PCC_A622_>0.85. Top hits were assessed for their overall expression level and used as a query in PEO.

### Protein purification

Codon-optimised NaGR (A622, UniProt Accession: IFRH_TOBAC), NaGT (UGT1, 2017 *N. tabacum* genome: Nitab4.5_0006222g0020^36^) were expressed using the pET28a(+) vector as C-terminal (NaGR) or N-terminal (NaGT) His-tagged proteins, using *E. coli* B834 or SoluBL21 respectively. NaGR was expressed as a Δ5 N-terminal truncate. Proteins were purified using nickel affinity chromatography followed by size-exclusion chromatography. NicGS (BBLa, 2017 *N. tabacum* genome: Nitab4.5_ 0006307g0010) and NicGH (*β*-GD1, 2017 *N. tabacum* genome: Nitab4.5_0000884g0020) were expressed as N-terminal His-tagged proteins using a *Komagataella phaffii* secretion system based on the pPICZα vector. Proteins were purified from the media using tangential flow filtration followed by nickel affinity chromatography and size-exclusion chromatography. OLADO (UniProt Accession: A0A1S3ZKS9_TOBAC) was expressed and purified as described previously^41^. Proteins were stored at −70 ℃. Details of sequences are available in Table S1 and S2), further methodological details are available in the Supplementary Information.

### In vitro assays

*In vitro* enzyme assays and multi-enzyme cascades were carried out in 50 μL reactions, in 50 mM phosphate buffer pH 7.4, 100 mM NaCl. Reactions contained 1 mM nicotinic acid, 1 mM *N*-methylpyrrolinium, 0.5 mg mL^-1^ NaGT (UGT), 0.5 mg mL^-1^ NaGR (A622), 50 μg mL^-1^ NicGS (BBLa) and 5 μg mL^-1^ NicGH (*β*-GD). Other combinations were tested by removing component(s) of this mix to make different combinations of enzymes/substrates. The reactions were incubated for 16 hours at 37 ℃, then quenched (200 μL, 90:10 acetonitrile:water, 20 mM ammonium formate pH 3), centrifuged (21,000 xg, 10 min) and the supernatant was transferred to vials for LC-MS analysis. The *N*-methylpyrrolinium was prepared by incubation of 1 μL γ-methylaminobutyraldehyde diethyl acetal with 49 μL 1 M HCl at 70 ℃ for 1 hour before neutralisation with 40 μL 1 M NaOH, incubated on ice until adding to assay on the same day. 1-pyrroline for nornicotine assays was prepared in the same way from aminobutyraldehyde diethyl acetal. Enzyme assays were analysed by LC-MS, with Hydrophilic Interaction Liquid Chromatography (HILIC) separation using a Waters XBridge BEH Amide column (5 μm, 2.1 × 100 mm) on a Thermo Scientific Vanquish UHPLC. Detection was performed on a Thermo Scientific LTQ XL Linear Ion Trap Mass Spectrometer. Details on LC-MS analytical procedures can be found in the Supplementary Information.

### Chiral nicotine analysis of *in vitro* reaction

*In vitro* enzyme assays for chiral analysis were carried out in 3 x 50 μL reactions for each condition, in 50 mM phosphate buffer pH 7.4, 100 mM NaCl. Reactions contained 5 mM nicotinic acid, 5 mM *N*-methylpyrrolinium, 10 mM UDP-Glc, 2 U Glucose-6-phosphate dehydrogenase (Merck), 5 mM NADP and 10 mM glucose-6-phosphate, with 0.5 mg mL^-1^ NaGT (UGT), 0.5 mg mL^-1^ NaGR (A622), 50 μg mL^-1^ NicGS (BBLa) and 5 μg mL^-1^ NicGH (*β*-GD). Samples for chiral HPLC analysis were basified with 1 μL 1 M NaOH, then protein precipitated with 500 μL cold EtOH. After centrifugation (21,000 xg, 10 min), the protein pellet was washed with a further 500 μL EtOH, the EtOH was then combined, evaporated to dryness and resuspended in 50 μL propan-2-ol and then loaded into vials for analysis. Assays with NicGS were diluted 10-fold further. Nicotine chiral analysis was carried out on an Agilent 1200 HPLC by normal phase isocratic elution using a CHIRALPAK® AD-H 250 x 4.6 mm column using 99:1 hexane:IPA at 1 mLmin^-1^. Nicotine peaks were detected by UV absorbance at 260 nm.

### *In vitro* assay statistics

Statistical analysis of *in vitro* reaction product formation was performed using a one-way ANOVA (R-studio, function: aov) to determine EIC peak area differences across unique reactions, followed by a Tukey’s HSD post-hoc test (function: TukeyHSD), with peak area grouping assigned letters (function: multcompLetters) based on significant differences (*p*<0.05) in pairwise comparisons. Reactions were performed in triplicate (*n* = 3). For analysis of isotopologue ratios, d_3_/d_4_ isotopologue ratios were calculated per chemical and per sample through integration and division of EIC peak areas. Samples with zero peak area for either isotopologue were removed fom the analysis. Then isotopologue ratios (*n* = 3) for each chemical were compared across unique *in vitro* reactions, analysed by analysis of variance and Tukey’s HSD as described above. To calculate the nicotine concentration of the complete cascade, a standard curve of known concentrations was run in the same batch as the samples and fitted using log-log linear regression. Model parameters were used to convert sample peak areas into concentrations. To accurately estimate the mean and standard error, we first estimated the uncertainty of individual concentrations through the regression parameter errors and residual variability, before combining these using inverse-variance weighting of the triplicates to yield a pooled mean and standard error.

### Protein crystallization

Initial screening of crystallization conditions was performed using commercially available INDEX (Hampton Research), PACT premier and CSSI/II (Molecular Dimensions) screens in 96-well sitting drop trays which were stored at 4 °C. Ligand complexes of A622 were obtained by co-crystallisation with 5 mM nicotinic acid *N*-glucoside **1**, derived from a 100 mM stock solution in H_2_O, which was added to the NaGR (A622) solution and incubated on ice for 30 min before the trays were set up. Crystals were obtained in 0.1 M succinic acid pH 7.0 and 15 % (w/v) PEG 3350. NicGS (BBLa) protein was enzymatically de-glycosylated before setting up crystallization screens. 53 μL 20 mg mL^-1^ NicGS (BBLa) was incubated overnight at 37 ℃ with 7 μL EndoH and 7 μL GlycoBuffer 3 (New England Biolabs). Initial screening of crystallization conditions was performed as for NaGR but at 16 °C. Initial *apo*-crystals of NicGS (BBLa) were obtained in drops containing 0.14 M LiSO_4_, 0.1 M Bis-Tris pH 5.5, 21% PEG 3350, by co-crystallisation with 10 mM (*S*)-nicotine glucoside **13**, derived from a 100 mM stock solution in H_2_O. Ligand bound crystals were obtained by co-crystallisation as described but with 50 mM (*S*)-nicotine glucoside. Crystals were harvested directly into liquid nitrogen with nylon CryoLoops^TM^ (Hampton Research), using the mother liquor without any further cryoprotectant (NaGR and NicGS *apo*) or by addition of 10% ethylene glycol (NicGS ligand-bound) to the drop prior to fishing.

### X-ray structure elucidation

The datasets described in this report were collected at the Diamond Light Source, Didcot, Oxfordshire, U.K. on beamline I03. Data were processed and integrated using XDS^50^ and scaled using SCALA^51^ included in the Xia2^52^ processing system. Data collection statistics are provided in Table S5. The crystals of NaGR (A622) were obtained in space group *P*1, with one molecule in the asymmetric unit; the crystals of NicGS (BBLa) were obtained in space group *P*2_1_2_1_2_1_, with two molecules in the asymmetric unit. The structures were solved by molecular replacement using MOLREP^53^ with the AlphaFold structures AF-P52579-F1-v4 and AF-F1T160-F1-v4 used as the models for NaGR and NicGS respectively (https://alphafold.ebi.ac.uk/)^54^. The structures were built and refined using iterative cycles in Coot^55^ and REFMAC^56^ employing local NCS restraints in the refinement cycles. The final structure of NaGR (A622) ligand complex exhibited % *R*_cryst_ /*R*_free_ values of 17.1/21.0; the final structures of *apo*-NicGS (BBLa) and the NicGS (BBLa) ligand complex exhibited % *R*_cryst_ /*R*_free_ values of 29.3/32.4 and 21.5/26.9 respectively. Refinement statistics for the structures are presented in Table S5. The structures of NaGR (A622)-NADP+ nicotinic acid *N*-glucoside, NicGS (BBLa)-FAD nicotine *N*-glucoside and NicGS (BBLa)-FAD have been deposited in the Protein Databank (PDB) with accession codes 9RDD, 9RDR and 9RUW respectively.

### *In planta* pathway reconstruction

The genes for transient overexpression were synthesized and cloned into a pHREAC vector (Addgene plasmid #134908)^57^ by Twist Bioscience (South San Francisco, USA). Coding sequences of the genes of interest were obtained from the *N. tabacum* v1.0 Edwards 2017 genome (scaffold)^36^ and confirmed/refined by examining mapped RNAseq data in the genome browser on the website of the Sol genomics Network^49^. A list of gene identifiers and coding sequences can be found in Table S6. The cloned sequences of all pHREAC gene inserts were checked by Sanger sequencing and the plasmids were separately transformed into the electrocompetent *Agrobacterium tumefaciens* strain AGL1. Transient expression was performed in leaves of 4 week-old greenhouse-grown wildtype *N. benthamiana* plants according to Chuang and Franke 2022^58^. pHREAC-GFP (kindly provided by Hadrien Peyret and George Lomonossoff) was used as control and as placeholder in the drop-out experiment. Four days post-infiltration, leaves were reinfiltrated with 400 µM nicotinamide-d_4_ (D4 98%; Cambridge Isotope Laboratories, Tewksbury, MA, USA) dissolved in infiltration buffer without acetosyringone. Leaf disks were harvested after an additional 2 days (6 disks/plant: 3 disks/leaf x 2 leaves per plant). The harvested samples were flash-frozen in liquid nitrogen and were stored at −70 °C until tissue homogenization using chrome balls and a TissueLyzer (Qiagen). The homogenized samples were stored at −70 °C until metabolite extraction.

### Metabolite analysis of *N. benthamiana* pathway reconstruction

Metabolites were extracted from homogenized flash-frozen plant tissue (*n* = 8) as described by Vollheyde and Dudley *et al.* 2023^31^. For the analysis, 75 ppm (386.2 µM) caffeine was used as the internal standard in the extraction solution and the proportion of tissue to extractant (µL) was 40 mg/100 µL. After the incubation step, samples were spun down for 5 min at 16200 x*g* (max speed table top centrifuge) and supernatants were diluted in ultrapure water at a proportion of 1:15. Samples were stored at −70 °C after extraction until filtering and after filtering until further analysis. The methanolic plant extracts were analysed via reversed-phase LC-MS according to Vollheyde and Dudley *et al.* 2023^31^ with an injection volume of 10 µL. Compounds were identified by comparison to commercial and synthesized standards. The chiral LC-MS analysis of methanolic plant extracts was performed according to Vollheyde and Dudley *et al.* 2023^31^ with an injection volume of 10 µL. Compounds were identified by comparison to the commercial standards listed in the above cited method.

Statistical analysis was performed in R-studio. For each isotopologue (D0, D3, D4) a one-way ANOVA was conducted to test for differences between gene combinations (groups = 7, *n* = 8). For the drop out, the construct containing all genes was set as reference. For the step-wise reconstruction, the construct containing GFP only was set as reference. Post-hoc comparisons were carried out using two-tailed Dunnett’s test (multcomp package) to compare each construct directly against the reference group.

### Nicotiana benthamina metabolite analysis

*N. benthamiana* WT and quintuple *bbla-d’* knock-out seeds (*ΔNicGS*) were kindly provided by Nicola Patron^31^. Seeds of *N. benthamiana a622* knock-out line 3-3-1 (*ΔNaGR*) were kindly provided by Boaz Negin and Georg Jander^30^. Prior to germination, seeds of each line were surface sterilized according to Florentine *et al* 2016^59^. Several seeds per line were incubated in 500 µL 1% sodium hypochlorite solution for 2 min and afterwards the seeds were washed three times with 500 µL ultrapure water. The seeds were germinated in petri dishes on three layers of filter paper wetted with ultra-pure water. The plates were sealed with leucopore and kept in a growth cabinet under long day conditions (16 h light, 8 h dark) at day:night temperatures of 22 °C : 21 °C, 60% humidity and 200 µmol light intensity. The germinated seedlings were transferred to soil and the plants were grown in a growth cabinet under long day conditions (16 h light, 8 h dark) at day:night temperatures of 22 °C : 21 °C, 60% humidity and 200 µmol light intensity. Leaf and root samples were harvested from about 2-months-old plants. The root material was mostly collected from the soil-pot interface. To clean the roots, the soil was washed off with water, and excess water was removed by taping the roots on tissue paper. The harvested samples were flash-frozen in liquid nitrogen and were stored at −70 °C until tissue homogenization using chrome balls and a TissueLyzer (Qiagen). The homogenized samples were stored at −70 °C until further use. The samples were extracted and analysed by LC-MS as described above.

### *Nicotiana tabacum* metabolite analysis

*Nicotiana tabacum var. Samsun* were grown in a temperature-controlled glasshouse with 21/18 (SD ±3)°C Day/Night and 16 hour photoperiod. Further details of growing conditions is in the Supplementary Information. Plants’ apical and axillary growing tips were removed as soon as flower buds appeared (after about 1 month), these were regularly removed (monthly) until roots were harvested from 3 month-old plants and freeze dried. 10 mg of freeze dried root tissue was extracted into 1 mL MeOH, centrifuged (21,000 *x*g, 10 min) and loaded into LCMS vials for analysis. LC-MS analysis was carried out with Hydrophilic Interaction Liquid Chromatography (HILIC) separation on a Waters ACQUITY UPLC I-Class with detection performed on a Thermo Scientific Orbitrap Fusion™ Tribrid™ mass spectrometer. Details on LC-MS analytical procedures can be found in the Supplementary Information.

### Synthetic methods

Generally, glucoside intermediates were synthesised by glucosylation of the pyridine nitrogen using acetobromo-α-D-glucose, followed by deacetylation with aqueous HBr. Detailed synthetic methods and compound characterisation are available in the Supplementary Information.

## Acknowledgments

We thank: Sam Hart and Dr. Johan P. Turkenburg for assistance with X-ray data collection; Diamond Light Source Didcot UK for access to beamline I03 under grant number mx24948; Jared Cartwright, Tony Larson and the Biosciences Technology Facility; Ed Bergstrom, Karl Heaton, Mariela González, Jack Olsen and the Centre of Excellence in Mass Spectrometry; Matthew Davy and the NMR facility; Caragh Whitehead, Tessa Keenan, Zhouqian Jiang, Inesh Amarnath and Lachlan Waddell for preliminary work and materials; Peter Bølge Sørensen, Jason Daff and the University of York Horticulture team. We thank Hadrien Peyret and George Lomonossoff for the pHREAC-GFP plasmid, Georg Jander and Boaz Negin for the gift of the *N. benthamiana a622* line, and Nicola Patron for the *N. benthamiana* wildtype and quintuple *bbla-d’* knock-out line. We acknowledge funding from UKRI and BBSRC (MR/S01862X/1, MR/X010260/1, BB/T007222/1) and Independent Research Fund Denmark (DFF-FTP 2035-00038B, 0136-00410B).

## Author contributions

B.T.W.S. performed chemical synthesis; protein expression, purification and crystallography; and *in vitro* enzyme assays, pathway reconstruction and analysis. I.M.A. performed and analysed *Nicotiana benthamiana* transient expression experiments and ran the analysis of mutant plants. K.V. designed plant expression constructs and co-supervised I.M.A. Z.I. contributed to construct design, protein purification and method development. J. L. developed methods for chiral analysis. K.S.S. and C.D.S. synthesised deuterated substrates. M.A.F. supervised the design and execution of chemical synthesis. G.G. supervised protein preparation, crystallography and contributed to the X-ray structural analysis. F.G-F. supervised the *in planta* transient expression and metabolic analysis. B.R.L. identified gene candidates, supervised the biocatalytic reconstruction and coordinated the project. All authors contributed to data interpretation and writing the manuscript.

## Competing interests

C.D.S. and K.S.S. are inventors on a pending patent related to the production of deuterated compounds.

## Materials and Correspondence

Correspondence and material requests should be addressed to Benjamin R. Lichman.

## Data availability

Crystal structure files have been deposited on the RCSB Protein Data Bank under the accessions: NaGR *holo* (nicotinic acid-*N* glucoside) 9RDD; NicGS *holo* (nicotine *N*-glucoside) 9RDR; NicGS *apo* 9RUW. All other data are available in the main text or the supplementary information.

## Extended Data

**Extended Data Figure 1.**
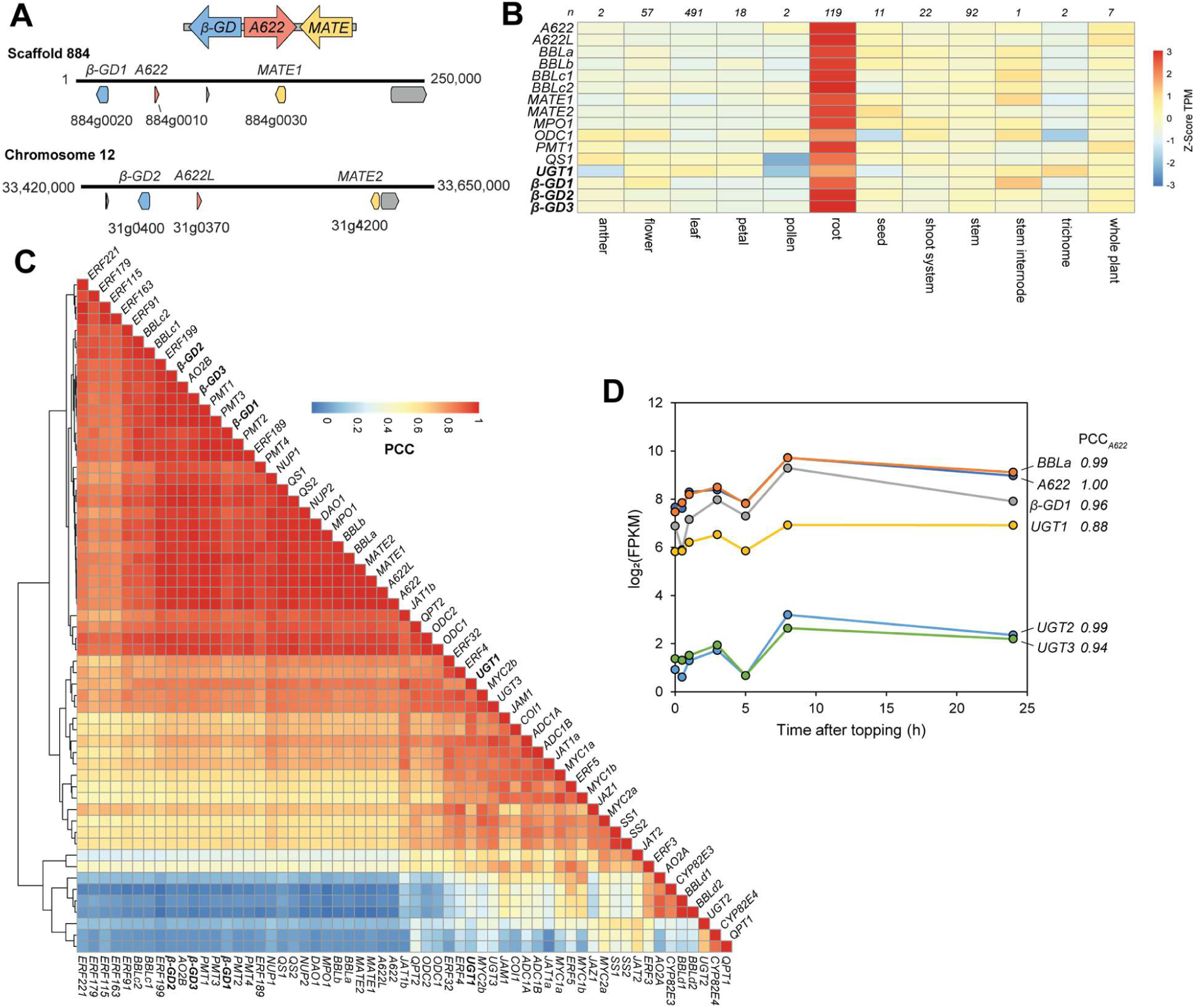
Identification of biosynthetic gene clusters and gene candidates. (**A**) Homeologous biosynthetic gene clusters identified in the *N. tabacum* 2017 genome assembly^36^ consisting of β-glucosidase (*β-GD*, blue), reductase (*A622*, red) and transporter (*MATE*, yellow) genes. Scaffold/chromosome number is described above the linear genome strand, with numbers at either end representing nucleotide position. Scaffold 884 is unplaced in this assembly but in the Sierro *et al* 2024 assembly it is part of chromosome 16^60^. The gene model between *A622* and *MATE1* in scaffold 884 (*884g0040*) encodes a protein which contains a domain with homology to a reverse transcriptase domain (PANTHER ID PTHR33116). (**B**) Heatmap of expression of selected nicotine genes including *β-GD* and *UGT* candidates across tissue types. Average expression calculated in each tissue type and values normalized per gene. Number of samples in each category above each column. (**C**) Clustered heatmap showing all-by-all correlations (PCC) of nicotine biosynthesis and related genes across 824 samples. TPM values obtained from PEO^39^. Details of genes available in Table S1. (**D**) Root expression levels of selected nicotine biosynthesis genes and UGT candidates following topping treatment, and their correlation (PCC) with A622 responses. Data obtained from Qin *et al* 2020^40^; gene details available in Table S1.

**Extended Data Figure 2.**
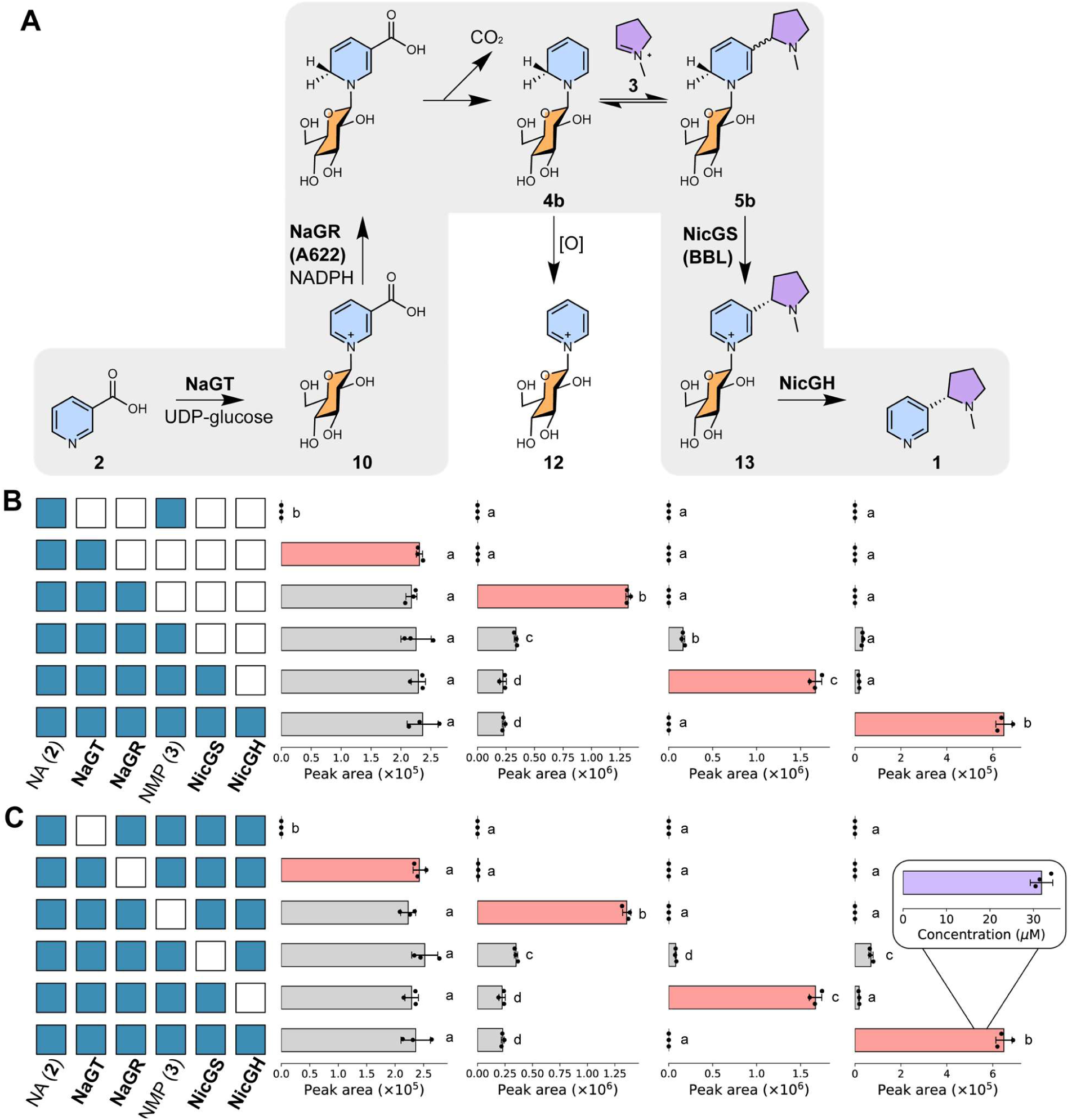
Quantification of *in vitro* reconstitution of nicotine biosynthesis. (**A**) Proposed chemical transformations in nicotine biosynthesis reconstitution. (**B-C**) Products of *in vitro* cascades showing (**B**) build up or (**C**) drop-out of components. Each row of bars corresponds to an *in vitro* reaction, with the matrix showing presence/absence (blue/white) of reaction components. Each column of bars shows the peak area of EICs (*m/z*±0.15) corresponding to the masses of the chemical depicted above (left-to-right: **10**, **12**, **13**, **1**). Compound identity was validated by standard retention time (Fig. 2) and MS^2^ spectra (Fig. S3). Red bars highlight key products. The letters above bars are significant groupings of peak area, determined per chemical with combined (B) and (C) data (*p*<0.05, Tukey post-hoc test, see Table S3). The concentration of nicotine produced (from 1 mM of **2**) was determined via an external standard curve; error bars show standard error (inset).

**Extended Data Figure 3.**
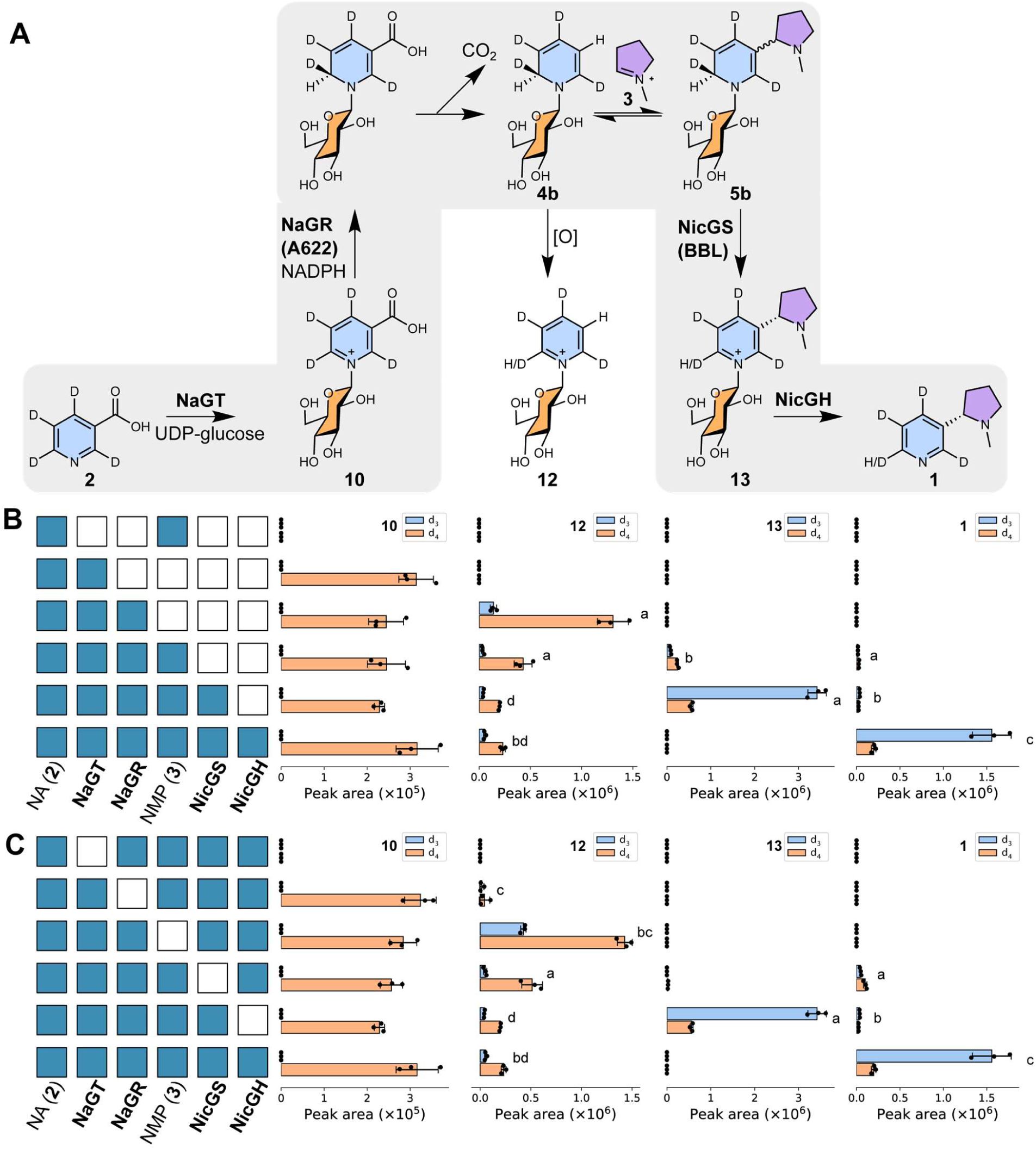
*In vitro* enzymatic cascade with nicotinic acid-d_4_ (2-d_4_). (**A**) Schematic of reaction cascade with deuterium labelling. (**B**) Build-up and (**C**) drop-out of *in vitro* cascades with **2**-d_4_ substrate. Bar charts show EIC peak areas of product (from L-to-R: **10**, **12, 13, 1**) isotopologues (all *m/z*±0.15 except for d_3_-**12** which was *m/z*±0.5 due to peak shape distortion). Reactions performed in triplicate, bars show mean peak area, points show individual measurements, error bars show standard deviation. Each row corresponds to an *in vitro* reaction, with the matrix showing presence/absence (blue/white) of reaction components. Letters above bars are significant groupings of d_3_/d_4_ isotopologue ratios, determined per chemical with combined panel **B** and **C** data (*p*<0.05, Tukey’s HSD test, see Table S4).

**Extended Data Figure 4.**
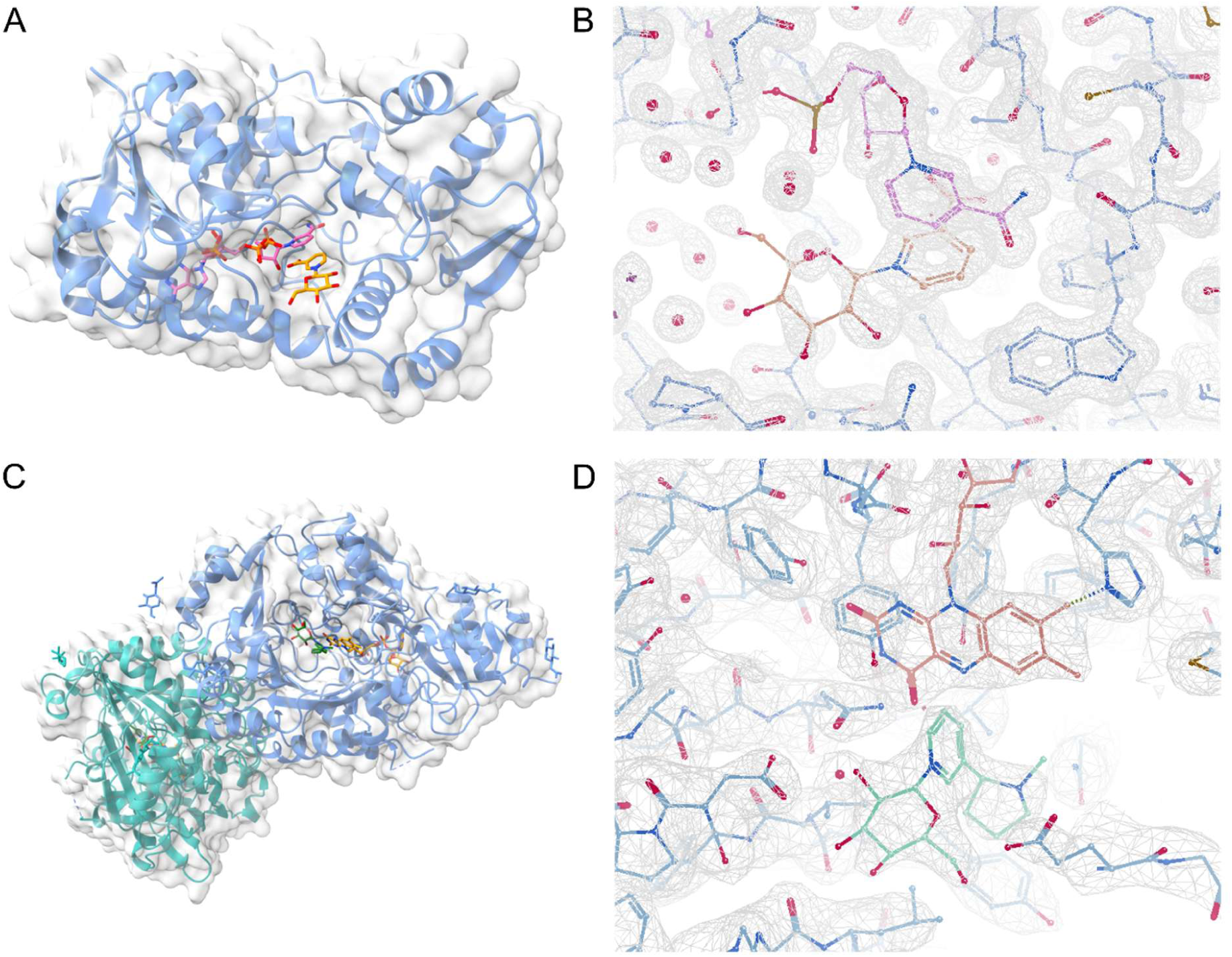
X-ray crystal structures of oxidoreductases in nicotine biosynthesis. (**A**) NaGR (A622) overall fold (9RDD). Ribbon shows backbone, solvent-excluded surface shown in transparent white. Ligand (nicotinic acid *N*-glucoside) shown as sticks. (**B**) Refined density (2Fo-Fc at 1 σ) around NaGR (A622) active site. (**C**) NicGS (BBLa) overall fold (9RDR). Ribbon shows backbone, solvent-excluded surface shown in transparent white. Ligand (nicotine glucoside) shown as sticks. (**D**) Refined density (2Fo-Fc at 1 σ) around NicGS (BBLa) active site.

**Extended Data Figure 5.**
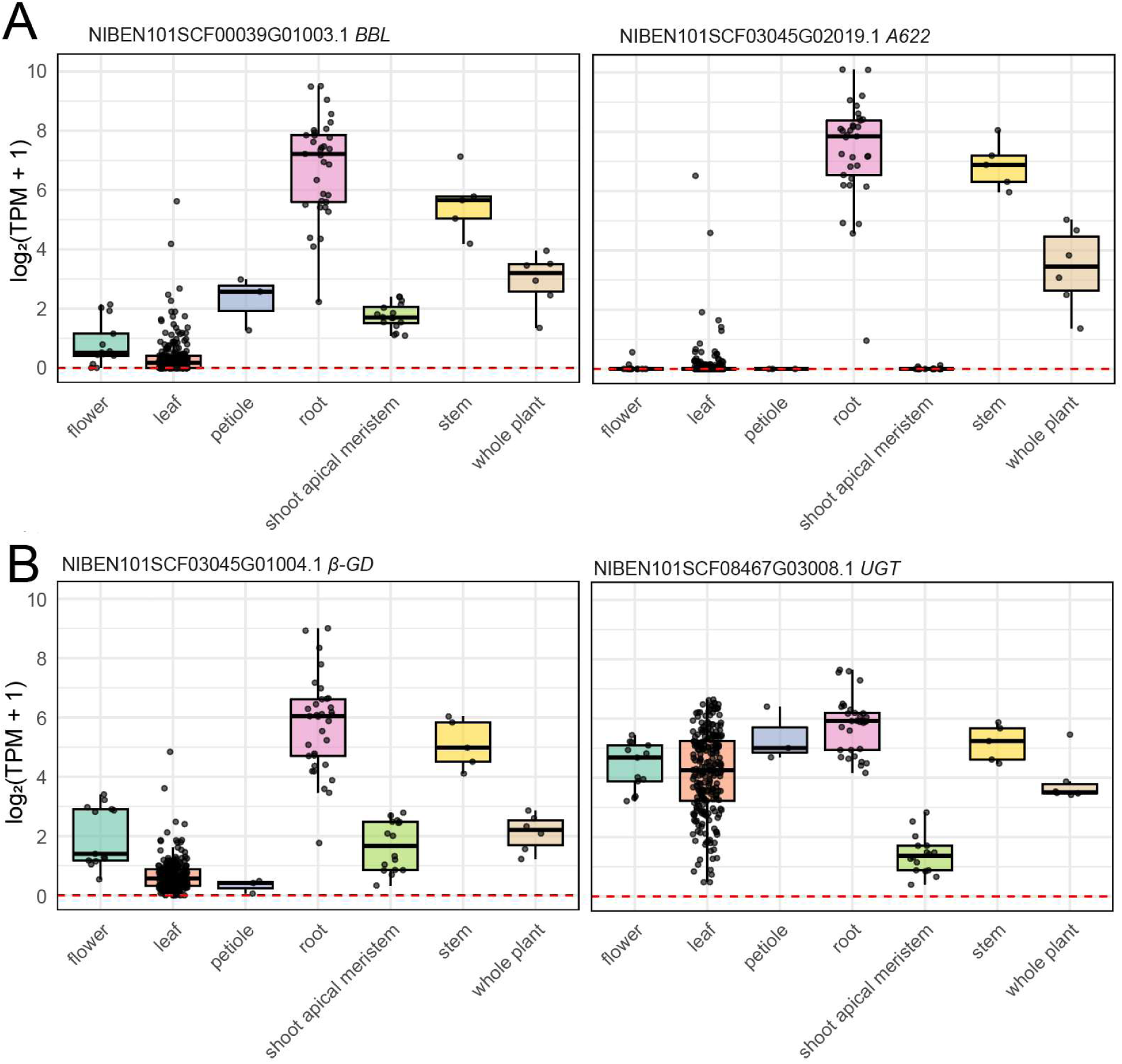
Expression of nicotine biosynthesis genes in *N. benthamiana*. Genes were identified as top blast hits using *N. tabacum* genes as queries. Expression data was compiled for *N. benthamiana* (Niben1.0.1) from the Plant Gene Expression Omnibus^39^. Expression values are in transcripts per million (TPM) and were calculated for seven tissue types across 476 samples. (**A**) Expression of the known genes *BBL* and *A622*. (**B**) Expression of the newly reported genes *β*-GD and UGT.

**Extended Data Figure 6.**
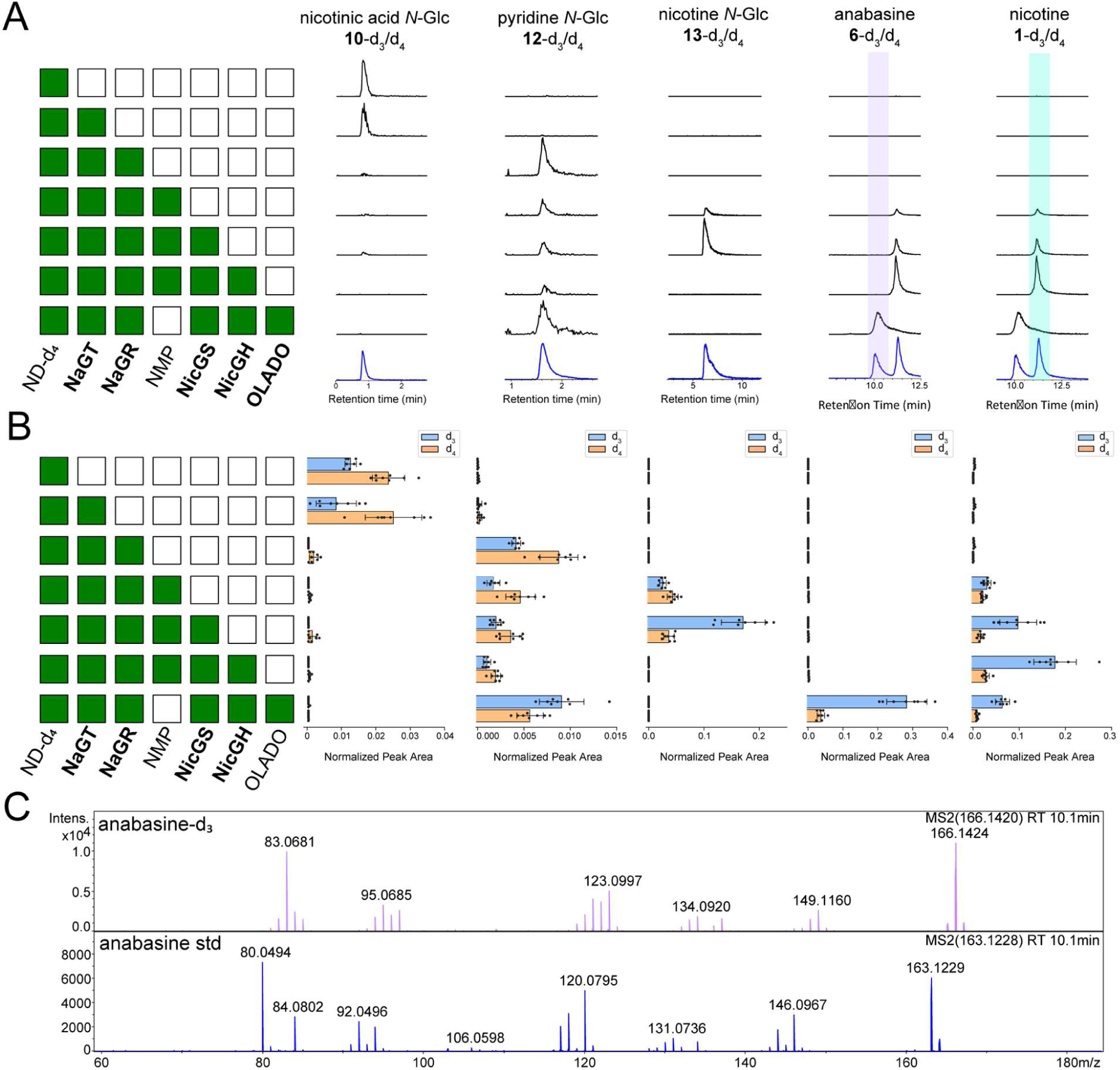
Stepwise pyridine alkaloid pathway reconstitution in leaves of *N. benthamiana*, as revealed by labelled precursor feeding with nicotinamide-d_4_ (ND-d_4_). The gene combination producing the *N-*methylpyrrolinium (**3**) (ODC, PMT and MPO) is abbreviated as NMP. (**A**) Extracted ion chromatograms corresponding to the labeled compounds indicated at the top (d_3_ and d_4_ versions together), matching with chemically verified standards (blue, unlabeled). Nicotine (**1**) and anabasine (**6**) standards (identical exact mass) were analyzed as a compound mix. (**B**) Individual quantification of d_3_ and d_4_ isotopologues corresponding to the individual compounds shown in panel (A). See Table S8 for statistical analysis. (**C**) Validation of anabasine peak identity by comparison of MS^2^ spectra between an unlabelled standard (blue, bottom) and the corresponding anabasine-d_3_ peak from the full gene combination where OLADO replaced the NMP genes (purple, top).

**Extended Data Figure 7.**
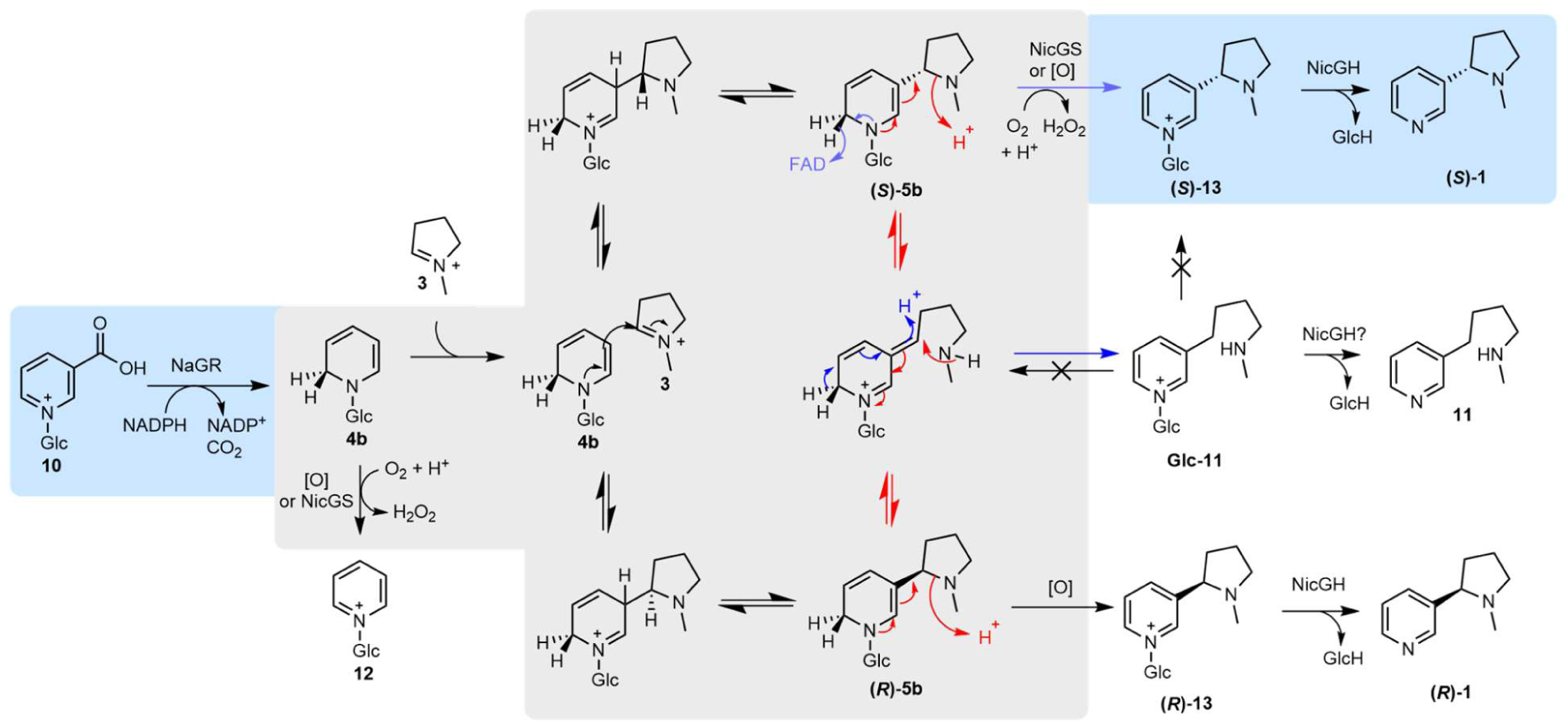
Putative activities of NicGS (BBL). Possible mechanistic routes towards (*S*)-**1** from **10.** The blue background shows key NaGR and NicGH reactions and major products. The grey background shows possible reactions that are either non-enzymatically or catalyzed by NicGS. Only compounds with blue or white background have been directly detected. To induce the stereoselective outcome, NicGS may operate through: dynamic kinetic resolution (catalyzing just purple arrows and allowing other intermediates to non-enzymatically equilibrate), epimerization of **5b** via a ring opened intermediate (catalyzing red and purple arrows), catalysis of the stereoselective Mannich reaction between **3** and **10** prior to oxidation, reversal of the non-desired Mannich reaction (opening (*R*)-**5b** into **3** and **10**). Blue arrows show the predicted mechanism of Glc-**11** formation.

**Extended Data Table 1.** Top correlating genes with A622 in *N. tabacum*.

| Rank | Gene | Names | PCC Value |
| --- | --- | --- | --- |
|  | NITAB4.5_0000884G0010 | *A622* | 1 |
| 1 | NITAB4.5_0006307G0010 | *BBLa* | 0.977 |
| 2 | NITAB4.5_0000884G0030 | *MATE1* | 0.973 |
| 3 | NITAB4.5_0003284G0010 | *PMT3* | 0.973 |
| 4 | NITAB4.5_0000031G0420 | *MATE2* | 0.973 |
| 5 | NITAB4.5_0000031G0370 | *A622L* | 0.969 |
| 6 | NITAB4.5_0003284G0040 | *PMT2* | 0.965 |
| 7 | NITAB4.5_0011232G0010 | *PMT4* | 0.96 |
| 8 | NITAB4.5_0000013G0390 | *PMT1* | 0.953 |
| 9 | NITAB4.5_0002522G0070 | *QS1* | 0.951 |
| 10 | NITAB4.5_0000795G0060 | ***β-GD3*** | 0.944 |
| 11 | NITAB4.5_0000316G0090 | *BBLb* | 0.939 |
| 12 | NITAB4.5_0008037G0020 | *QS2* | 0.937 |
| 13 | NITAB4.5_0000884G0020 | ***β-GD1*** | 0.926 |
| 14 | NITAB4.5_0000031G0400 | ***β-GD2*** | 0.917 |
| 15 | NITAB4.5_0000013G0380 |  | 0.916 |
| 16 | NITAB4.5_0002338G0020 | *AO2B* | 0.913 |
| 17 | NITAB4.5_0000983G0060 | *MPO1* | 0.909 |
| 18 | NITAB4.5_0000093G0170 |  | 0.9 |
| 19 | NITAB4.5_0010897G0020 |  | 0.888 |
| 20 | NITAB4.5_0007413G0010 |  | 0.888 |
| 21 | NITAB4.5_0000396G0230 |  | 0.883 |
| 22 | NITAB4.5_0000025G0290 | *DAO1* | 0.88 |
| 23 | NITAB4.5_0000180G0090 |  | 0.877 |
| 24 | NITAB4.5_0004658G0040 |  | 0.876 |

Pearson correlation coefficients (PCC), calculated from TPM values across 824 samples, extracted from Plant Gene Expression Omnibus<sup>39</sup>. Top 24 highest correlating genes shown in descending order. Names of nicotine biosynthesis genes are added with the newly identified $\beta$ -glucosidases in bold. Information about gene identity can be found in Table S1.

**Extended Data Table 2.** Top correlating genes with UGT1 in *N. tabacum*.

| Rank | Gene | Name | PCC Value |
| --- | --- | --- | --- |
|  | NITAB4.5_0006222G0020 | *UGT1* | 1 |
| 1 | NITAB4.5_0006307G0010 | *BBLa* | 0.792 |
| 2 | NITAB4.5_0000742G0010 | *QPT2* | 0.791 |
| 3 | NITAB4.5_0003263G0040 |  | 0.789 |
| 4 | NITAB4.5_0011232G0010 | *PMT4* | 0.785 |
| 5 | NITAB4.5_0003284G0040 | *PMT2* | 0.779 |
| 6 | NITAB4.5_0000884G0020 | ***β-GD1*** | 0.776 |
| 7 | NITAB4.5_0000884G0010 | *A622* | 0.776 |
| 8 | NITAB4.5_0000013G0390 | *PMT1* | 0.776 |
| 9 | NITAB4.5_0000884G0030 | *MATE1* | 0.775 |
| 10 | NITAB4.5_0003284G0010 | *PMT3* | 0.77 |
| 11 | NITAB4.5_0000180G0090 |  | 0.77 |
| 12 | NITAB4.5_0000093G0170 |  | 0.768 |
| 13 | NITAB4.5_0000031G0420 | *MATE2* | 0.766 |
| 14 | NITAB4.5_0000396G0230 |  | 0.762 |
| 15 | NITAB4.5_0003090G0030 | *ERF199/NIC1* | 0.761 |
| 16 | NITAB4.5_0004658G0040 |  | 0.759 |
| 17 | NITAB4.5_0010897G0020 |  | 0.756 |
| 18 | NITAB4.5_0012847G0010 |  | 0.752 |
| 19 | NITAB4.5_0001029G0080 | *BBLc2* | 0.748 |
| 20 | NITAB4.5_0000795G0060 | ***β-GD3*** | 0.746 |
| 21 | NITAB4.5_0002522G0070 | *QS1* | 0.743 |
| 22 | NITAB4.5_0000983G0060 | *MPO1* | 0.743 |
| 23 | NITAB4.5_0007413G0010 |  | 0.742 |
| 24 | NITAB4.5_0000881G0120 |  | 0.737 |

**Extended Data Table 3.** Top correlating genes with A622 in *N. benthamiana*.

| Rank | Gene | Annotation | PCC |
| --- | --- | --- | --- |
|  | NIBEN101SCF03045G02019.1 | *A622* | 1 |
| 1 | NIBEN101SCF03045G02020.1 | *MATE* | 0.986 |
| 2 | NIBEN101SCF00039G01003.1 | *BBL* | 0.979 |
| 3 | NIBEN101SCF03687G01012.1 | *QPT2* | 0.976 |
| 4 | NIBEN101SCF06783G00003.1 | *PMT1* | 0.973 |
| 5 | NIBEN101SCF15645G02001.1 | *MPO1* | 0.966 |
| 6 | NIBEN101SCF03045G01004.1 | *β-GD* | 0.963 |
| 7 | NIBEN101CTG08674G00002.1 | *BBL* | 0.952 |
| 8 | NIBEN101SCF00069G03001.1 |  | 0.947 |
| 9 | NIBEN101SCF06357G03002.1 | *BBL* | 0.946 |
| 10 | NIBEN101SCF02077G13016.1 | *MPO1* | 0.945 |
| 11 | NIBEN101SCF03006G02002.1 | *AO2* | 0.944 |
| 12 | NIBEN101SCF04788G00023.1 | *QPT2* | 0.941 |
| 13 | NIBEN101SCF00918G00001.1 | *AO2* | 0.923 |
| 14 | NIBEN101CTG11222G00002.1 |  | 0.921 |
| 15 | NIBEN101SCF02552G00006.1 |  | 0.917 |
| 16 | NIBEN101SCF01112G02010.1 |  | 0.913 |
| 17 | NIBEN101SCF00293G07014.1 | *BBL* | 0.912 |
| 18 | NIBEN101SCF11414G00007.1 |  | 0.912 |
| 19 | NIBEN101CTG08518G00003.1 | *PMT1* | 0.909 |
| 20 | NIBEN101SCF05437G04016.1 | *QS1* | 0.907 |
| 21 | NIBEN101SCF02530G05010.1 | *OLADO* | 0.9 |
| 22 | NIBEN101SCF05601G00019.1 | *β-GD* | 0.891 |
| 23 | NIBEN101SCF08195G07026.1 |  | 0.891 |
| 24 | NIBEN101SCF05918G03001.1 |  | 0.887 |
| Pearson correlation coefficients (PCC), calculated from TPM values across 476 samples, extracted from Plant Gene Expression Omnibus<sup>39</sup>. Highest correlating genes shown in descending order. Gene annotations performed using top blast hits using queries from *N. tabacum*. |  |  |  |

