## Supplementary Information for "Nicotine biosynthesis completed by cryptic activating glucosylation"

Supplementary Methods 2
Protein purification 2
NaGR (A622) expression and purification 2
NaGT (UGT) expression and purification 2
NicGS (BBLa) cloning, expression and purification 2
NicGH (*β*-GD1) expression and purification 4
OLADO expression and purification 4
LC-MS analytical procedures for *in vitro* reactions 5
Nicotiana tabacum growth 5
*Nicotiana tabacum* metabolite analysis 6
Synthetic methods 6
General methods 6
*N*-(tetra-*O*-acetyl-β-D-glucopyranosyl)-3-methyl nicotinate bromide 7
*N*-(β-D-glucopyranosyl)-nicotinic acid 7
*N*-(tetra-*O*-acetyl-β-D-glucopyranosyl)-pyridinium bromide 8
*N*-(β-D-glucopyranosyl)-pyridinium bromide 8
*N*-(tetra-*O*-acetyl-β-D-glucopyranosyl)-(*S*)-nicotine bromide 9
*N*-(β-D-glucopyranosyl)-(*S*)-nicotine bromide 9
*N*-(β-D-glucopyranosyl)-(±)-nicotine bromide 10
*N'*-boc-dihydrometanicotine 11
*N*-(tetra-*O*-acetyl-β-D-glucopyranosyl)-dihydrometanicotine bromide 11
*N*-(β-D-glucopyranosyl)-dihydrometanicotine bromide 12
Nicotinic acid-$d^6$ 12
Supplementary Figures 13
Supplementary References 33

**Supplementary Methods**

Protein purification

NaGR (A622) expression and purification

The sequence of NaGR (A622) was obtained based on the UniProt accession (IFRH_TOBAC). After multiple attempts to purify and crystalize NaGR (A622), an Δ5N-A622 N-terminal truncate construct was designed, with five N-terminal residues removed to improve stability, based on the observation of low confidence structural prediction in this region in an AlphaFold CoLab model[1]. The truncated gene was ordered from GenScript in the pET28a(+) vector, codon optimised for *E. coli* expression and with a non-cleavable C-terminal His-tag (Table S2). The plasmid was transformed into *E. coli* B834 protein expression strain (Novagen) via a standard heat shock protocol, inoculated into 5 mL LB media and incubated overnight (37 °C, 200 rpm). This was then used to inoculate 2x YT media (1 L) which was incubated (37 °C, 200 rpm) until OD600 = 0.3-0.4, after which it was incubated at a lower temperature (18 °C, 200 rpm) until OD600 = 0.6-0.9 when protein expression was induced with IPTG (500 µM). Cells were incubated overnight (16 h at 18 °C) prior to harvesting by centrifugation (4000 g, 20 min). The cell pellet was resuspended in Ni-NTA binding buffer (50 mL, 20 mM Tris-HCl, 20 mM imidazole, 300 mM NaCl, pH 7.4) containing cOmplete EDTA-free protease inhibitor, and lysed by cell disruption (one shot, 25 kPsi). The lysate was then clarified by centrifugation (25,000 xg, 40 min), before sterile filtration through 0.4 µM PES filters and loading onto a Cytiva HisTrap FF 5 ml column installed on an ÄKTA Pure protein chromatography system. The protein was eluted from the column by imidazole gradient elution (20 to 300 mM), and fractions were collected and analysed by SDS-PAGE. Fractions containing desired protein were combined, diluted in SEC buffer (20 mM Tris-HCl, 300 mM NaCl, pH 7.4), concentrated in Vivaspin protein concentrator (10 kDa MW cutoff) then loaded onto HiLoad Superdex 75 16/600 size exclusion column. Fractions from the column were collected and analysed by SDS-PAGE, and those containing desired protein were combined and concentrated to 10 mg $mL^{-1}$ for crystallography. Aliquots were also flash frozen in liquid nitrogen and stored at -70 °C for subsequent activity assays.

NaGT (UGT) expression and purification

The coding sequence for NaGT (UGT) was obtained (Nitab4.5_0006222g0020 from Edwards *et al* 2017[2]) (Table S1 and Table S2). An expression construct was ordered as a synthetic gene from Twist Bioscience, codon optimized for *E. coli*, and cloned into pET28a(+) within the NdeI and XhoI restriction enzyme sites. The expression and purification procedure was identical to A622 above, except that SoluBL21 *E. coli* expression strains were used (Genlantis).

NicGS (BBLa) cloning, expression and purification

The NicGS (BBLa) coding sequence (Nitab4.5_0006307g0010 from Edwards *et al* 2017[2]) was ordered as a gene fragment from GenScript with codon optimization for *E. coli* expression and subcloned into pET28b (Table S2). After initial expression tests in *E. coli* did not produce soluble protein, the expression strategy switched to a *Komagataella phaffii* secretion based system (formerly *Pichia pastoris*). Analysis by SignalP 6.0[3] indicated the presence of an N-terminal signal peptide with a cleavage site between amino acid 22 and 23. Therefore the gene was subsequently cloned into a TFPP-modified version of pPICZα to yield a construct without the first 22 native N-terminal amino acids but instead containing the α-factor signal sequence with a retained N-terminal hexahistidine tag. To achieve this the coding region was amplified from the pET28b plasmid by PCR using the forward and reverse primers: d(CAT CAC CAC CAC CAC GCG GTG ACC AAC CTG AGC) and d(GAG TTT TTG TTC TAG AAT TTA TTC GCT GCT ATA CTT GTG CTC). After PCR amplification, the DNA was cloned into a linearised pPICZB-NHis vector using InFusion (Takara Bio), which was confirmed by Sanger sequencing. The isolated plasmid DNA (~30 μg) was linearised with SacI (New England BioLabs), purified and adjusted to ~1μg/μl. *K. phaffii* X-33 expression strains were transformed with the linear DNA by electroporation and colonies were selected by transferring 150 μL of cell suspension onto YPDS plates containing 100 μg/mL of zeocin and incubating at 30 °C for 3 days. Colonies were further purified by re-streaking on YPD containing 100 μg/ml zeocin and incubation at 30°C, prior to storage as glycerol stocks at -80 °C.

To identify cell-lines capable of expression and secretion of the protein of interest, transformed cell-lines were resuscitated from a glycerol stock by inoculating onto YPD plates containing 100μg/ml zeocin and incubating at 30 °C for 72 hours. Single colonies from YPD/zeocin plates were used to inoculate 5 mL BMGY starter cultures with incubation performed at 30 °C, 200 rpm for 24 hours. Expression was initiated by the addition of 250 μL of BMGY starter culture into 5 mL of BMMY and following a brief mix, a 250 μL T=0hr sample was removed before incubation was continued at 30 °C, 180 rpm for 72 hours. Further sampling was performed every 24 hours by the removal of 250 μL of culture followed by a 250 μL addition of feed solution (BMMY containing 10% v/v methanol). The samples removed during the expression time-course (T=0, 24, 48 & 72hr post induction) were centrifuged at 5,000 g for 2 min at room temperature and the supernatant recovered and stored at -20 °C until analysis. Protein expression was examined by SDS-PAGE of extracted expression culture samples followed by western blotting using an anti-8xHIS-HRP conjugate antibody to identify cell lines with high expression of the protein of interest.

For large scale expression and purification, the frozen *K. phaffii* expression strain with validated expression and secretion of the protein of interest was re-streaked onto YPD-Zeocin plate, grown for 2 days and a single colony was used to inoculate BMGY media (30 mL) which was incubated (30 °C, 220 rpm, 24 hours). This starter culture was used to inoculate BMMY media (600 mL) and was incubated (30 °C, 180 rpm) feeding every 24 h with 5% v/v methanol in BMMY media to maintain MeOH at 0.5% v/v. After 144 h, cultures were harvested by centrifugation (6000 xg, 20 min) and the media was concentrated by tangential flow filtration (TFF, KrosFlo® KR2i), centrifuged (25,000 xg, 40 min), filtered through 0.8 μM PES filters, then loaded onto Cytiva HisTrap FF 5 ml column on an ÄKTA Pure protein chromatography system. The protein was eluted from the column by gradient elution with imidazole up to 300 mM, and fractions were collected and analysed by SDS-PAGE. Fractions containing the desired protein were combined, diluted in SEC buffer (20 mM Tris-HCl, 300 mM NaCl, pH 7.4), concentrated in Vivaspin protein concentrator (10 kDa MW cutoff) and then loaded onto HiLoad Superdex 75 16/600 size exclusion column. Fractions from the column were collected and analysed by SDS-PAGE. Those containing the desired protein were combined and centrifugally concentrated (VivaSpin) to 20 mg $mL^{-1}$ ready for crystallography. Aliquots were also flash frozen in liquid nitrogen and stored at -70 °C for subsequent crystallography experiments and activity assays.

NicGH (*β*-GD1) expression and purification

The coding sequence for NicGH (*β*-GD1) was obtained (Nitab4.5_0000884g0020 from Edwards *et al* 2017[2]) (Table S1 and Table S2). An expression construct was ordered as a synthetic gene from Twist Bioscience, codon optimized for *E. coli*, and cloned into pET28a(+) within the NdeI and XhoI restriction enzyme sites. Protein expression in *E. coli* was attempted as for the UGT above, however no soluble protein could be obtained. Therefore the gene was subsequently cloned into a TFPP-modified version of pPICZα (the EAEA repeat after the α-factor signal replaced by hexahistidine tag) to yield a construct without the first 22 native N-terminal amino acids but instead containing the α-factor signal sequence with a retained N-terminal hexahistidine tag. To achieve this the coding region was amplified out of the pET28a(+) plasmid with primers Fwd: CAT CAC CAC CAC CAC ATG TGC CAT TTA ACG GAT CAG G and Rev: GAG TTT TTG TTC TAG CTA TTT CTG TGC AGT ATG CGC T. The PCR product was then cloned into a linearised pPICZB-NHis vector using In-Fusion® (Takara Bio) then transformed into *E. coli* Stellar and positive clones were identified by colony PCR. Plasmid from a positive clone was then purified, confirmed by whole plasmid sequencing (Plasmidsaurus), linearized by PmeI digestion, and transformed into electrocompetent *K. phaffii* X-33. After 4 days of incubation, 12 clones were re-streaked overnight and selected for small scale expression screening. These were each inoculated into 2 mL BMGY in a 24 well block and incubated (30 °C, 220 rpm, 24 h). The cultures were then centrifuged (4000 xg, 5 min), the media removed, and cell pellets resuspended in 2 mL BMMY. Cultures were grown for 120 hours, and every 24 hours were centrifuged (4000 xg, 5 min). A portion (200 µL) of the media supernatant was removed and replaced with 10X methanol BMMY (200 µL). Clones showing successful protein expression were identified via an anti-His dot blot of media samples from the time course. The selected clone was then grown and the protein purified using the same method as described for the BBL above. Protein identity was validated by de-glycosylation, giving a band at the correct molecular weight and by trypsin digest and MALDI MS/MS. The protein was concentrated to 5 mg $mL^{-1}$ and aliquots were flash frozen in liquid nitrogen and stored at -70 °C for activity assays.

OLADO expression and purification

We obtained *Nicotiana tabacum* OLADO (A0A1S3ZKS9_TOBAC) in a pET-28a(+) expression vector[4] (Table S2). For protein expression, *E. coli* SoluBL21 (DE3) (Genlantis) was used. Single colonies transformed with plasmids containing genes of interest were used to inoculate overnight cultures with antibiotic selection (2xYT media, 37 °C, 200 rpm). These were diluted with 2xYT with antibiotics (OD600 = 0.1) and grown to OD600 = 0.4-0.6 (37 °C, 200 rpm). Then the cells were induced with IPTG (1 mM) and incubated overnight (~14 h, 200 rpm, 18 °C). Cells were harvested by centrifugation (3200 xg, 10 min, 4 °C), the supernatant removed, and the pellet resuspend in lysis buffer (10% v/v compared to original culture; 50 mM Tris-HCL, 50 mM glycine, 5% v/v glycerol, 0.5 M NaCl, 20 mM imidazole, pH 8; per 50 mL, 1x EDTA-free protease inhibitor and 10 mg lysozyme) and incubated (30 min, 4 °C). The cells were lysed through a cell disruptor (26 kPsi) and clarified by centrifugation (20 min, 35,000 xg, 4 °C). Purification was achieved through nickel affinity purification using an ÄKTA start FPLC. Clarified lysate was loaded onto a Ni-NTA column (5 mL) and then the column washed with binding buffer (50 mM Tris-HCL, 50 mM glycine, 5% v/v glycerol, 0.5 M NaCl, 40 mM imidazole, pH 8) and the protein eluted (50 mM Tris-HCL, 50 mM glycine, 5% v/v glycerol, 0.5 M NaCl, 500 mM imidazole, pH 8). A protein purification outcome was inspected by SDS-PAGE, and if a pure was protein obtained, samples were buffer exchanged into 50mM HEPES, pH 7.5 using a PD10 column, using a gravity method as per manufacturer instructions. The protein was concentrated by centrifugation (Amicon® Ultra-2 mL Centrifugal Filters, 30 kDa MWCO) prior to addition of glycerol (10% v/v), flash freezing and storage at –70 °C.

LC-MS analytical procedures for *in vitro* reactions

Enzyme assays were analysed by LC-MS, with Hydrophilic Interaction Liquid Chromatography (HILIC) separation using a Waters XBridge BEH Amide column (5 µm, 2.1 x 100 mm) on a Thermo Scientific Vanquish UHPLC. The mobile phase consisted of buffer A (water with 20 mM ammonium formate pH 3) and buffer B (90:10 acetonitrile:water with 20 mM ammonium formate pH 3). Separation was achieved using a gradient elution: 0-13 min, 100 % B to 76% B, 0.4 mL $min^{-1}$; 13-14 min, 50% B, 0.5 mL $min^{-1}$; 14-20 min, 100% B, 0.5 mL $min^{-1}$. Detection was performed on a Thermo Scientific LTQ XL Linear Ion Trap Mass Spectrometer. MS parameters were controlled by Thermo Xcalibur 4.1 software. Eluent was diverted to the H-ESI source between 1 and 19 min. The H-ESI source was maintained at 300 °C and a spray voltage of 3000 V (positive mode). Nitrogen was used as sheath, aux, and sweep gas, set to 60, 20, and 0 arbitrary units, respectively. Data was collected in data-dependent $MS^2$ mode. $MS^1$ data was collected in centroid mode, over a *m/z* range of 70-800. $MS^2$ data was collected in a data-dependent manner on the three most intense ions, but with targeted inclusion of expected biosynthetic intermediates. $MS^2$ spectra were recorded as centroid data in the ion trap detector, using a collisional induced dissociation (CID) fragmentation mode with precursor quadrupole isolation windows set to 2 *m/z*. CID data was collected at a normalised collision energy of 35%.

Nicotiana tabacum growth

*Nicotiana tabacum var. Samsun* were grown in Clover Professional Pot & Bedding (East Riding Horticulture) growing media in 7 cm pots, in a compartment within the University of York 'P Block' Venlo-style research greenhouse located at 53.94808, -1.05734 at 14.6 m elevation above sea level. The compartment had a floor area of 12.6 m by 6.6 m and a bench area of 45 $m^2$ elevated 0.75 m above the floor. The ridge of the greenhouse was oriented North–South. The eave (gutter) height was 4 m and the ridge extended 4.7 m above the floor. The greenhouse was mechanically ventilated with intake fans located in the North-facing side wall and screened ridge vents in the roof. The greenhouse section was clad with 4 mm thick tempered glass and outfitted with an automated shade curtain providing a reduction in light transmission of 50%, deployed when sunlight radiation flux exceeded 300 $W/m^2$ (measured at the greenhouse's external pyranometer). The greenhouse was equipped with a supplementary lighting system powered by Valoya (Helsinki, Finland) RX325 LED luminaires (Solray 385 spectrum) capable of providing a PAR flux of 230 $\mu mol\ m^{-2}s^{-1}$ (SD±20) at bench height. During the light period, the supplementary lighting system automatically turned on when the external sunlight radiation flux fell below 120 $W/m^2$, a 10 minute control delay was used to prevent excessive cycling of the supplementary lighting system. The greenhouse section was heated by a hot-water distribution system consisting of under-bench pipes. Air temperature at canopy height was maintained at 21/18 (SD±3) °C Day/Night. A 16 hour photoperiod was imposed and centred on solar noon by operating the supplementary lighting system when no sunlight was available. The relative humidity in the greenhouse compartment was ambient, typically within the region of 40-60%. All sensors and instruments were calibrated annually following manufacturers' specifications.

*Nicotiana tabacum* metabolite analysis

LC-MS analysis was carried out with Hydrophilic Interaction Liquid Chromatography (HILIC) separation using a Waters Atlantis Premier BEH Z-HILIC Column (1.7 µm, 2.1 mm X 100 mm) on a Waters ACQUITY UPLC I-Class. The mobile phase consisted of buffer A (water with 10 mM ammonium formate pH 3) and buffer B (90:10 acetonitrile:water with 10 mM ammonium formate pH 3). Separation was achieved using a gradient elution: 0-13 min, 100 % B to 76% B, 0.4 mL min$^{-1}$; 13-14 min, 50% B, 0.5 mL min$^{-1}$; 14-20 min, 100% B, 0.5 mL min$^{-1}$. Detection was performed on a Thermo Scientific Orbitrap Fusion™ Tribrid™ mass spectrometer. MS parameters were controlled by Thermo Xcalibur 4.1 software. Eluent was diverted to the H-ESI source between 1 and 19 min. The H-ESI source was maintained at 350 °C and a spray voltage of 3500 V (positive mode). Nitrogen was used as sheath, aux, and sweep gas, set to 50, 10, and 1 arbitrary units, respectively. The ion transfer tube was held at 325 °C. Data was collected in data-dependent MS$^2$ mode. MS$^1$ data was collected in profile mode using a cycle time of 0.6 s, over a *m/z* range of 70-800 with orbitrap resolution set to 120,000 (FWHM @ 200 *m/z*). Easy-IC internal calibration was used to reduce MS$^1$ mass errors to ~< 1 ppm. High resolution MS$^2$ data was collected in a data-dependent manner on the most intense ion, but with targeted inclusion of expected biosynthetic intermediates. MS$^2$ spectra were recorded as profile data in the orbitrap detector with a resolution set to 30,000, using a higher energy collisional dissociation (HCD) fragmentation mode with precursor quadrupole isolation windows set to 1.6 *m/z*. HCD data was collected at stepped normalised collision energies of 20, 35, and 60%. Dynamic exclusion was set to collect one HCD MS$^2$ scan from each precursor ion and to exclude further fragment scans from the same precursor for 6 s.

Synthetic methods

General methods

All commercially available reagents were used as received and were supplied by Sigma-Aldrich, Fisher Scientific and VWR International. Dihydrometanicotine was purchased from Toronto Research Chemicals. NMR spectra were recorded on a Jeol ECX-400 (400 MHz) spectrometer. All chemical shifts are quoted on the δ scale in ppm using residual solvent as the internal standard. Coupling constants (J) are reported in Hz with the following splitting abbreviations: s = singlet, d = doublet, t = triplet, q = quartet, m = multiplet, app = apparent, br = broad. Mass spectra (high-resolution) were obtained by the University of York Mass Spectrometry Service, using Electrospray Ionisation (ESI) on a Bruker Daltonics, Micro-tof spectrometer. Nominal and exact *m/z* values are reported in Daltons (Da).

N-(tetra-O-acetyl-β-D-glucopyranosyl)-3-methyl nicotinate bromide

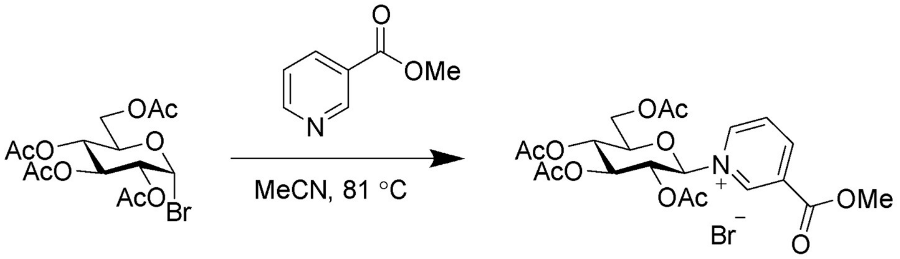


Acetobromo-α-D-glucose (2.0 g, 4.86 mmol, 1.0 equiv.) and methyl nicotinate (0.72 g, 5.25 mmol, 1.1 equiv.) were dissolved in anhydrous acetonitrile (10 mL) and refluxed for 16 h under nitrogen. The reaction mixture was concentrated *in vacuo* and the residue (2.79 g) was dissolved in DCM/$H_2O$ (20 mL, 1:1). The organic layer was washed 3x with $H_2O$, the aqueous phase was combined and freeze dried to yield a crude solid (1.289 g). This was dissolved in a minimal amount of water and purified on a Teledyne chromatography system with RediSep Gold® C18 Reversed Phase Column, 5.5 Gram, with gradient elution (10 to 50% MeCN in $H_2O$). Fractions were analysed by TLC (7:3:1.5 isopropanol : water : acetic acid), those containing the desired product were combined and freeze dried to yield *N*-(tetra-*O*-acetyl-β-D-glucopyranosyl)-3-methyl nicotinate bromide (0.400 g, 0.730 mmol, 15% yield, α/β=4:96). (Fig. S11) 1H NMR (400 MHz, D2O) δ 9.75 (t, J = 1.6 Hz, 1H), 9.39 (dt, J = 6.2, 1.5 Hz, 1H), 9.24 (dt, J = 8.2, 1.4 Hz, 1H), 8.38 (dd, J = 8.2, 6.2 Hz, 1H), 6.36 (d, J = 8.9 Hz, 1H), 5.72 (t, J = 9.4 Hz, 1H), 5.55 (t, J = 9.3 Hz, 1H), 5.47 – 5.39 (m, 1H), 4.55 – 4.45 (m, 2H), 4.43 – 4.34 (m, 1H), 4.08 (s, 3H), 2.15 (s, 6H), 2.09 (s, 3H), 1.96 (s, 3H). 13C NMR (101 MHz, D2O) δ 173.61, 172.83, 172.63, 171.79, 162.75, 149.24, 145.41, 143.69, 131.25, 129.01, 92.45, 75.03, 72.16, 71.98, 67.45, 61.72, 54.12, 20.20, 20.11, 20.04, 19.51. HRMS (ESI) = *m/z* $[M]^+$ calcd. for $C_{21}H_{26}NO_{11}^+$, 468.1500; found, 468.1503

N-(β-D-glucopyranosyl)-nicotinic acid

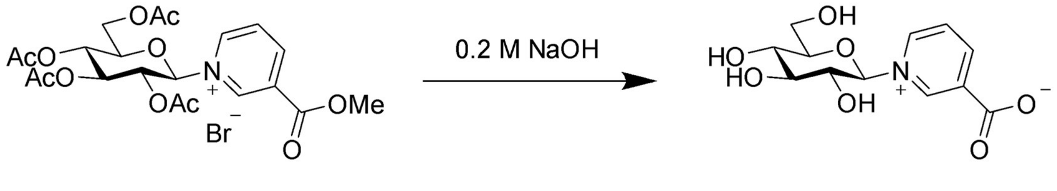


*N*-(tetra-*O*-acetyl-β-D-glucopyranosyl)-3-methyl nicotinate (88 mg, 0.188 mmol, 1 equiv.) was dissolved in aqueous NaOH (3.30 mL, 0.2 M 3.5 equiv.) and stirred for 16 h. Reaction progress followed by TLC (7:3:1.5 isopropanol:water:acetic acid) and further NaOH added until reaction reached completion. The reaction mixture was neutralised with Amberlite IRC 120 ($H^+$), the resin was filtered off and water removed *in vacuo*, yielding *N*-(β-D-glucopyranosyl)-nicotinic acid (24 mg, 0.084 mmol, 45% yield, α/β=4:96). (Fig. S12) Major anomer: $^1$H NMR (400 MHz, $D_2O$) δ 9.33 (d, *J* = 1.7 Hz, 1H), 9.12 – 9.07 (m, 1H), 9.02 – 8.97 (m, 1H), 8.21 (dd, *J* = 8.0, 6.2 Hz, 1H), 5.85 (d, *J* = 8.9 Hz, 1H), 4.03 – 3.63 (m, 6H). $^{13}$C NMR (400 MHz, D2O) δ 165.18, 145.40, 140.69, 140.42, 134.88, 125.63, 92.90, 77.31, 73.10, 71.79, 66.43, 58.18. HRMS (ESI) = *m/z* $[M+Na]^+$ calcd. for $C_{12}H_{16}NO_7Na^+$, 309.0819; found, 309.0810.

N-(tetra-O-acetyl-β-D-glucopyranosyl)-pyridinium bromide

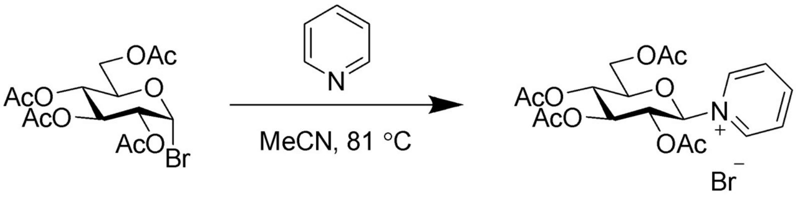


Acetobromo-α-D-glucose (1.0 g, 2.43 mmol, 1.0 equiv.) and pyridine (294 µL, 3.65 mmol, 1.5 equiv.) were dissolved in anhydrous acetonitrile (5 mL) and refluxed for 16 h under nitrogen. The reaction mixture was concentrated *in vacuo* and the residue was dissolved in DCM/$H_2O$ (20 ml, 1:1). The organic layer was washed 3x with $H_2O$, the aqueous phase was combined and freeze dried to yield a crude solid (1.000 g). This was dissolved in a minimal amount of water and purified on a Teledyne chromatography system with RediSep Gold® C18 Reversed Phase Column, 5.5 Gram, with gradient elution by increasing MeCN concentration (from 10 to 50% in $H_2O$). Fractions were analysed by TLC (7:3:1.5 isopropanol:water:acetic acid), those containing the desired product were combined and freeze dried to yield *N*-(tetra-*O*-acetyl-β-D-glucopyranosyl)-pyridinium bromide as a yellow-white crystalline solid (0.745 g, 1.519 mmol, 63% yield, α/β=12:88). (Fig. S13) Major anomer: $^{1}$H NMR (400 MHz, $D_2O$) δ 9.09 – 9.03 (m, 2H), 8.64 (tt, *J* = 7.8, 1.4 Hz, 1H), 8.14 – 8.08 (m, 2H), 6.14 (d, *J* = 9.1 Hz, 1H), 5.58 (t, *J* = 9.4 Hz, 1H), 5.40 (t, *J* = 9.3 Hz, 1H), 5.35 – 5.27 (m, 1H), 4.41 – 4.33 (m, 2H), 4.28 – 4.20 (m, 1H), 2.03 (s, 6H), 1.96 (s, 3H), 1.84 (s, 3H). $^{13}$C NMR (101 MHz, $D_2O$) δ 173.61, 172.85, 172.63, 171.77, 149.43, 142.41, 128.64, 92.14, 74.86, 72.19, 72.02, 67.50, 61.73, 20.18, 20.10, 20.04, 19.46. HRMS (ESI) = *m/z* $[M]^+$ calcd. for $C_{19}H_{24}NO_9^+$, 410.1446; found, 410.1454.

N-(β-D-glucopyranosyl)-pyridinium bromide

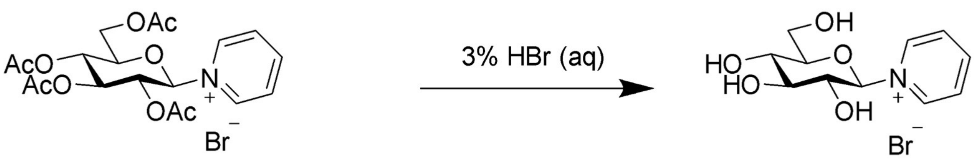


*N*-(tetra-*O*-acetyl-β-D-glucopyranosyl)-pyridinium bromide (140 mg, 0.286 mmol, 1 equiv.) was dissolved in 3% aqueous HBr (14 ml) and stirred for 24 h. Reaction progress followed by TLC (7:3:1.5 isopropanol:water:acetic acid). The reaction mixture was concentrated *in vacuo*, and the yellow oily residue was re-dissolved into EtOH, before using $Et_2O$ to precipitate *N*-(β-D-glucopyranosyl)-pyridinium bromide (43 mg, 0.133 mmol, 47% yield, α/β=12:88). (Fig. S14) 1H NMR (400 MHz, $D_2O$) δ 9.08 – 9.03 (m, 2H), 8.74 – 8.67 (m, 1H), 8.21 – 8.15 (m, 2H), 5.80 (d, J = 8.7 Hz, 1H), 3.99 (dd, J = 12.2, 1.9 Hz, 1H), 3.90 (d, J = 5.4 Hz, 1H), 3.87 – 3.82 (m, 2H), 3.77 (t, J = 8.9 Hz, 1H), 3.67 (dt, J = 14.2, 9.1 Hz, 2H). 13C NMR (101 MHz, D2O) δ 148.35, 147.36, 142.41, 142.12, 128.18, 128.04, 95.07, 91.14, 79.50, 77.85, 75.36, 74.06, 72.30, 70.33, 68.73, 68.60, 60.43, 60.18. HRMS (ESI) = *m/z* $[M]^+$ calcd. for $C_{11}H_{16}NO_5^+$, 242.1023; found, 242.1026.

N-(tetra-O-acetyl-β-D-glucopyranosyl)-(S)-nicotine bromide

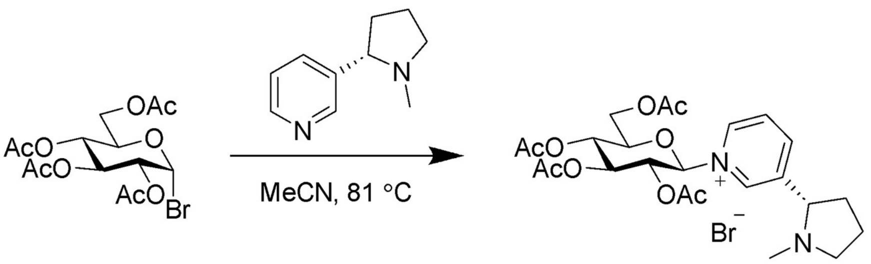


Acetobromo-α-D-glucose (1.0 g, 2.43 mmol, 1.0 equiv.) and (*S*)-nicotine (585 µl, 3.65 mmol, 1.5 equiv.) were dissolved in anhydrous acetonitrile (5 ml) and refluxed for 16 h under nitrogen. The reaction mixture was concentrated *in vacuo* and the residue was dissolved in DCM/$H_2O$ (20 ml, 1:1). The organic layer was washed 3x with $H_2O$, the aqueous phase was combined and freeze dried to yield a crude solid (1.153 g). This was dissolved in a minimal amount of water and purified on a Teledyne chromatography system with RediSep Gold® C18 Reversed Phase Column, 5.5 Gram, with gradient elution by increasing MeCN concentration (from 10 to 50% in $H_2O$). Fractions were analysed by TLC (7:3:1.5 isopropanol : water : acetic acid), those containing the desired product were combined and freeze dried to yield *N*-(tetra-*O*-acetyl-β-D-glucopyranosyl)-(*S*)-nicotine bromide (0.805 g, 1.404 mmol, 58% yield, α/β=21:79). (Fig. S15) 1H NMR (400 MHz, D2O) δ 9.15 (t, J = 1.5 Hz, 2H), 9.07 (ddt, J = 9.1, 6.3, 1.4 Hz, 4H), 8.72 (dt, J = 7.9, 1.6 Hz, 2H), 8.70 – 8.63 (m, 1H), 8.67 – 8.59 (m, 1H), 8.20 (ddd, J = 8.6, 6.1, 2.5 Hz, 3H), 6.83 (d, J = 2.8 Hz, 1H), 6.24 (d, J = 9.1 Hz, 2H), 5.71 (t, J = 9.4 Hz, 2H), 5.60 (t, J = 3.0 Hz, 1H), 5.53 (t, J = 9.2 Hz, 2H), 5.47 – 5.35 (m, 3H), 5.29 (dtt, J = 13.7, 5.5, 3.2 Hz, 1H), 5.23 – 5.06 (m, 2H), 4.98 – 4.84 (m, 1H), 4.78 – 4.64 (m, 3H), 4.54 – 4.43 (m, 5H), 4.46 – 4.32 (m, 2H), 4.36 – 4.28 (m, 1H), 3.63 (t, J = 8.4 Hz, 4H), 3.24 (ddd, J = 10.0, 6.4, 3.8 Hz, 3H), 2.70 (s, 1H), 2.58 – 2.35 (m, 6H), 2.28 – 2.19 (m, 13H), 2.18 (s, 3H), 2.17 (s, 1H), 2.16 – 2.04 (m, 25H), 2.08 – 1.92 (m, 15H), 1.90 (s, 1H), 1.94 – 1.78 (m, 3H). 13C NMR (101 MHz, D2O) δ 173.72, 173.59, 172.84, 172.64, 171.84, 170.94, 148.38, 147.33, 141.91, 141.40, 140.77, 140.57, 128.69, 128.42, 92.31, 87.31, 74.92, 72.08, 68.54, 67.53, 67.44, 66.70, 61.71, 61.30, 56.52, 39.30, 39.25, 33.87, 22.21, 20.21, 20.10, 20.04, 19.64, 19.46. HRMS (ESI) = *m/z* $[M+H]^+$ calcd. for $C_{24}H_{33}N_2O_9^+$, 493.2181; found, 493.2182

N-(β-D-glucopyranosyl)-(S)-nicotine bromide

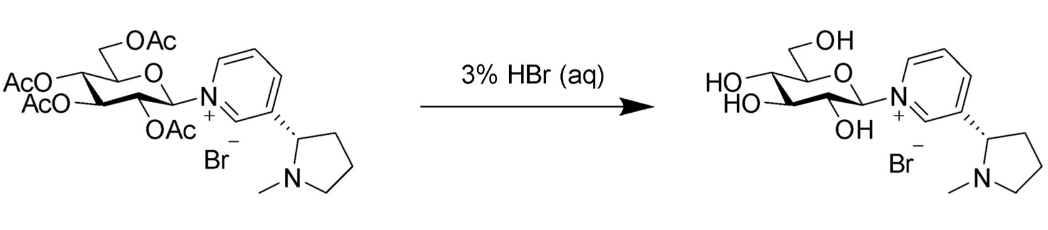


*N*-(tetra-*O*-acetyl-β-D-glucopyranosyl)-(*S*)-nicotine bromide (139 mg, 0.242 mmol, 1 equiv.) was dissolved in 3% aqueous HBr (14 ml) and stirred for 24 h. Reaction progress followed by TLC (7:3:1 isopropanol:water:triethylamine). The reaction mixture was concentrated *in vacuo*, and the yellow oily residue was re-dissolved into EtOH, before using $Et_2O$ to precipitate *N*-(β-D-glucopyranosyl)-(*S*)-nicotine bromide (76 mg, 0.188 mmol, 78% yield, α/β=21:79). (Fig. S16).

[1]H NMR (400 MHz, $D_2O$) δ 9.36 (s, 1H), 9.23 – 9.18 (m, 1H), 8.92 (dt, *J* = 8.0, 1.6 Hz, 1H), 8.32 (ddd, *J* = 11.4, 8.2, 6.3 Hz, 1H), 5.88 (d, *J* = 8.7 Hz, 1H), 4.02 – 3.63 (m, 6H), 3.42 (d, *J* = 15.6 Hz, 1H), 2.90 (s, 3H), 2.74 (dt, *J* = 13.8, 6.8 Hz, 1H), 2.50 – 2.28 (m, 3H). 13C NMR (101 MHz, D2O) δ 143.82, 129.04, 95.38, 79.59, 75.29, 74.08, 68.57, 60.28, 21.56. HRMS (ESI) = *m/z* $[M+H]^+$ calcd. for $C_{16}H_{25}N_2O_5^+$, 325.1758; found, 325.1757

<u>*N*-(β-D-glucopyranosyl)-(±)-nicotine bromide</u>

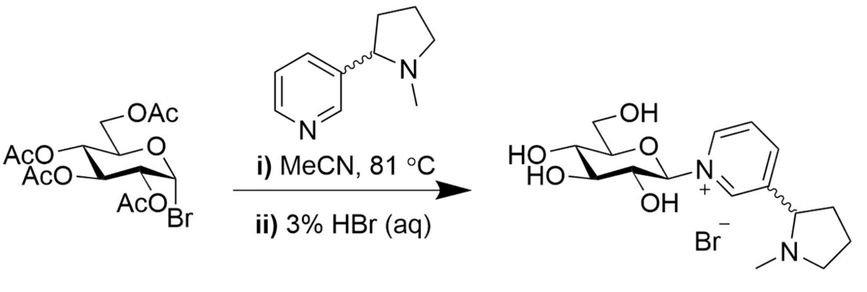


Acetobromo-α-D-glucose (115 mg, 0.280 mmol, 1.0 equiv.) and (±)-nicotine (50 mg, 0.309 mmol, 1.1 equiv.) were dissolved in anhydrous acetonitrile (1 ml) and stirred at 50 ºC for 24 h under nitrogen. The reaction mixture was concentrated *in vacuo* and the residue was dissolved in DCM/$H_2O$ (20 ml, 1:1). The organic layer was washed 3x with $H_2O$, the aqueous phase was combined and freeze dried to yield a crude solid. This was dissolved in a minimal amount of water and purified on a Teledyne chromatography system with RediSep Gold® C18 Reversed Phase Column, 5.5 Gram, with gradient elution by increasing MeCN concentration (from 10 to 50% in $H_2O$). Fractions were analysed by TLC (7:3:1 isopropanol : water : triethylamine), those containing the desired product were combined and freeze dried. NMR showed the product had already undergone partial deprotection, so the freeze dried solid was dissolved directly in 3% aqueous HBr (7 mL) and stirred for 24 h. Reaction progress followed by TLC (7:3:1.5 isopropanol : water : acetic acid). The reaction mixture was concentrated *in vacuo*, and the yellow oily residue was re-dissolved into EtOH, before using $Et_2O$ to precipitate *N*-(β-D-glucopyranosyl)-(±)-nicotine bromide (9 mg. 0.022 mmol, 8% yield). (Fig. S17) 1H NMR (700 MHz, D2O) δ 9.36 (d, J = 8.7 Hz, 1H), 9.23 (ddd, J = 6.1, 4.6, 1.3 Hz, 1H), 8.96 – 8.91 (m, 1H), 8.56 (d, J = 8.2 Hz, 0H), 8.35 (ddd, J = 7.9, 4.5, 1.4 Hz, 1H), 8.03 (t, J = 6.7 Hz, 0H), 5.89 (dt, J = 8.9, 1.8 Hz, 1H), 4.67 – 4.62 (m, 0H), 4.04 – 3.97 (m, 1H), 3.96 – 3.84 (m, 2H), 3.81 (tdd, J = 9.2, 2.3, 1.2 Hz, 1H), 3.76 – 3.69 (m, 1H), 3.65 (dtd, J = 27.5, 9.1, 1.3 Hz, 1H), 3.45 (s, 1H), 3.42 – 3.37 (m, 0H), 2.91 (s, 3H), 2.88 (s, 1H), 2.74 (s, 1H), 2.74 – 2.67 (m, 1H), 2.45 (s, 2H), 2.37 (s, 4H). 13C NMR (176 MHz, D2O) δ 143.72, 142.53, 128.97, 128.92, 95.33, 95.27, 79.60, 79.53, 75.25, 75.24, 74.05, 74.03, 69.22, 68.55, 68.51, 60.21, 60.20, 56.68, 56.42, 38.53, 30.89, 30.51, 21.52, 21.49.

*N'*-boc-dihydrometanicotine

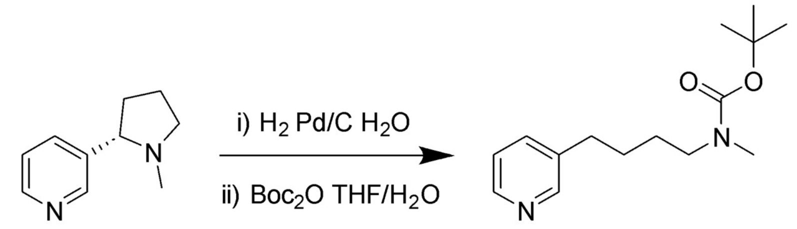


Nicotine (1.0 mL, 1.010 g. 6.23 mmol) dissolved in water (20 mL) Pd on carbon (250 mg, 10% palladium) added and solution degassed by evacuation and back fill with nitrogen. $H_2$ gas supplied by balloon, reaction progress was followed by TLC (7:3 isopropanol : water) and the reaction was stopped after 3 days. Boc anhydride (1.3 g, ) and DIPEA (1.1 mL, ) dissolved in THF (10 mL) and added to the reaction mixture under stirring. After 20 min the Boc protection appeared to be complete by TLC (9:1 DCM : MeOH). The reaction mixture was saturated with NaCl and THF layer removed. The THF layer was washed with brine (3 x 10 mL), then dried *in vacuo*, product purified by gradient elution from silica gel 0-4% MeOH in DCM. Fractions were analysed by TLC (9:1 DCM : MeOH), those containing the desired product were combined and dried *in vacuo* to yield *N'*-boc-dihydrometanicotine as a yellow oil (181 mg, 0.686 mmol, 11% yield). (Fig. S18) $^1$H NMR (400 MHz, CHLOROFORM-*D*) δ 8.47 – 8.38 (m, 2H), 7.47 (dd, *J* = 7.7, 2.2 Hz, 1H), 7.19 (dd, *J* = 7.8, 4.7 Hz, 1H), 3.22 (s, 2H), 2.80 (s, 3H), 2.63 (t, *J* = 7.3 Hz, 2H), 1.64 – 1.50 (m, 4H), 1.43 (s, 9H). $^{13}$C NMR (400 MHz, CHLOROFORM-*D*) δ 155.60, 149.69, 147.14, 135.58, 123.09, 79.04, 77.04, 48.31, 33.86, 32.43, 28.26, 27.91, 27.16. HRMS (ESI) = *m/z* $[M+H]^+$ calcd. for $C_{15}H_{25}N_2O_2^+$, 265.1911; found, 265.1913

*N*-(tetra-*O*-acetyl-β-D-glucopyranosyl)-dihydrometanicotine bromide

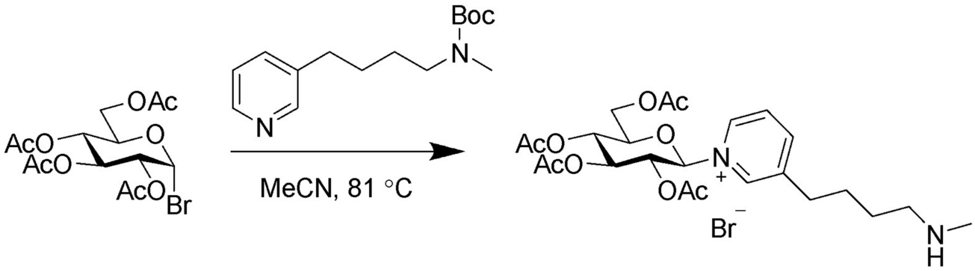


Acetobromo-α-D-glucose (334 mg, 0.812 mmol, 1.2 equiv.) and *N'*-boc-dihydrometanicotine (176 mg, 0.667 mmol, 1 equiv.) were dissolved in anhydrous acetonitrile (5 ml) and refluxed for 16 h under nitrogen. The reaction mixture was concentrated *in vacuo* and the residue was dissolved in DCM/$H_2O$ (20 ml, 1:1). The organic layer was washed 3x with $H_2O$, the aqueous phase was combined and freeze dried to yield a crude solid. This was dissolved in a minimal amount of water and purified on a Teledyne chromatography system with RediSep Gold® C18 Reversed Phase Column, 5.5 Gram, with gradient elution by increasing MeCN concentration (from 10 to 50% in $H_2O$). Fractions were analysed by TLC (7:3 isopropanol:water), those containing the desired product were combined and freeze dried to yield *N*-(tetra-*O*-acetyl-β-D-glucopyranosyl)-dihydrometanicotine bromide (79 mg, 0.169 mmol, 17% yield, α/β=38:62). (Fig. S19) 1H NMR (400 MHz, D2O) δ 9.08 (d, J = 1.8 Hz, 1H), 8.95 (dq, J = 6.3, 1.9 Hz, 2H), 8.58 (dt, J = 8.0, 1.5 Hz, 1H), 8.07 (dd, J = 8.1, 6.2 Hz, 2H), 6.17 (d, J = 8.8 Hz, 1H), 5.67 (t, J = 9.4 Hz, 1H), 5.58 –

5.45 (m, 2H), 5.45 – 5.33 (m, 2H), 4.79 (s, 2H), 4.50 – 4.35 (m, 3H), 4.35 – 4.26 (m, 1H), 3.05 (t, J = 7.1 Hz, 4H), 2.94 (td, J = 7.3, 2.6 Hz, 4H), 2.67 (s, 4H), 2.13 – 2.07 (m, 9H), 2.04 (s, 3H), 1.91 (s, 3H), 1.83 – 1.66 (m, 8H). 13C NMR (101 MHz, D2O) δ 173.71, 173.58, 172.83, 172.65, 171.76, 171.49, 149.32, 148.30, 143.97, 143.36, 141.41, 140.78, 140.48, 139.28, 128.08, 127.83, 92.16, 87.03, 74.96, 74.89, 72.12, 71.99, 68.45, 67.53, 66.65, 61.72, 61.21, 48.59, 32.71, 31.51, 31.45, 26.71, 26.61, 24.85, 24.80, 20.30, 20.24, 20.17, 20.10, 20.04, 19.65, 19.48. HRMS (ESI) = *m/z* $[M+H]^+$ calcd. for $C_{24}H_{35}N_2O_9^+$, 495.2337; found, 495.2341

*N*-(β-D-glucopyranosyl)-dihydrometanicotine bromide

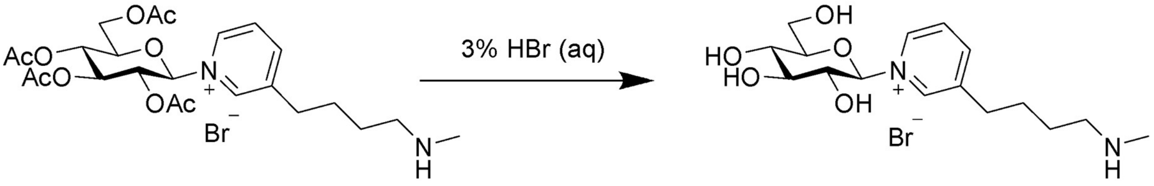


*N*-(tetra-*O*-acetyl-β-D-glucopyranosyl)-dihydrometanicotine bromide (27 mg, 0.047 mmol, 1 equiv.) was dissolved in 3% aqueous HBr (2.7 ml) and stirred for 24 h at 45 ºC. The reaction mixture was diluted with water, before being removed *in vacuo*. The yellow oily residue was re-dissolved into a minimum amount of EtOH, before using $Et_2O$ to precipitate *N*-(β-D-glucopyranosyl)-dihydrometanicotine bromide (9 mg, 0.022 mmol, 47% yield, α/β=38:62). (Fig. S20) 1H NMR (400 MHz, D2O) δ 8.98 – 8.82 (m, 2H), 8.69 – 8.43 (m, 1H), 8.03 (ddd, J = 11.0, 8.0, 6.2 Hz, 1H), 5.72 (d, J = 8.7 Hz, 1H), 3.99 – 3.56 (m, 5H), 3.03 (t, J = 7.2 Hz, 2H), 2.92 (t, J = 7.2 Hz, 2H), 2.66 (s, 3H), 1.83 – 1.65 (m, 4H). 13C NMR (101 MHz, D2O) δ 148.20, 143.23, 141.27, 139.92, 127.69, 95.06, 79.47, 75.35, 74.01, 72.31, 68.68, 60.35, 48.61, 32.69, 31.46, 26.57, 24.82.

Nicotinic acid-$d^6$

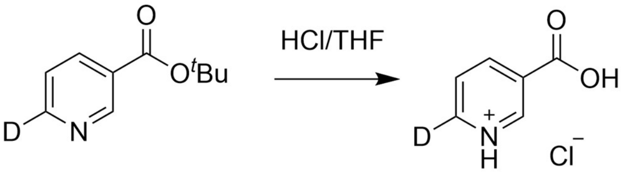


Nicotinic acid-$d^6$ hydrochloride salt was generated from the corresponding tert-butyl ester and used directly. Briefly, nicotinic acid-$d^6$ tert-butyl ester (10 mg, 80 µmol) was suspended in a mixture of hydrochloric acid (2 M, 0.5 mL) and THF (0.5 mL) and the mixture stirred for 2 h. The solvent was then removed *in vacuo* to provide the product which was used directly in further experiments. $^1$H NMR [nicotinic acid-$d^6$ tert-butyl ester] (400 MHz, $CDCl_3$): δ 9.15-9.17 (1H, m, H2), 8.21-8.26 (1H, m, H4), 7.33-7.38 (1H, m, H3), 1.61 (9H, s, OtBu); $^{13}$C NMR (101 MHz, $CDCl_3$): δ 164.46 (-$CO_2{}^t$Bu), 150.94 (C2), 145.08 (C6), 136.99 (C4), 127.80 (C2), 123.06 (C5), 82.13 (-$CMe_3$), 28.23 (Me); HRMS: *m/z* ($ESI^+$, ionises as acid): Calcd. for $C_6H_4DNO_2$ $[M+H]^+$ = 125.0456, Obs. 125.0456.

`

### Supplementary Figures

**Predicted localizations:** Lysosome/Vacuole
**Predicted signals:** Signal peptide

| Localization | Cytoplasm | Nucleus | Extracellular | Cell membrane | Mitochondrion | Plastid | Endoplasmic reticulum | Lysosome/Vacuole | Golgi apparatus | Peroxisome |
|---|---|---|---|---|---|---|---|---|---|---|
| **Probability** | 0.1510 | 0.0803 | 0.4261 | 0.2885 | 0.0309 | 0.2011 | 0.5265 | 0.7244 | 0.2148 | 0.0209 |

Predicted Signals: Signal peptide

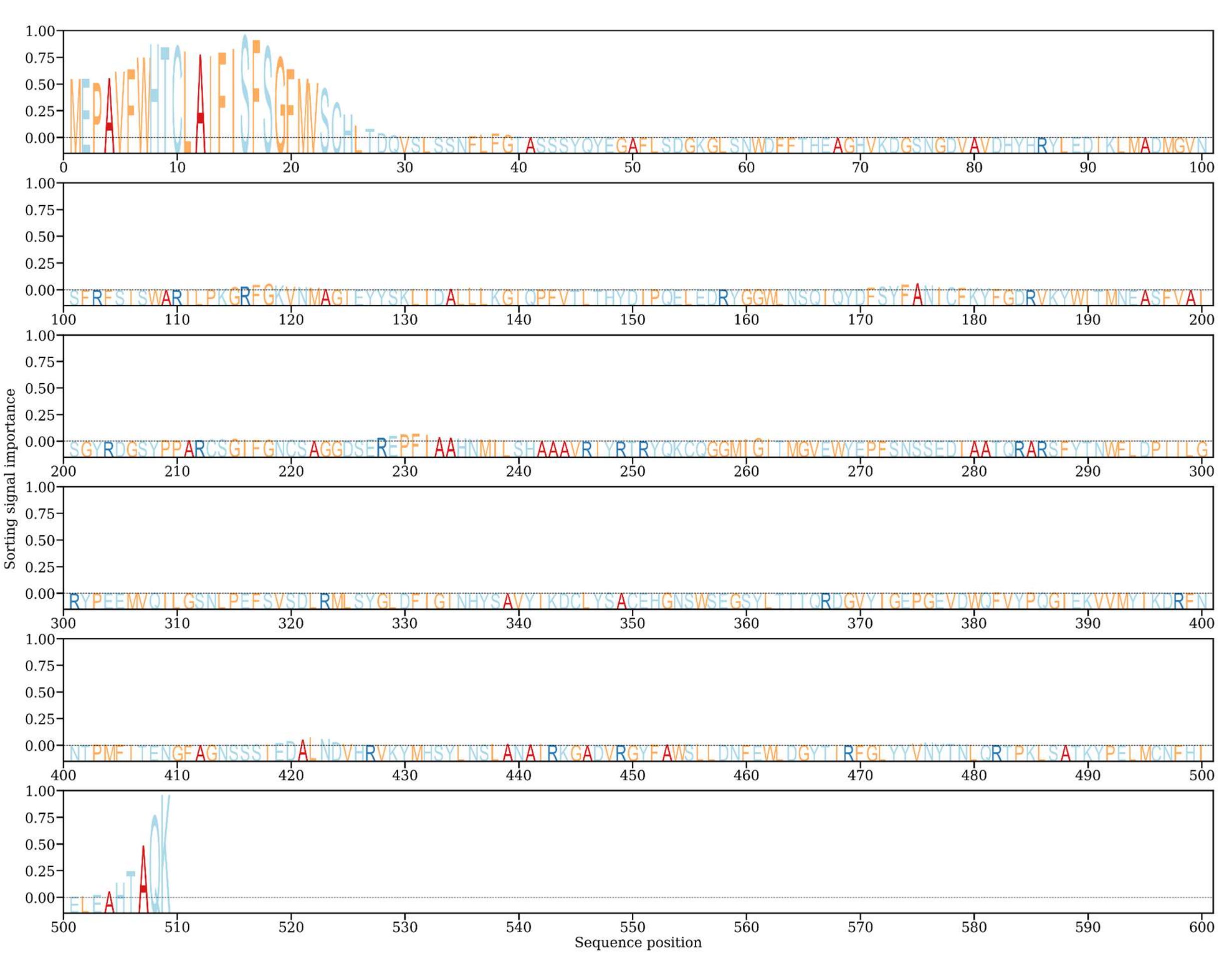


**Figure S1. Predicted vacuolar subcellular location of *β-GD1*.** Sequence based prediction of subcellular localization using DeepLoc2.0[5]. Standard output presented highlighting localization probability (top) and predicted signal peptide sequence (bottom).

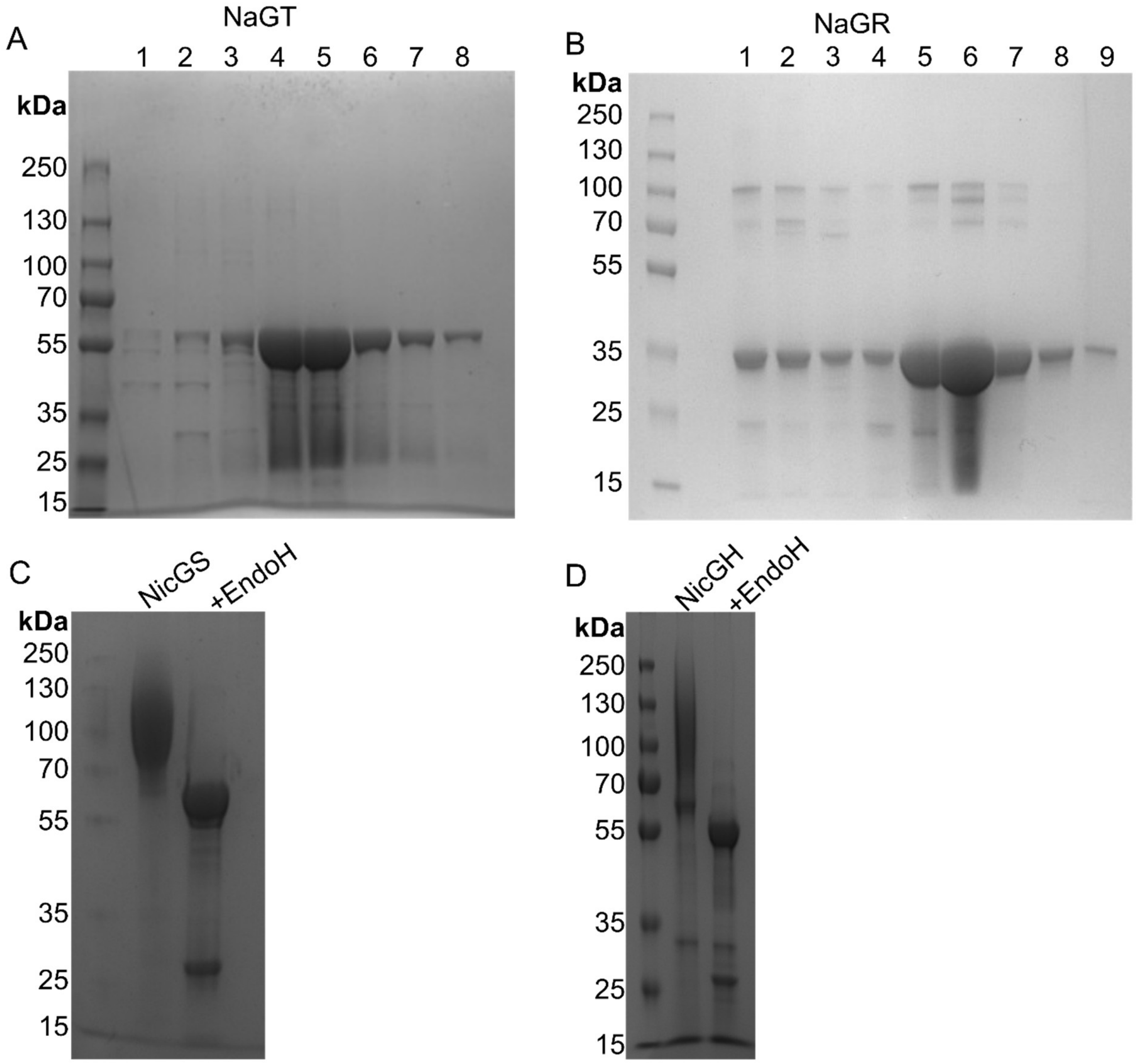


**Figure S2. SDS-Page gels showing the heterologously produced *N. tabacum* enzymes.** (**A**) NaGT (UGT1) expression and purification from *E. coli*. Gel shows size-exclusion column (SEC) output following a NiNTA column purification. Fractions represented by 4-8 were combined and used for enzyme assays. (**B**) NaGR (A622) expression and purification from *E. coli*. Gel shows size-exclusion column (SEC) output following a NiNTA column and anion exchange purification. Fractions represented by 5-9 were combined and used for X-ray crystallography and enzyme assays. (**C**) NicGS (BBLa) expression and purification from *K. phaffii*; gel shows validation of protein identity through deglycosylation with EndoH. (**D**) NicGH (β-GD1) produced in *K. phaffii*; gel shows validation of protein identity through deglycosylation with EndoH.

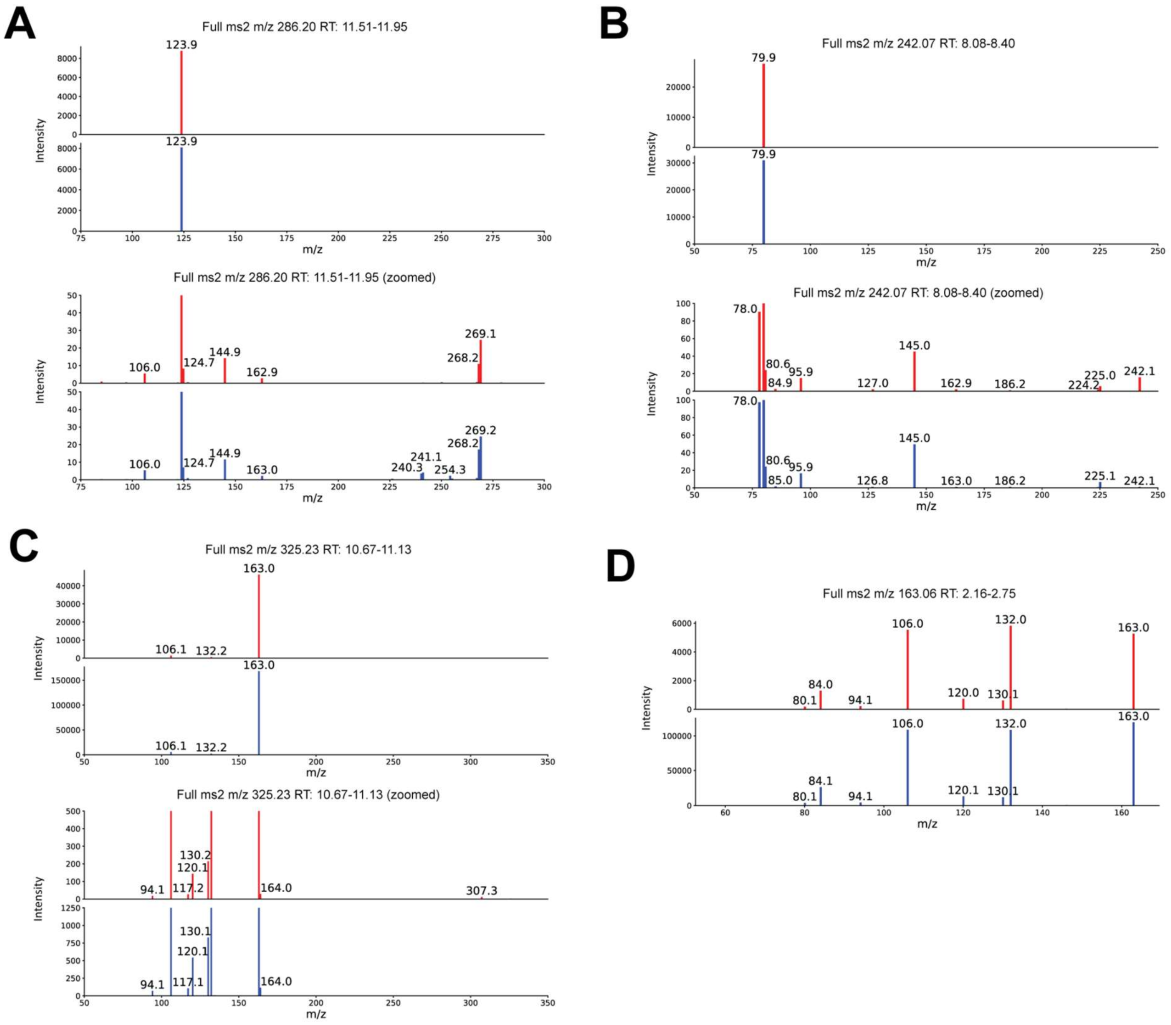


**Figure S3. MS$^2$ validation of peak identity in *in vitro* reactions.** MS$^2$ fragmentation spectra corresponding to Fig. 2, comparing chemically verified standards (blue spectra) to products from enzyme assays (red spectra). (**A**) Nicotinic acid *N*-glucoside **10.** (**B**) Pyridine glucoside **12.** (**C**) Nicotine glucoside (**13**)**.** (**D**) nicotine (**1**).

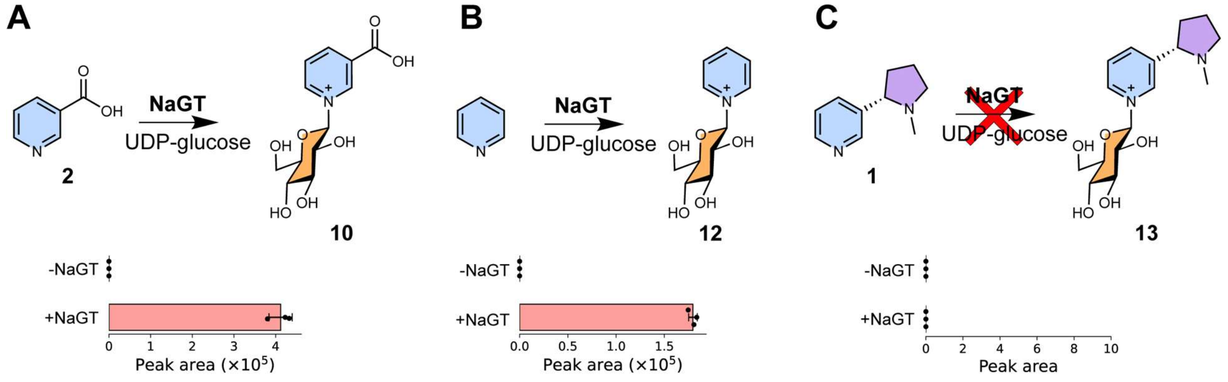


**Figure S4. NaGT substrate scope.** *In vitro* reactions using purified NaGT (UGT1) and UDP-glucose with different substrates. Control (-NaGT) was without NaGT enzyme but with UDP-glucose present. Bars show the peak area of EICs ($m/z$±0.15) corresponding to the masses of the chemical depicted above; error bars indicate standard error. (**A**) Nicotinic acid (**2**) substrate, showing enzymatic formation of nicotinic acid *N*-Glc (**10**). (**B**) Pyridine as substrate, showing enzymatic formation of pyridine *N*-Glc (**12**). (**C**) Nicotine ((*S*)-**1**) as substrate, with no product observed.

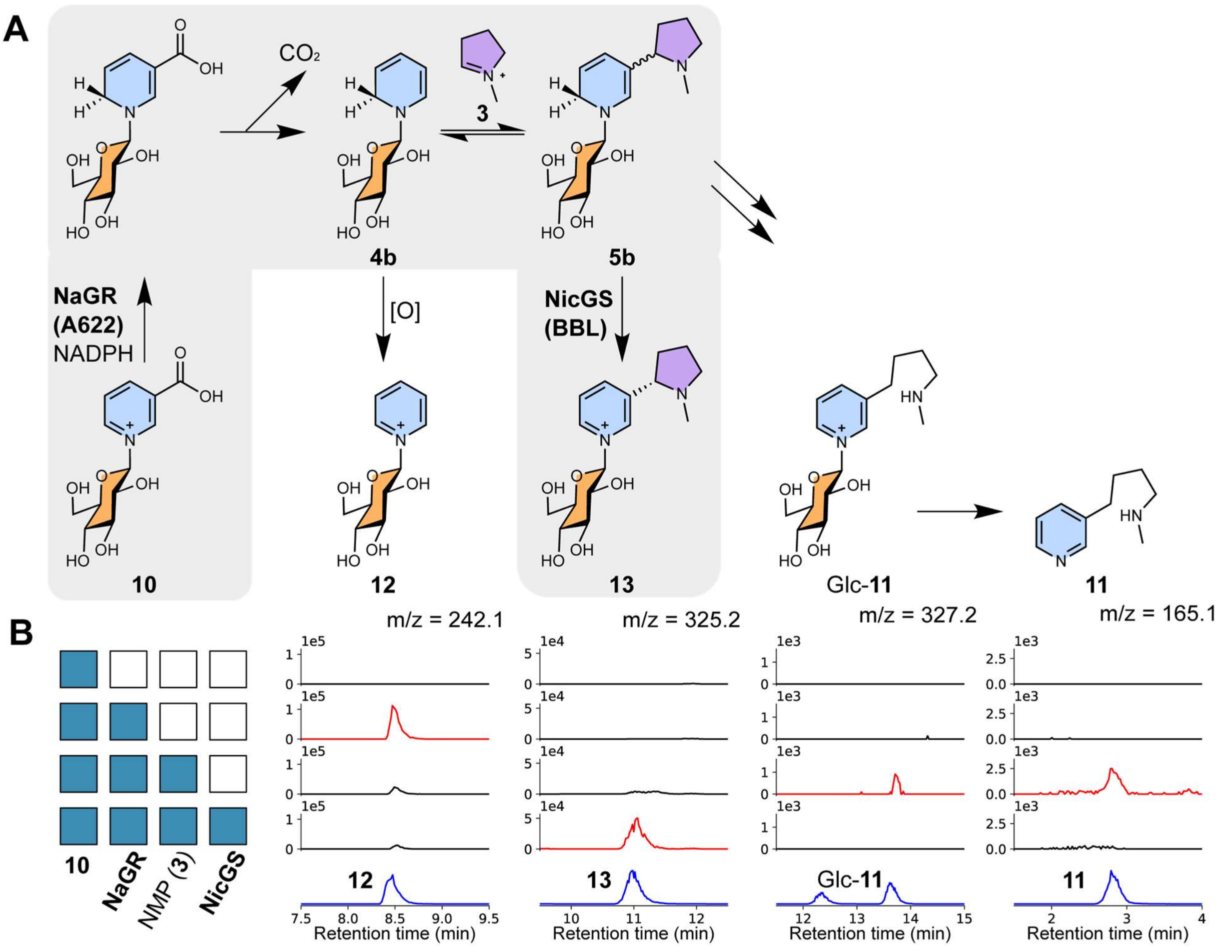


**Figure S5. Activities of NaGR (A622) and NicGS (BBLa).** (**A**) Proposed chemical transformations. (**B**) Products of *in vitro* reactions. Each row of chromatograms corresponds to an *in vitro* reaction, with the matrix showing presence/absence (blue/white) of reaction components, alongside NADPH. Each column of chromatograms shows EIC intensity (*m/z*±0.15) corresponding to the chemical depicted above (left-to-right: **12**, **13**, Glc-**11**, **11**). Compound identity was validated by chemically verified standards (blue). The Glc-**11** standard is present as both the α and β anomers. Signal intensity (*y*-axes) are comparable within each column, except for the standards which were scaled for clarity. Red chromatogram traces highlight key products formed as additional reaction components were included. The low intensity of Glc-**11** and **11** indicates trace formation.

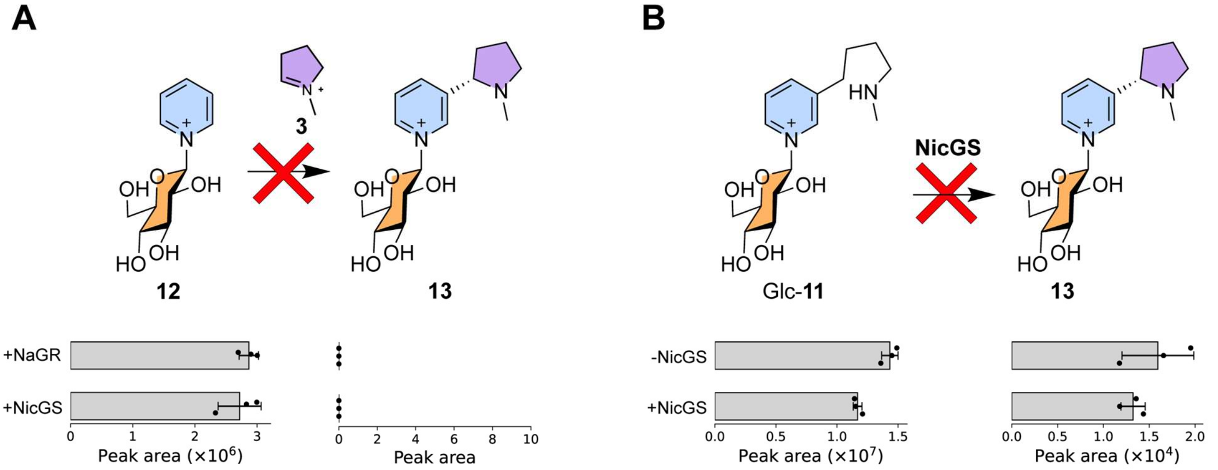


**Figure S6. Shunt products. (A)** Lack of activity of NaGR (with NADPH) or NicGS on pyridine glucoside (**12**) and **3**. **(B)** Glc-**11** incubated without and with NicGS showing no product formation. Low quantities of **13** are present in the standard for Glc-**11** (note the difference in peak area scaling)**.**

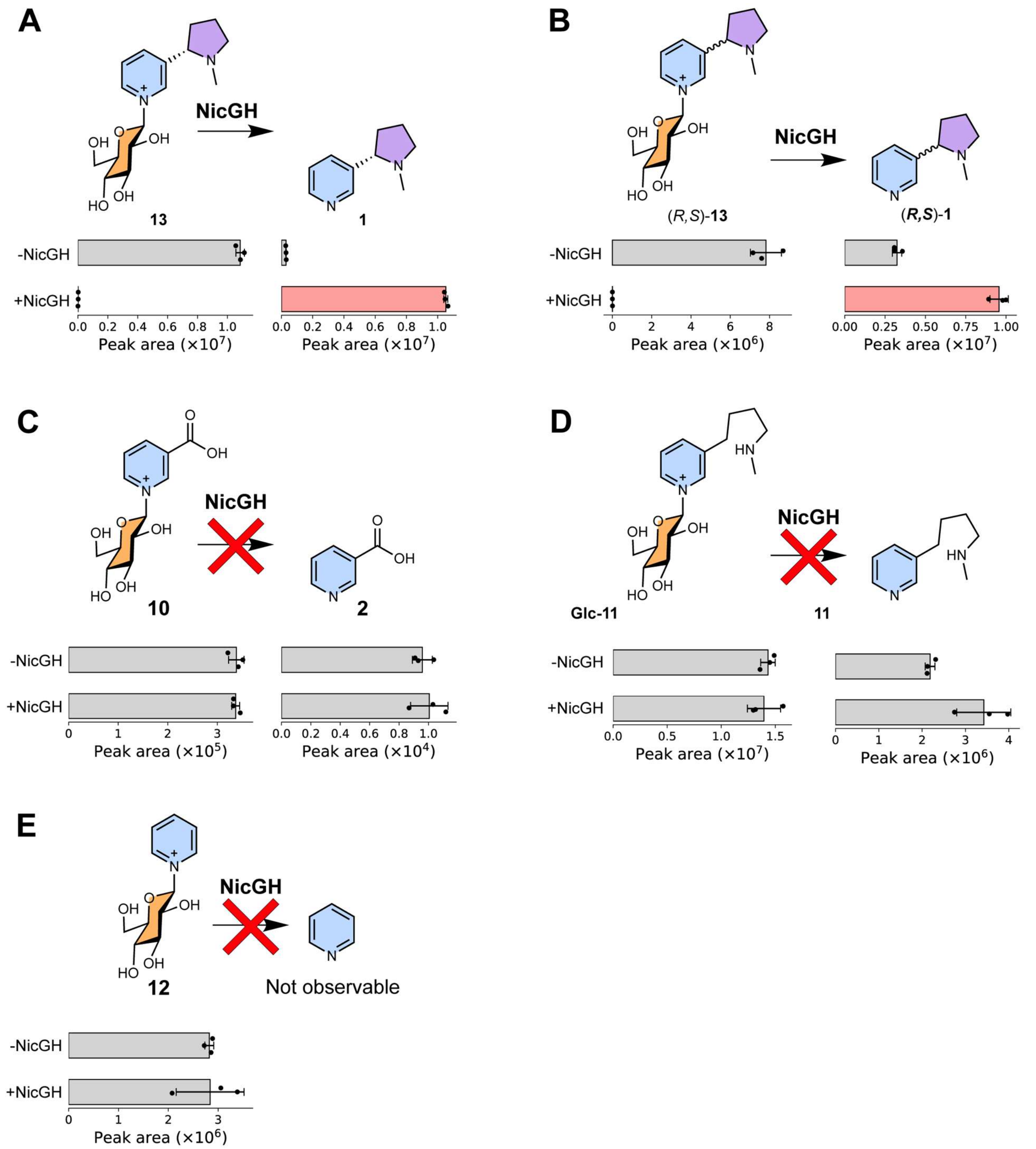


**Figure S7. *In vitro* activity of NicGH (*β*-GD1).** *In vitro* reactions using purified NicGH (*β*-GD1) with different substrates. (**A**) (*S*)-nicotine glucoside (*S*)-**13** as substrate, showing formation of nicotine (**1**). (**B**) (*R*,*S*)-nicotine glucoside ((*R*,*S*)-**13**) a substrate showing formation of nicotine (**1**). The lack of remaining substrate indicates both substrate diastereomers are consumed in the reaction. (**C**) Nicotinic acid N-glucoside (**10**) as substrate, showing no detectable product matching nicotinic acid (**2**) greater than the negative control. (**D**) Dihydrometanicotine glucoside (**Glc-11**) as substrate, showing no detectable consumption of substrate or formation of product above control. (**E**) Pyridine glucoside (**12**) as a substrate, showing no consumption of substrate.

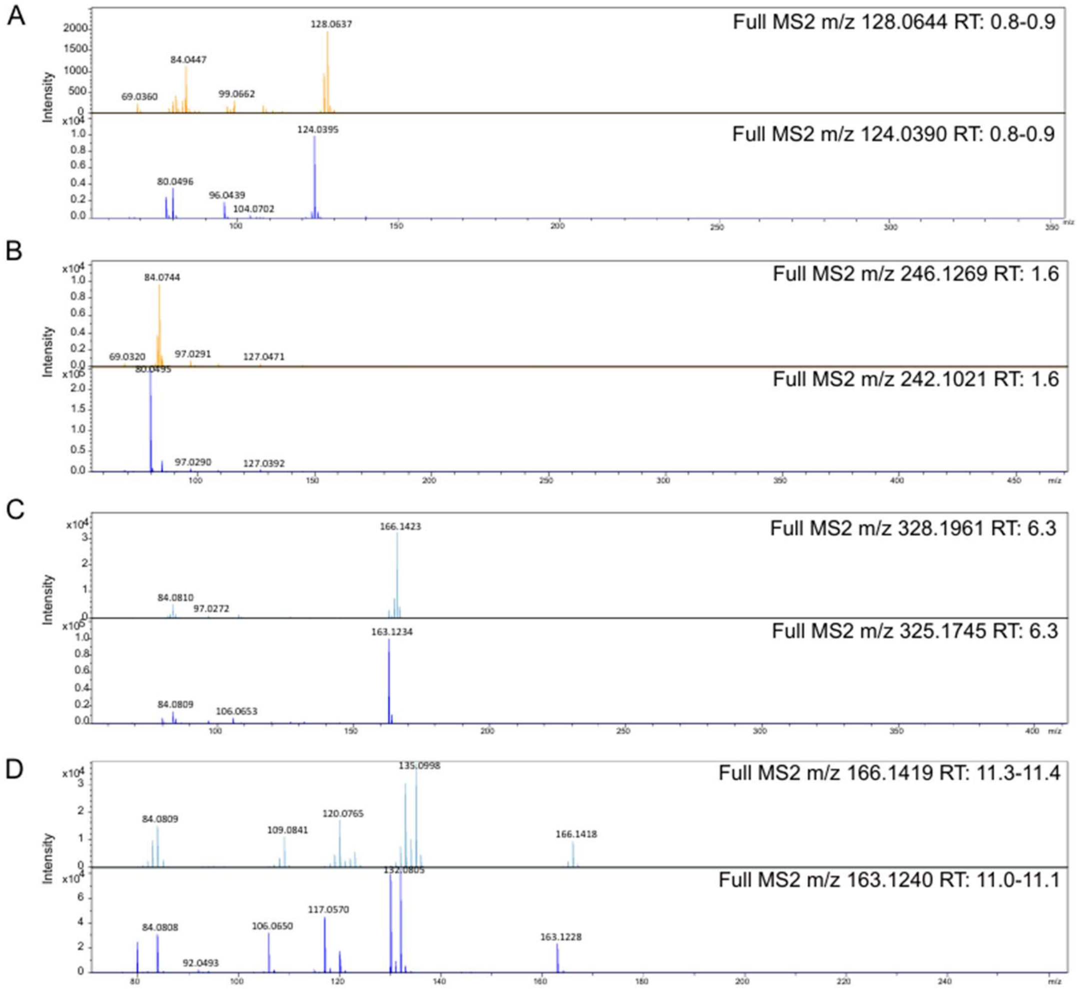


**Figure S8. MS² validation of peak identity in the *in planta* reconstitution.** Data were extracted from Fig. 4 with chemically verified standard MS² (dark blue) alongside MS² peaks from $d_3$ (light blue) or $d_4$ (orange) labelled compounds. (**A**) Nicotinic acid *N*-glucoside-$d_4$ **10**-$d_4$. (**B**) Pyridine glucoside-$d_4$ **12**-$d_4$. (**C**) Nicotine glucoside-$d_3$ **13**-$d_3$. (**D**) Nicotine-$d_3$ **1**-$d_3$

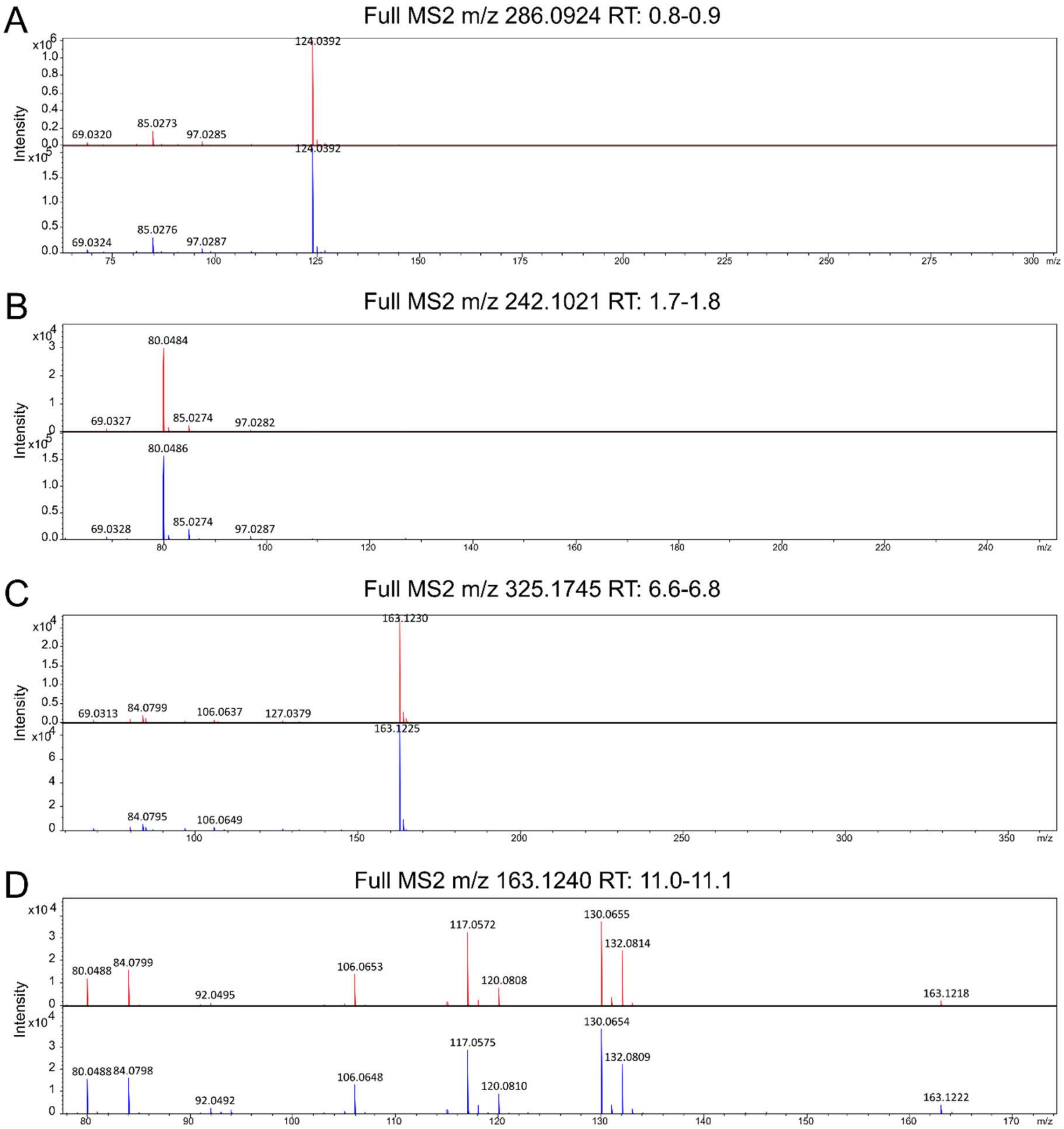


**Figure S9. MS$^2$ validation for metabolite identification in *N. benthamiana* roots.** Fragmentation data is derived from standards (blue) and samples (red) shown in Fig. 5A. (**A**) Nicotinic acid *N*-glucoside **10**. (**B**) Pyridne glucoside **12**. (**C**) Nicotine glucoside **13**. (**D**) Nicotine **1**.

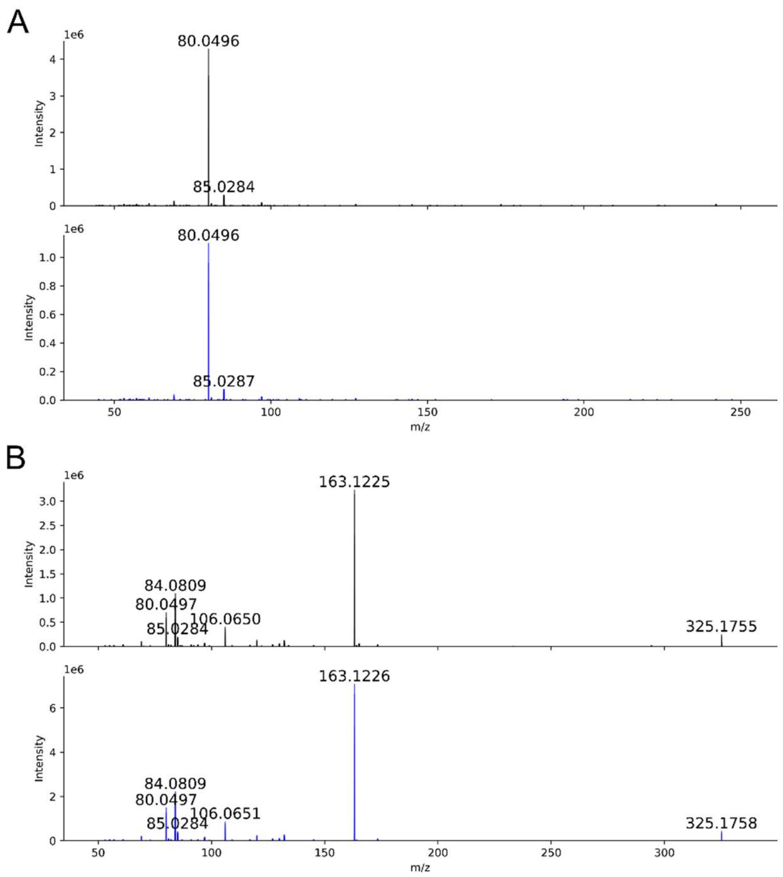


**Figure S10. MS² validation for metabolite identification in *N. tabacum* roots.** Fragmentation data is derived from standards (blue) and samples (black) shown in Fig. 5B. (**A**) Pyridine glucoside **12.** (**B**) Nicotine glucoside **13.**

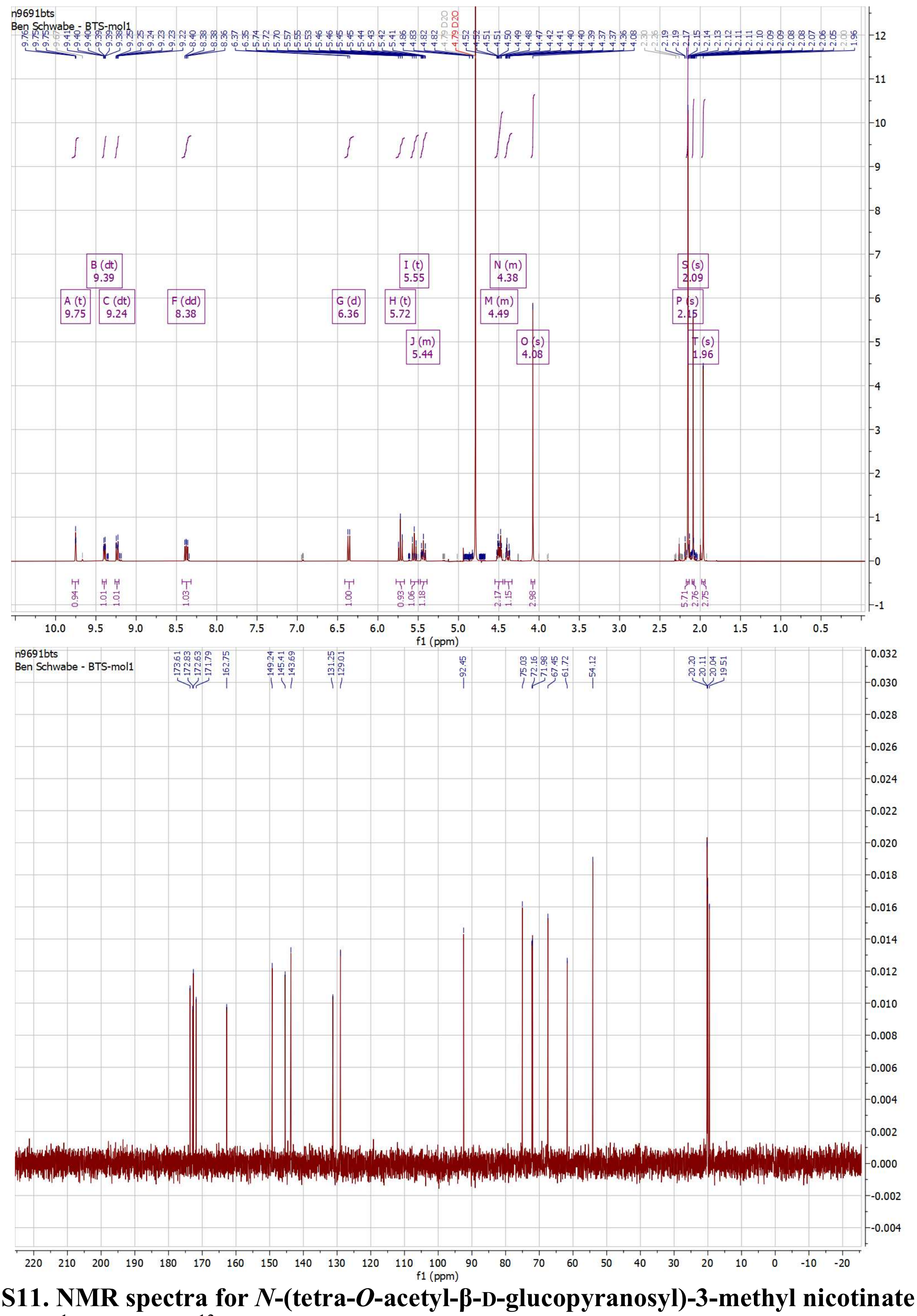

**Fig. S11. NMR spectra for *N*-(tetra-*O*-acetyl-β-D-glucopyranosyl)-3-methyl nicotinate bromide.** $^{1}$H (top) and $^{13}$C (bottom).

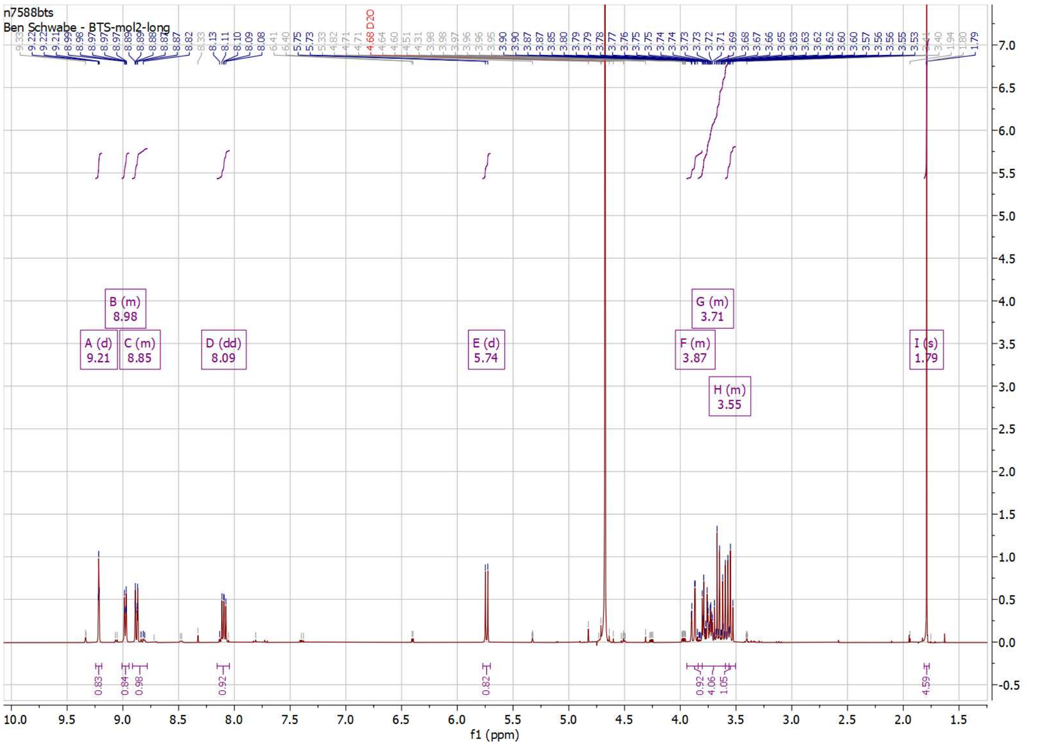


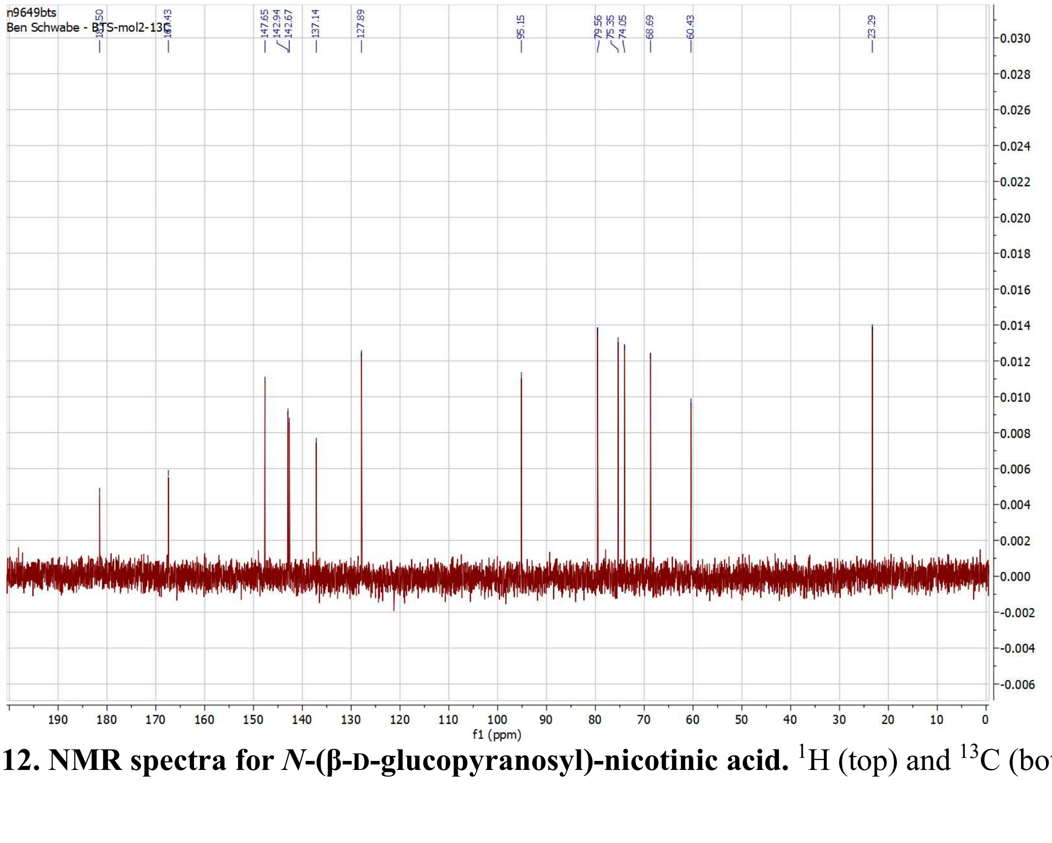


**Fig. S12. NMR spectra for *N*-(β-D-glucopyranosyl)-nicotinic acid.** $^{1}$H (top) and $^{13}$C (bottom).

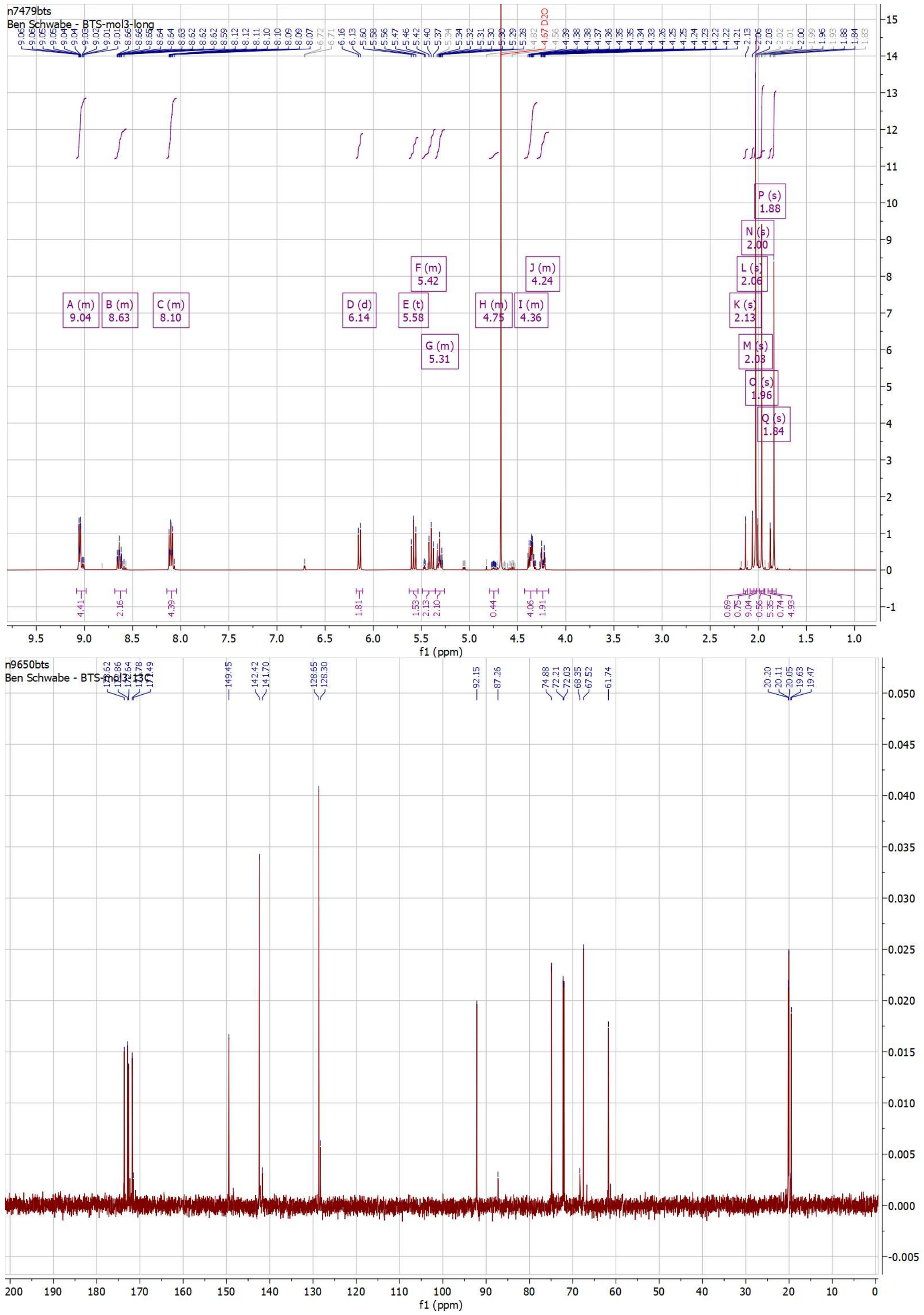


**Fig S13. NMR spectra for *N*-(tetra-*O*-acetyl-β-D-glucopyranosyl)-pyridinium bromide.** $^{1}$H (top) and $^{13}$C (bottom).

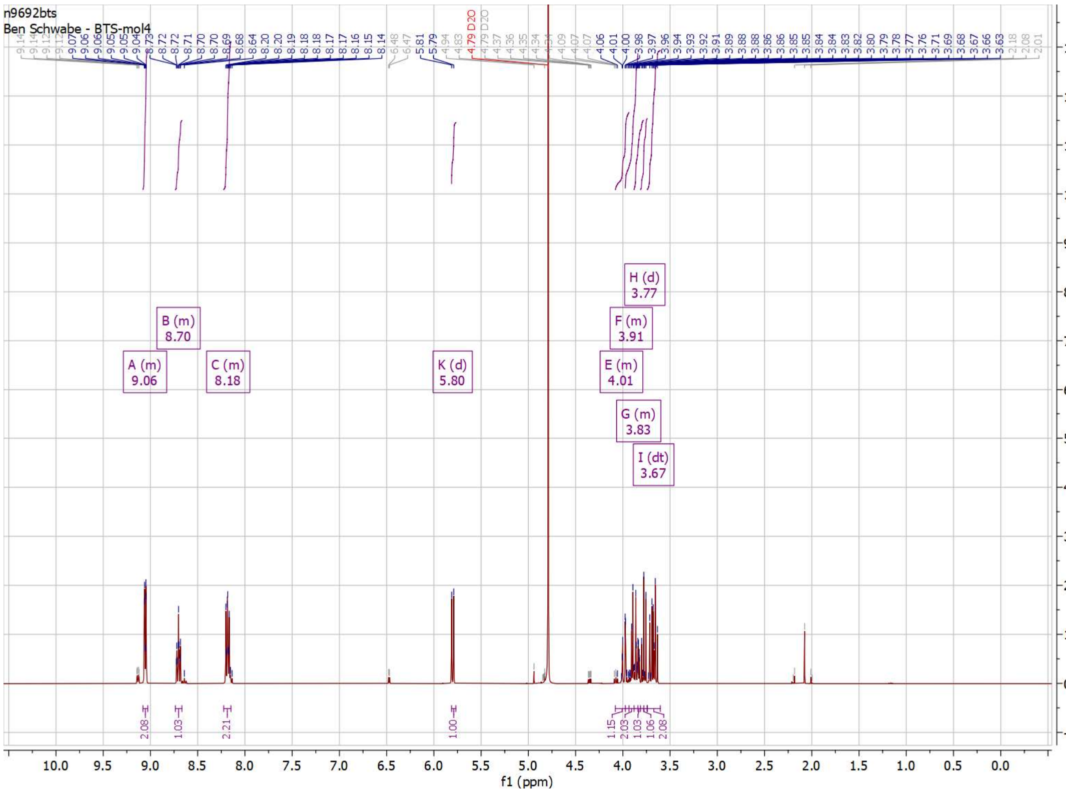


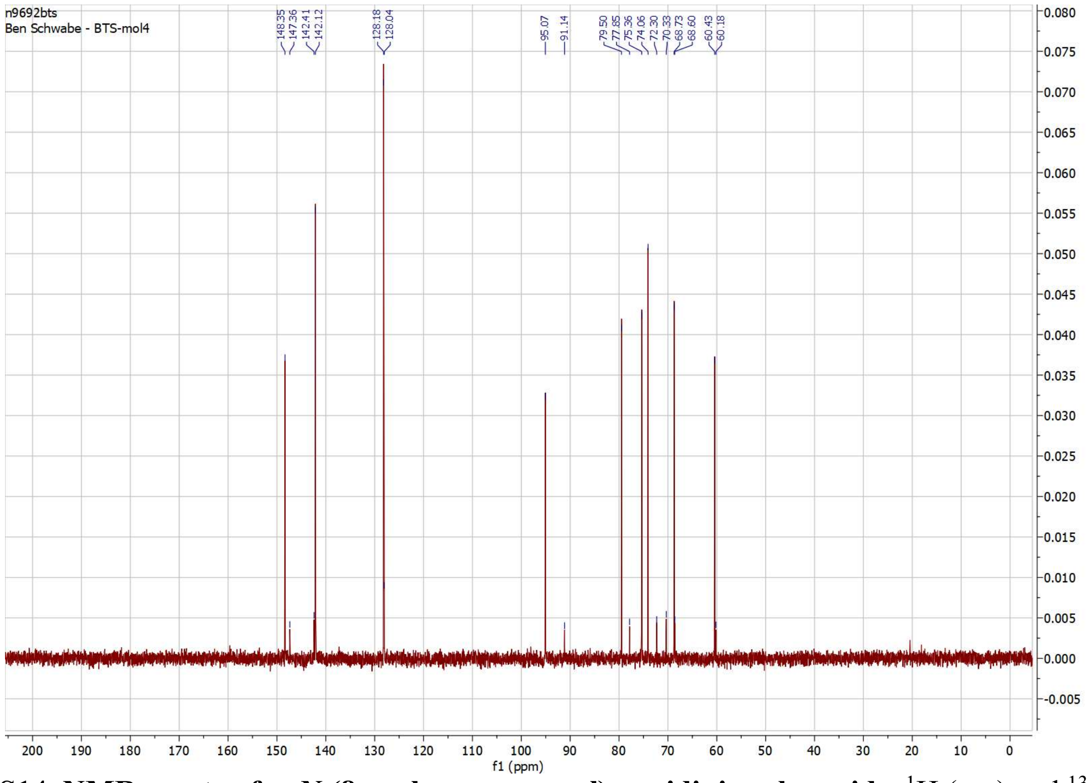


**Fig S14. NMR spectra for *N*-(β-D-glucopyranosyl)-pyridinium bromide.** $^{1}$H (top) and $^{13}$C (bottom).

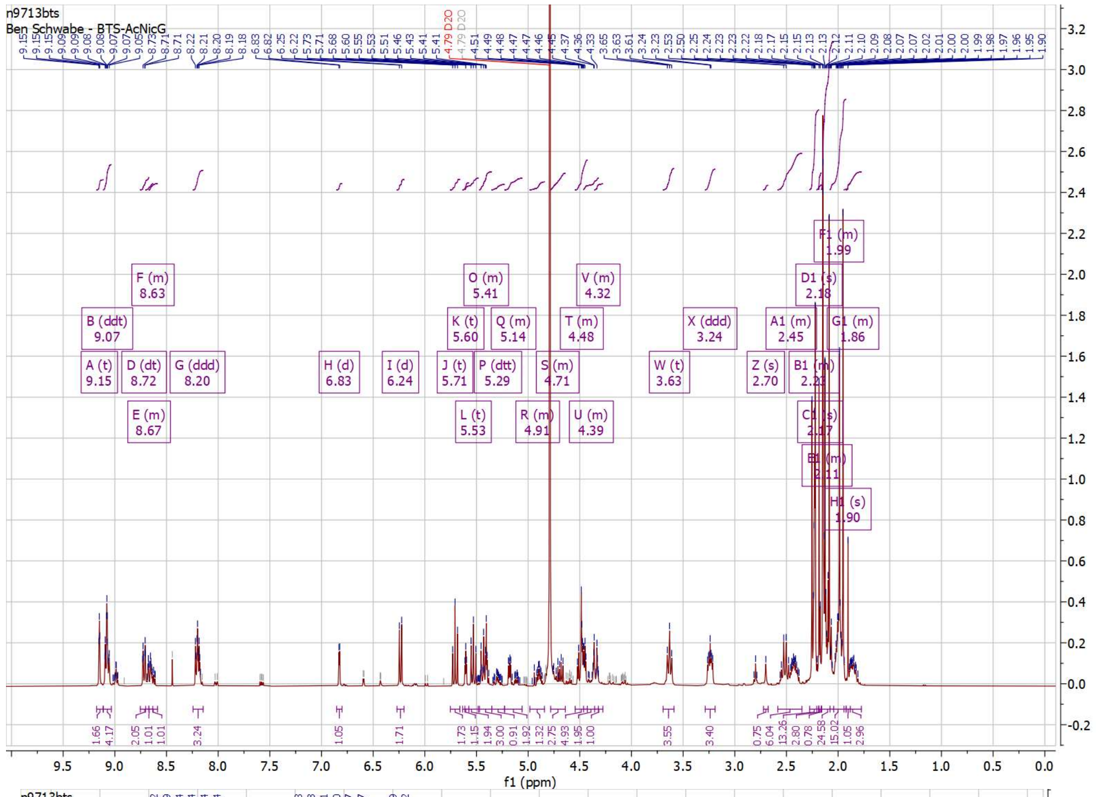


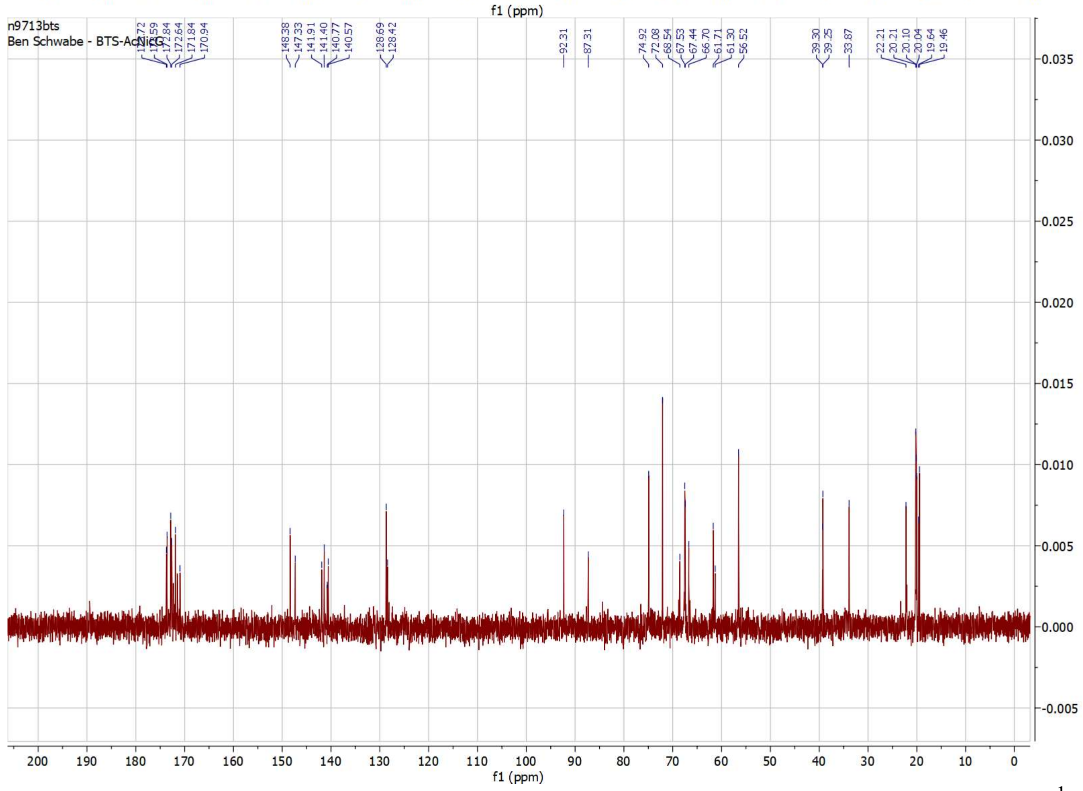


**Fig S15. NMR spectra for *N*-(tetra-*O*-acetyl-β-D-glucopyranosyl)-nicotine bromide.** $^{1}$H (top) and $^{13}$C (bottom).

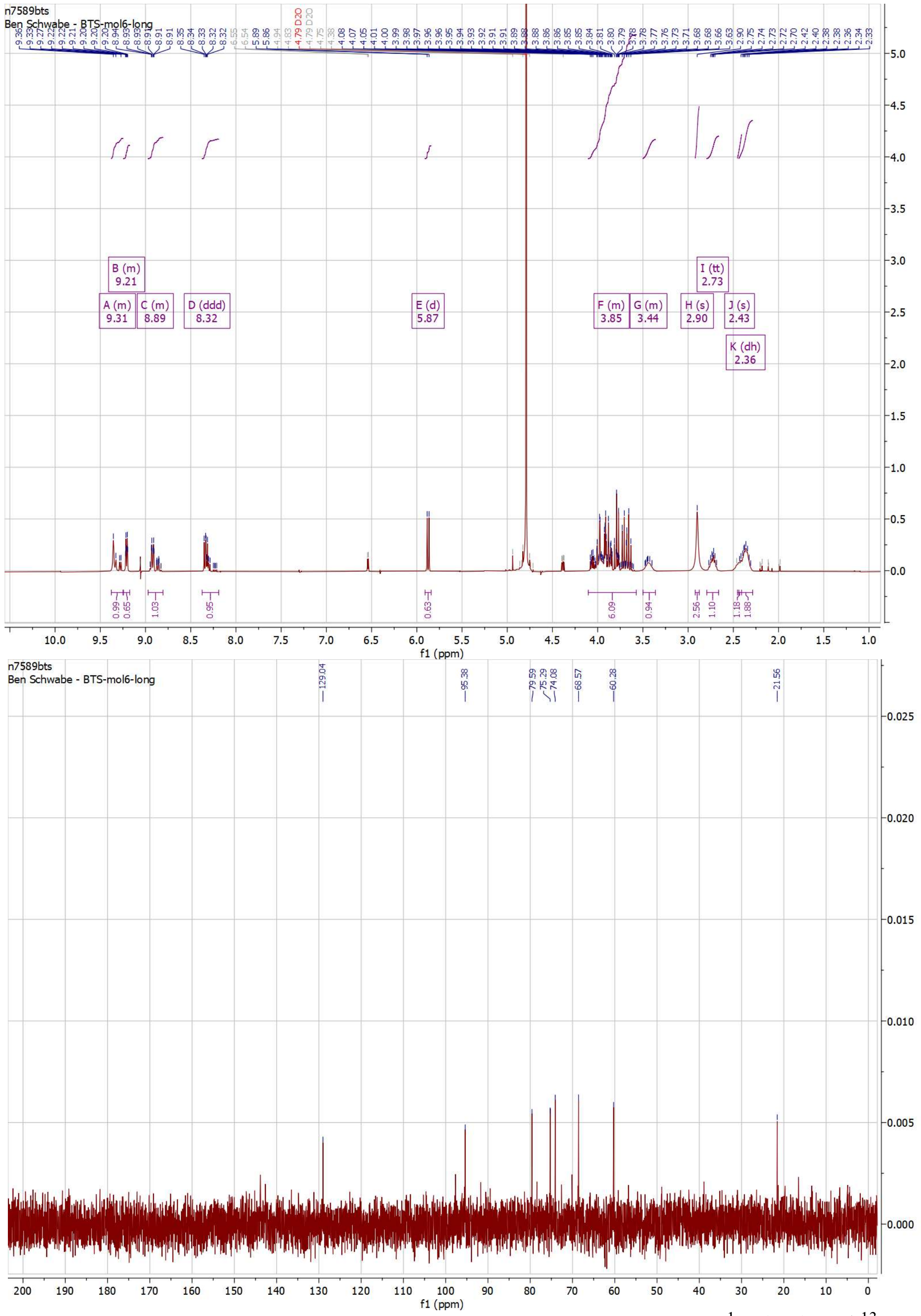


**Fig S16. NMR spectra for *N*-(β-D-glucopyranosyl)-nicotine bromide.** $^{1}$H (top) and $^{13}$C (bottom).

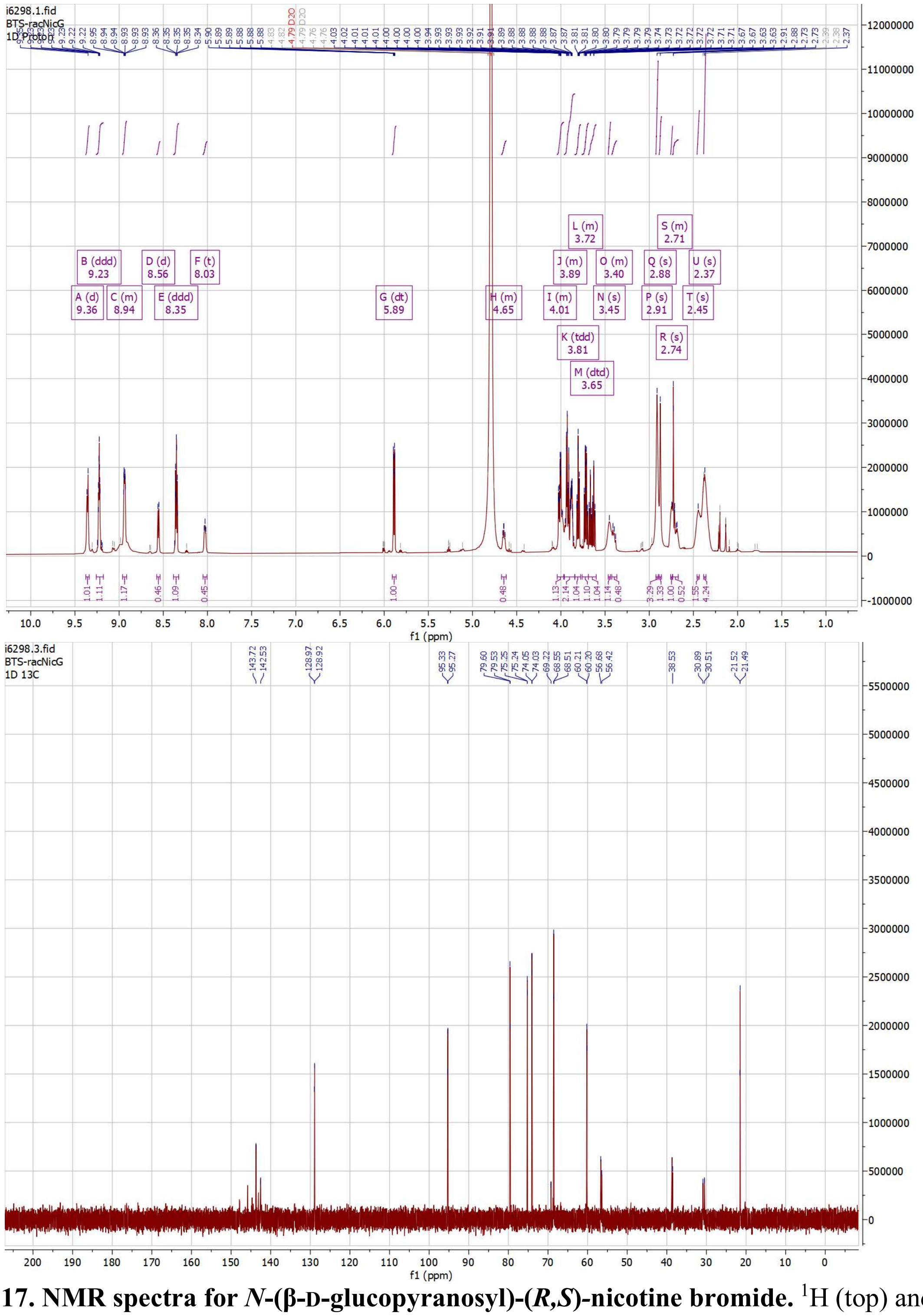


**Fig. S17. NMR spectra for *N*-(β-D-glucopyranosyl)-(*R,S*)-nicotine bromide.** $^{1}H$ (top) and $^{13}C$ (bottom).

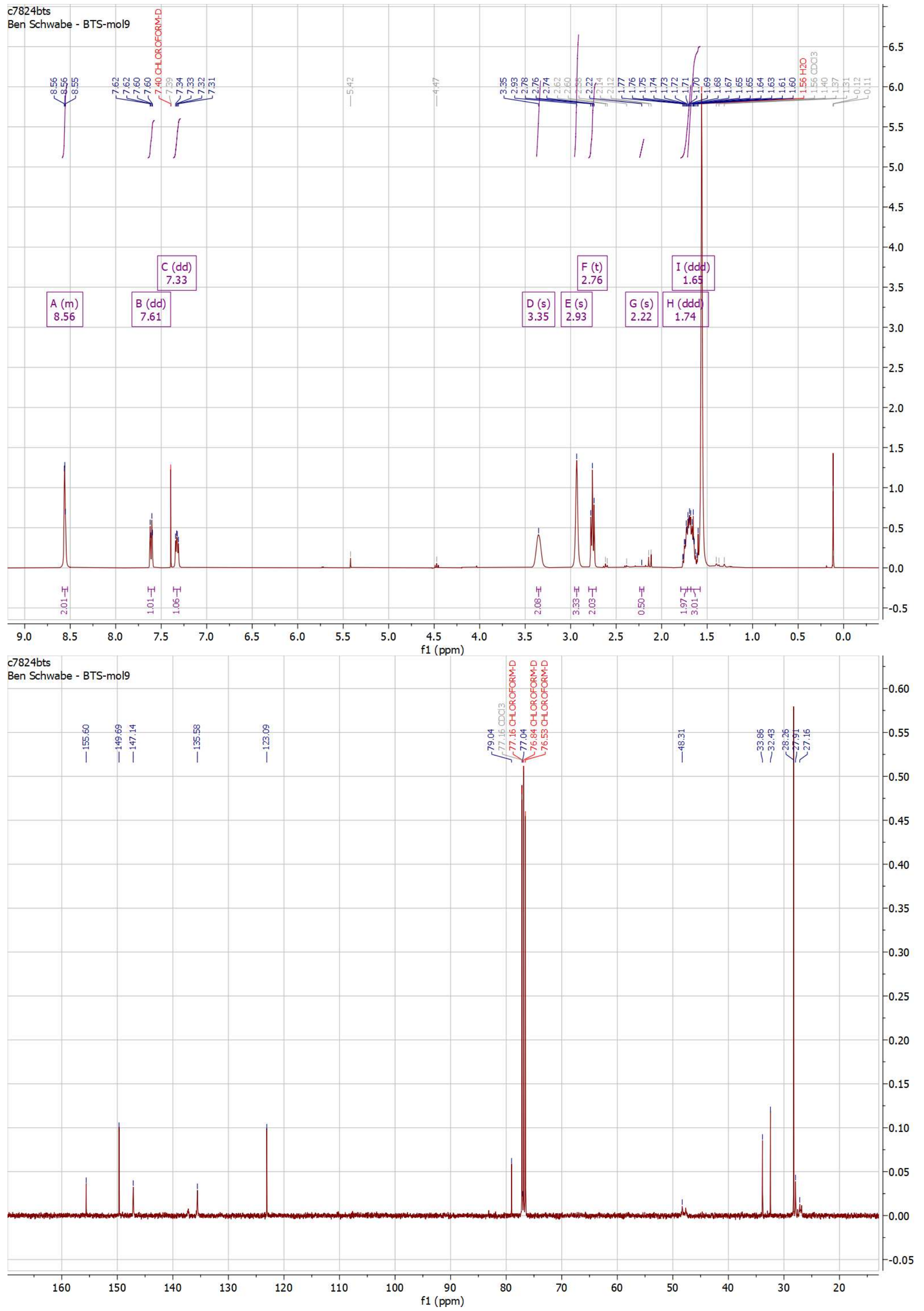


**Fig S18. NMR spectra for *N'*-boc-dihydrometanicotine.** $^{1}$H (top) and $^{13}$C (bottom).

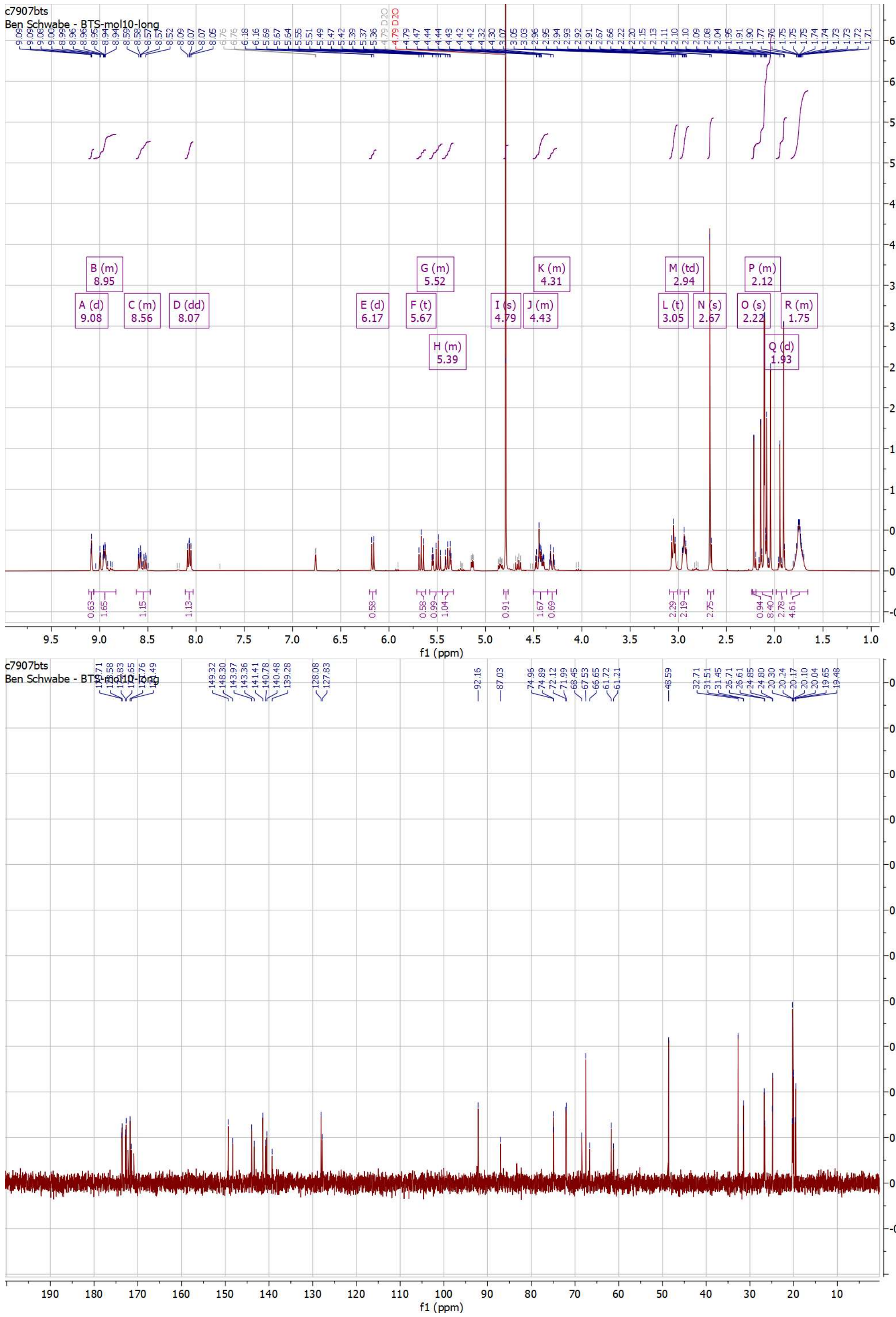


**Fig S19. NMR spectra for *N*-(tetra-*O*-acetyl-β-D-glucopyranosyl)-dihydrometanicotine bromide.** $^{1}H$ (top) and $^{13}C$ (bottom).

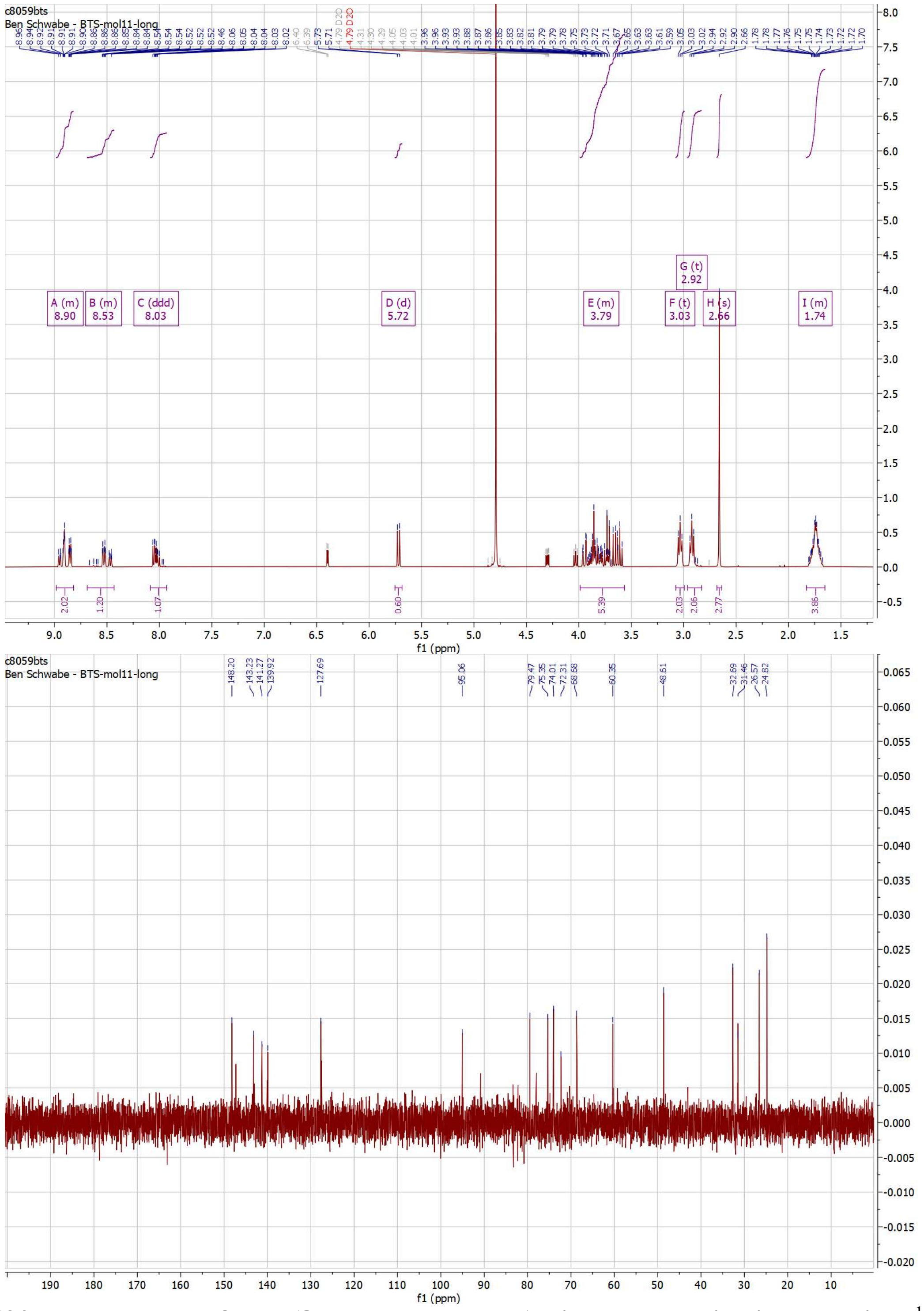


**Fig. S20. NMR spectra for *N*-(β-D-glucopyranosyl)-dihydrometanicotine bromide.** $^{1}$H (top) and $^{13}$C (bottom).
